# K-FIT: An accelerated kinetic parameterization algorithm using steady-state fluxomic data

**DOI:** 10.1101/612994

**Authors:** Saratram Gopalakrishnan, Satyakam Dash, Costas Maranas

## Abstract

Kinetic models predict the metabolic flows by directly linking metabolite concentrations and enzyme levels to reaction fluxes. Robust parameterization of organism-level kinetic models that faithfully reproduce the effect of different genetic or environmental perturbations remains an open challenge due to the intractability of existing algorithms. This paper introduces K-FIT, an accelerated kinetic parameterization workflow that leverages a novel decomposition approach to identify steady-state fluxes in response to genetic perturbations followed by a gradient-based update of kinetic parameters until predictions simultaneously agree with the fluxomic data in all perturbed metabolic networks. The applicability of K-FIT to large-scale models is demonstrated by parameterizing an expanded kinetic model for *E. coli* (307 reactions and 258 metabolites) using fluxomic data from six mutants. The achieved thousand-fold speed-up afforded by K-FIT over meta-heuristic approaches is transformational enabling follow-up robustness of inference analyses and optimal design of experiments to inform metabolic engineering strategies.

## Introduction and Background

The pressing need for the rapid development of truly predictive models of metabolism to accelerate build-design-test cycles for metabolic engineering has been widely reported (Cheng and Alper, 2014; Dromms and Styczynski, 2012; Long et al., 2015). Advances in synthetic biology (Chae et al., 2017; Cho et al., 2018; Stovicek et al., 2017) have alleviated the challenge of genome editing, placing the onus on the identification of suitable genetic modifications in metabolic engineering projects. Kinetic models of metabolism can alleviate the limitations of stoichiometric strain design techniques by quantitatively describing the relationship between fluxes, enzyme levels, and metabolite concentrations based on mechanistic and/or approximate rate law formalisms. The promise of superior product yield and production rate prediction offered by kinetic models comes at the expense of substantially increased experimental data requirements and complexity in model assembly, parameterization and interpretation of results. Long parameterization times stemming from poor scalability of existing parameterization frameworks ultimately preclude the deployment of follow up statistical analyses for large-scale kinetic models limiting insights into (i) the robustness of resolution of kinetic parameters given mutant flux datasets, (ii) kinetic parameter confidence levels, and (iii) the need for follow up measurements to improve prediction. These challenges motivate the development of K-FIT, a decomposition-based approach for parameterization of kinetic models using steady-state fluxomic and/or metabolomic data collected for multiple perturbation mutants. K-FIT builds upon the concept of Ensemble Modeling (EM) by anchoring concentrations and kinetic parameters to a reference strain. Unlike earlier efforts that employed a genetic algorithm (Khodayari et al., 2014) to parameterize kinetic models, K-FIT achieves three orders of magnitude improvement in efficiency by relying on a customized decomposition approach to compute steady-state fluxes in mutant networks. K-FIT was first benchmarked against EM for three test kinetic models of increasing size ranging from 100 to 953 kinetic parameters to demonstrate the increase in computational savings with model size. K-FIT remained tractable even for a near genome-scale kinetic model containing 307 reactions, 258 metabolites, and 2,367 kinetic parameters parameterized with 1,728 steady-state fluxes from six single gene-deletion mutants determined using 13C-metabolic flux analysis (13C-MFA) (Long et al., 2018). The parameterization was carried out 100 times with random initializations and was completed within 48 hours of cumulative computation time. The best solution was recovered 44 out of 100 times providing confidence that convergence to the true optimum was indeed achieved. The kinetic model k-ecoli307 accurately recapitulated fluxes to within 15 mmol/gdw-h of the fluxes reported by 13C-MFA while also predicting fluctuations in glucose uptake in response to genetic perturbation and flux rerouting through energy metabolism to meet biosynthetic NADPH demands. The yield predictions of acetate, lactate, and malate for engineered strains were found to be within 30% of the experimental yield for metabolites derived from central metabolism. K-FIT offers, with no additional effort, local kinetic parameter sensitivity information through the computation of gradients allowing for optimality testing of all obtained solutions. This gradient output also enables the straightforward computation of parametric uncertainty, establishment of confidence intervals for kinetic parameters and derivation of metabolic control coefficients. This post-parameterization information can aid in the interpretation of kinetic model predictions, suggest efficient experiment design to improve prediction fidelity and ultimately inform metabolic engineering strategies.

## Results

In this section, first a schematic representation of the workflow of K-FIT is described. The performance of K-FIT is benchmarked against the Ensemble Modeling (EM) (Khodayari et al., 2014) using three test kinetic models to assess the impact of model scale-up on the computational savings afforded by K-FIT. The applicability of K-FIT to near genome-scale models is then demonstrated by parameterizing an expanded kinetic model for *E. coli* (k-ecoli307) containing 307 reactions, 258 metabolites, and 2,367 kinetic parameters parameterized using 13C-labeling data for ten proteinogenic amino acids from six single gene-deletion mutants (Long et al., 2018). Local regression based standard deviations for all estimated elementary kinetic and Michaelis-Menten parameters are computed by leveraging the gradient calculations embedded within K-FIT. The predictive capability of k-ecoli307 is then assessed by comparing predicted product yields against experimentally measured yields for six over-producing strains not used during model parameterization.

### The K-FIT Algorithm

K-FIT is a gradient-based kinetic parameter estimation algorithm using steady-state flux measurements from multiple genetic perturbation mutants. The schematic workflow for K-FIT is shown in Figure 1 and Supplementary Figure S1. Reaction fluxes are related to metabolite concentrations using mass-action kinetics after decomposition of the enzyme catalytic mechanism into elementary steps (see supplementary methods for the detailed procedure for elementary step decomposition) as described by Tran *et al* (Tran et al., 2008). Conservation of mass across enzyme complexes and metabolites is therefore expressed as a system of bilinear equations. This formalism was chosen because it is mechanistically sound, obeys mass conservation laws and is inherently thermodynamically feasible (Saa and Nielsen, 2017). At the same time, it allows for easy integration of allosteric regulation without the need to derive cumbersome case-specific nonlinear rate laws. The K-FIT algorithm iteratively applies the following three steps till the optimal set of kinetic parameters is found: (i) K-SOLVE, (ii) Steady-State Flux Evaluator (SSF-Evaluator) and (iii) K-UPDATE.

**Figure 1:**
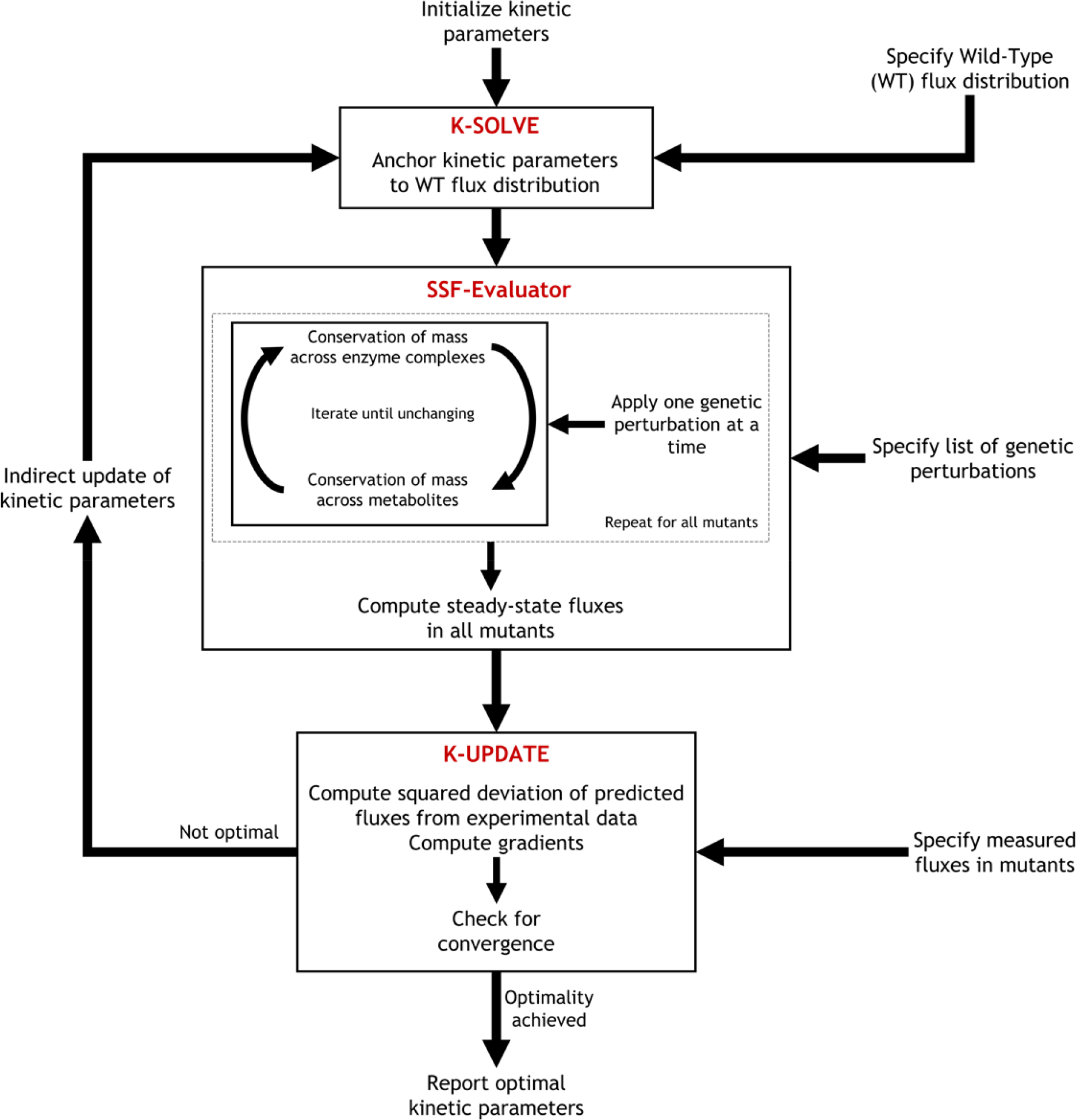
Overview of the core loop of the K-FIT algorithm

The objective of K-SOLVE is to anchor kinetic parameters to a reference state which is typically the wild-type (WT) network similar to the procedure described by Tran *et al.* (Tran et al., 2008). This is needed because for arbitrary assignment of values to the elementary kinetic parameters enzyme and metabolite conservation balances are not always satisfied. K-SOLVE uses as input the WT reverse fluxes of all elementary steps and the WT enzyme fractions. WT net fluxes are treated differently than mutant net fluxes. While net fluxes from the mutant networks form the sum of squares objective function to be minimized, net fluxes in the WT network are fixed in the model to their experimentally resolved values. This ensures that the inferred kinetic parameters remain consistent with both experimental WT flux measurements and inherently satisfy mass conservation laws for both enzymes and metabolites. Consequently, the specified (i) reverse WT elementary fluxes and (ii) WT enzyme fractions along with (iii) the specified net WT fluxes fix all degrees of freedom in the system of equations. Unique values for all kinetic parameters can thus be assigned so as they inherently satisfy mass balances across metabolites and enzymes in the WT network. This is an important consideration as mass balances are not always satisfied under metabolic steady-state for an arbitrary assignment of values to the kinetic parameters.

Kinetic parameters anchored by K-SOLVE are then used by SSF-Evaluator to compute the steady-state fluxes and concentrations across all mutants, one at a time. The system of bilinear algebraic equations in metabolite and enzyme fractions resulting from the elementary step decomposition of enzyme catalysis is partitioned into two sub-problems. The first bilinear sub-problem, representing conservation of mass across enzyme complexes, is reduced to a system of linear algebraic equations in enzyme fractions in the mutant network when the metabolite concentrations in the mutant network are specified. Similarly, the second bilinear sub-problem describing conservation of mass across all metabolites reduces to a system of linear algebraic equations in metabolite concentrations in the mutant network when the enzyme fractions in the mutant network are specified. By iterating between these two linear sub-problems, the steady-state enzyme levels and metabolite concentrations in each mutant is identified. SSF-Evaluator begins with fixed-point iterations and switches to Newton steps closer to the solution. A semi-implicit Euler integration step is carried out whenever the Jacobian matrix in the Newton step becomes singular. Convergence is achieved when the concentrations of enzyme complexes and metabolites remain almost unchanged between successive iterations. This iterative scheme therefore enables the direct evaluation of steady-state fluxes (almost) without the need to integrate any ODEs and contributes to the speed-up of the kinetic parameterization process.

The calculated fluxes for all mutants are compared against the corresponding measured fluxes and the sum of squared residuals (SSR) representing the variance-weighted squared deviation of predicted fluxes from experimental data, as well as the first- and second-order gradients are computed by K-UPDATE. The WT reverse elementary fluxes and WT enzyme fractions are then updated using a Newton step and the core loop of K-FIT is repeated by returning to K-SOLVE until the minimum deviation of predicted fluxes from experimental measurements is reached. Note that the computation of first- and second-order gradients uses as input the local sensitivity of fluxes with respect to kinetic parameters. These local sensitivities can readily be computed by solving a system of linear equations (see supplementary methods) as opposed to having to perform costly forward sensitivity analysis. This enables K-FIT to confirm that any reported solution is indeed optimal while also allowing for the assembly of the covariance matrix from which approximate confidence intervals for all estimated kinetic parameters can efficiently be derived.

### Benchmarking K-FIT against Ensemble Modeling

The computational performance of the K-FIT algorithm was first compared against solution with a genetic algorithm (GA) operating on a population of models constructed using the Ensemble Modeling (EM) approach. Three test models of increasing sizes were used to assess the impact of model size on parameter estimation speed and solution reproducibility. The first small model containing 14 reactions, 11 metabolites, and 100 kinetic parameters was adapted from the three glycolytic pathways in *E. coli* and was parameterized using flux distributions from four single gene-deletion mutants (Supplementary Fig. S2a). The second medium-sized kinetic model containing 33 reactions, 28 metabolites, and 235 kinetic parameters was adapted from a previous study (Greene et al., 2017) and was parameterized using flux distributions from seven single gene-deletion mutants (Supplementary Fig. S2b). The third test model describing carbon flows through central and amino acid metabolism was adapted from the model developed by Foster *et. al.*, (2019 (Under Review)). This model (Supplementary Fig. S2c) contains 108 reactions, 65 metabolites, and 953 kinetic parameters and was parameterized using flux distributions from seven single gene-deletion mutants. The test models along with the data required for kinetic parameterization is provided in supplementary tables ST1 – ST12.

Kinetic parameters were estimated in 9 minutes, 30 minutes, and 4 hours for the three models, respectively, using K-FIT. In contrast, GA required 60 hours, 726 hours, and 4,278 hours, respectively, to parameterize the same three models. Computational speed-up increased from 100-fold for the small model to 1000-fold for the core kinetic model upon switching from GA to K-FIT. This dramatic reduction in parameterization time arises from the (largely) integration-free steady-state flux evaluation using the SSF-Evaluator step and the fact that K-FIT traverses the variable space in a highly economical manner (i.e., Newton steps) requiring fewer than 500 steady-state flux evaluations to identify the optimal solution. In contrast, the GA approach relies on iterative recombination of kinetic parameter vectors and required as many as 20,000 steady-state flux evaluations before finding the same solution (Khodayari et al., 2014) but without confirming optimality. SSF-Evaluator evaluated steady-state fluxes, on average in 0.42 seconds, 1.13 seconds and 6.12 seconds, respectively, for the three models, whereas, numerical integration required 3.5 seconds, 120 seconds and 440 seconds, respectively. Bypassing integration also enables SSF-Evaluator to handle stiff systems of ODEs arising from the large dynamic range of kinetic parameters and ensuring that steady-state fluxes are always within a mass imbalance of just 0.001 mol%. It is important to note that Newton’s method can only guarantee convergence to a local and not necessarily the global minimum of SSR. As a safeguard against failure to reach the true minimum, models were parametrized using K-FIT from 100 random initial starting points to determine the reproducibility of the obtained best solution. For the three test models K-FIT exhibited a best solution recovery of 98%, 93%, and 60%, respectively. This high solution reproducibility provides confidence that K-FIT is able to consistently converge to the lowest SSR of 0 for the small model, 8.9 for the medium-sized model, and 1.3 for the core model, respectively. Notably, no alternate optima in the vicinity of the best solution (within an SSR of 100) was detected for any of the three models implying that the best SSR minimization solution is the only good kinetic parameterization candidate. The inherent ability of K-FIT to quickly calculate local sensitivities of predicted fluxes to metabolite concentrations was leveraged to confirm whether SSFEstimator reported steady-state concentrations that are stable. To this end, the eigenvalues of the Jacobian matrix (Greene et al., 2017) at metabolic steady-state were calculated to confirm that the real part of all eigenvalues were strictly negative for all iterations and problems solved.

### Parameterization of a kinetic model (k-ecoli307) for *E. coli* with near-genome-wide coverage

Following the application to three test models of increasing models, K-FIT was then deployed for the parameterization of k-ecoli307, an *E. coli* kinetic model with near-genome-wide coverage similar to k-ecoli457 (Khodayari and Maranas, 2016). The expanded model containing 307 reactions, 259 metabolites and 2,367 kinetic parameters, and encompasses central metabolism, expanded amino acid, fatty acid, and nucleotide pathways and lumped pathways for peptidoglycan biosynthesis. Compared to k-ecoli457 (Khodayari and Maranas, 2016), k-ecoli307 lacks the pathways for anaerobic metabolism and secretion of organic acids as it was parameterized using data under aerobic growth only. Flux data for six single gene-deletion mutants were computed using 13C-Metabolic Flux Analysis (13C-MFA) to recapitulate the measured labeling distribution of 10 proteinogenic amino acids grown with 1,2-^13^C-glucose as the carbon tracer (Long et al., 2018). This provided a total of 1,728 MFA-determined fluxes across six single gene-deletion mutants from upper glycolysis for kinetic parameterization. All 69 substrate level regulatory interactions for 26 reactions in the expanded model were transferred from k-ecoli457. Complete cofactor balances were not included in k-ecoli307. Instead, k-ecoli307 only accounts for the net moles of ATP, NADH and NADPH produced or consumed in any reaction and disregards the recycling of ADP, AMP, NAD and NADP back to the phosphorylated/reduced form of the cofactor. This simplification is necessary to avoid metabolite pool dependencies in the stoichiometry matrix. These dependencies force the total concentration of (ATP + ADP + AMP), (NAD + NADH), and (NADP + NADPH) to be held constant across all mutants (Vallabhajosyula et al., 2006) while allowing only the ratios of metabolite concentrations ATP/ADP, ADP/AMP, NAD/NADH and NADP/NADPH to vary across mutants (Greene et al., 2017).

The expanded model was parameterized using flux distributions from the six mutants Δ*pgi*, Δ*gnd*, Δ*zwf*, Δ*eda*, Δ*edd*, and Δ*fbp* with 288 fitted fluxes per mutant.100 fluxes were inferred using 13C labeling data whereas 207 reactions were growth-coupled. Parameterization using the K-FIT algorithm was completed in 48 hours on an Intel-i7 (4-core processor, 2.6GHz, 12GB RAM) computer with a minimum SSR of 131.05 and a solution reproducibility of 44%. The recapitulation of the experimentally measured fluxes by K-FIT for the six mutants is shown in Supplementary Figure S3. All predicted fluxes were within 15 mmol/gdw-h of their corresponding flux reported by 13C-MFA. This corresponds to a maximum deviation of only 10% from the experimentally determined fluxes. Flux distributions for Δ*eda*, Δ*edd*, and Δ*fbp* mutants were largely unchanged from WT (Supplementary Figures S3a, S3b, and S3c) alluding to the dispensability of the corresponding genes. In contrast, carbon flux was significantly rerouted in response to the knockout of *pgi*, *zwf*, and *gnd* genes. Glucose uptake remained similar to WT for the Δ*zwf* mutant but routed completely via the EMP pathway (Supplementary Figure S3d). The non-oxidative pentose phosphate pathway (TKT and TAL reactions) operated in reverse to generate ribose-5-phosphate for nucleotide biosynthesis. Glucose catabolism solely via the EMP pathway increased acetate and biomass yields by 10%. The expanded model also revealed that the loss of NADPH production via the oxidative pentose phosphate pathway was compensated by a 90% increase in the flux through the transhydrogenase reaction in the Δ*zwf* strain (Supplementary table ST20). Glucose uptake for the Δ*gnd* mutant was decreased by less than 10% compared to WT (Supplementary Figure S3e). Δgnd was the only strain with a measurable flux through the ED pathway by rerouting 24.9 mmol/gdw-h of flux through EDD and EDA reactions is a ten-fold increased flux through the ED pathway relative to the WT network. Similar to Δ*zwf*, the reversal of flux through the non-oxidative pentose phosphate pathway generated the required ribose-5-phosphate for nucleotide biosynthesis.

Of all the mutants, Δ*pgi* involved the most significant flux rerouting relative to WT. Glucose uptake was reduced by 75% compared to WT resulting in a 70% reduction in growth rate (Supplementary Figure S3f). Flux redirection through the glyoxylate shunt and reduction of acetate secretion improved carbon routing towards biomass precursors, thereby increasing the biomass yield by 22% compared to WT. Interestingly, the ED pathway was found to carry only 2 mmol/gdw-h of flux with almost all of the carbon being metabolized via the pentose phosphate pathway (Supplementary Figure S3f). In addition, an 80% reduction in flux through glycolysis along with the absence of acetate secretion lowers overall glycolytic ATP production. This loss is compensated by the reversal of flux through the transhydrogenase reaction relative to WT which allows the oxidation of excess NADPH generated by the oxidative pentose phosphate pathway (Supplementary Table ST20). k-ecoli307 captured that non-competitive inhibition of the EDA reaction by glyceraldehyde-3-phosphate limits flux through the ED pathway in all strains but Δ*gnd*. In the Δ*gnd* mutant, a 37-fold increase in the concentration of 6-phosphogluconate provided the necessary driving force to overcome this product inhibition, thereby allowing a flux of 24.9 mmol/gdw-h through the ED pathway. In contrast, in mutant Δ*pgi* a two-fold increase in the concentration of glyceraldehyde-3-phosphate maintains the inhibition on the ED pathway which could not be overcome by a 40% increase in the concentration of 6-phosphogluconate (Hoque et al., 2011).

A total of 2,461 Km and Vmax values were subsequently assembled using the estimated elementary kinetic parameters (Supplementary Table ST13). Contrary to conventional thought, the total number of Michaelis-Menten parameters (i.e., *K*_*m*_ and *V*_*max*_) exceeded the number of elementary kinetic parameters (i.e., *k*_*p*_). This is because the number of elementary kinetic parameters per reaction increases linearly with the number of participating species (reactants, products or regulators) (see section 1 in the Supplementary methods). However, the Michaelis-Menten formalism requires parameters for every combination of species in the rate expression yielding a quadratic increase in parameters (Cleland, 1963). For example, for an ordered bi-substrate reaction *A + B ⇌ C + D*, ten elementary kinetic parameters are needed as described in the Supplementary methods. Using the Michaelis-Menten formalism requires two *V*_*max*_ terms (one for the forward reaction and one for the reverse reaction) and ten *K*_*m*_ terms (four *K*_*m*_s for individual metabolite concentrations [*A*], [*B*], [*C*] and [*D*] and six *K*_*m*_s one for each pair [*A*][*B*],[*C*][*D*], [*A*][*C*], [*B*][*D*] and triplet [*A*][*B*][*C*] and [*B*][*C*][*D*]). Note that the Michaelis-Menten formalism requires two fewer parameters than using elementary kinetics for a reversible, uni-uni reaction (Cleland, 1963) and only one fewer parameter for a bi-uni or uni-bi reaction. However, for bi-bi and higher-order reaction mechanisms, the Michaelis-Menten formalism always requires more parameters than the elementary kinetics description (Cleland, 1963). These two competing effects yield the relatively close number of elementary and MM parameters for k-ecoli307. Nevertheless, the true number of Michaelis-Menten parameters would have been much larger than the ones required for the elementary kinetics description if complete cofactor metabolism, protons, water and phosphate groups were included in k-ecoli307. The larger number of Michaelis-Menten parameters may explain the reported highly correlated Michaelis-Menten parameters leading to multicollinearity (Heijnen and Verheijen, 2013).

At the optimal solution, the Hessian matrix computed by K-UPDATE represents the inverse of the covariance matrix computed in linear regression (Wiechert et al., 1997). The positive square root of the diagonal terms of this covariance matrix correspond to the standard deviation, in the kinetic parameters. The relative uncertainty in parameter estimation is defined as the ratio of standard deviation computed from the covariance matrix to the estimated parameter value. The standard deviation conceptually quantifies the sensitivity of the SSR objective function to local changes in the value of the inferred parameter. Large values for the standard deviation imply insensitivity of the SSR objective function to changes in the value of the inferred parameter. Note that this inference of standard deviation using linear regression is a local approximation and generally should be viewed as an overestimate of the true uncertainty as it captures neither covariances nor nonlinear effects for larger changes in parameter values. Using this method, the standard deviation representing the uncertainty in estimation for WT reverse elementary fluxes, enzyme fractions, and elementary kinetic parameters was computed. All 1,129 reverse elementary fluxes in the WT network were resolved with a relative uncertainty (*σ*_*v*_/*v*_*r*_) of less than 1%. Furthermore, the average standard deviation for the inferred enzyme fractions (*σ*_*e*_) was only 0.04 mol/mol-total enzyme. A total of 793 out of 1,238 enzyme fractions had a standard deviation of less than 0.1 mol/mol-total enzyme and were resolved with high precision (Supplementary Figure S4a). However, 208 enzyme fractions (mostly with near zero values) were inferred with a standard deviation exceeding the corresponding parameter value leading to a relative uncertainty (*σ*_*e*_/*e*) exceeding 100% (Figure 2a) due to their small absolute value. These enzyme fractions include fractional abundances corresponding to 126 free enzymes and 19 inhibitor-bound enzyme complexes from the TCA cycle and aspartate metabolism, and 63 metabolite-bound enzyme complexes in the ED pathway, TCA cycle and aspartate metabolism. The large relative uncertainty in inference for these enzyme fractions alludes to insufficient fluxomic data for precise kinetic parameterization of these pathways. Uncertainty of inference propagates from the enzyme fractions to the elementary kinetic parameters, leading to 393 out of 2,367 elementary kinetic parameters having a relative uncertainty (*σ*_*kp*_/*k*_*p*_) exceeding 100% (Figure 2b). Notably, 50 out of 69 kinetic parameters describing inhibition of enzyme catalysis were resolved with a relative uncertainty of less than 10%. Well resolved inhibition kinetic parameters include the inhibition of the oxidative pentose phosphate pathway by NADPH, product inhibition of the ED pathway by glyceraldehyde-3-phosphate, and product inhibition of cis-Aconitase by isocitrate. The narrow standard deviation also places a non-zero lower bound on the *1σ*-confidence interval for these inhibition constants implying the essentiality of regulatory interactions in k-ecoli307 to explain the available experimental flux datasets.

**Figure 2:**
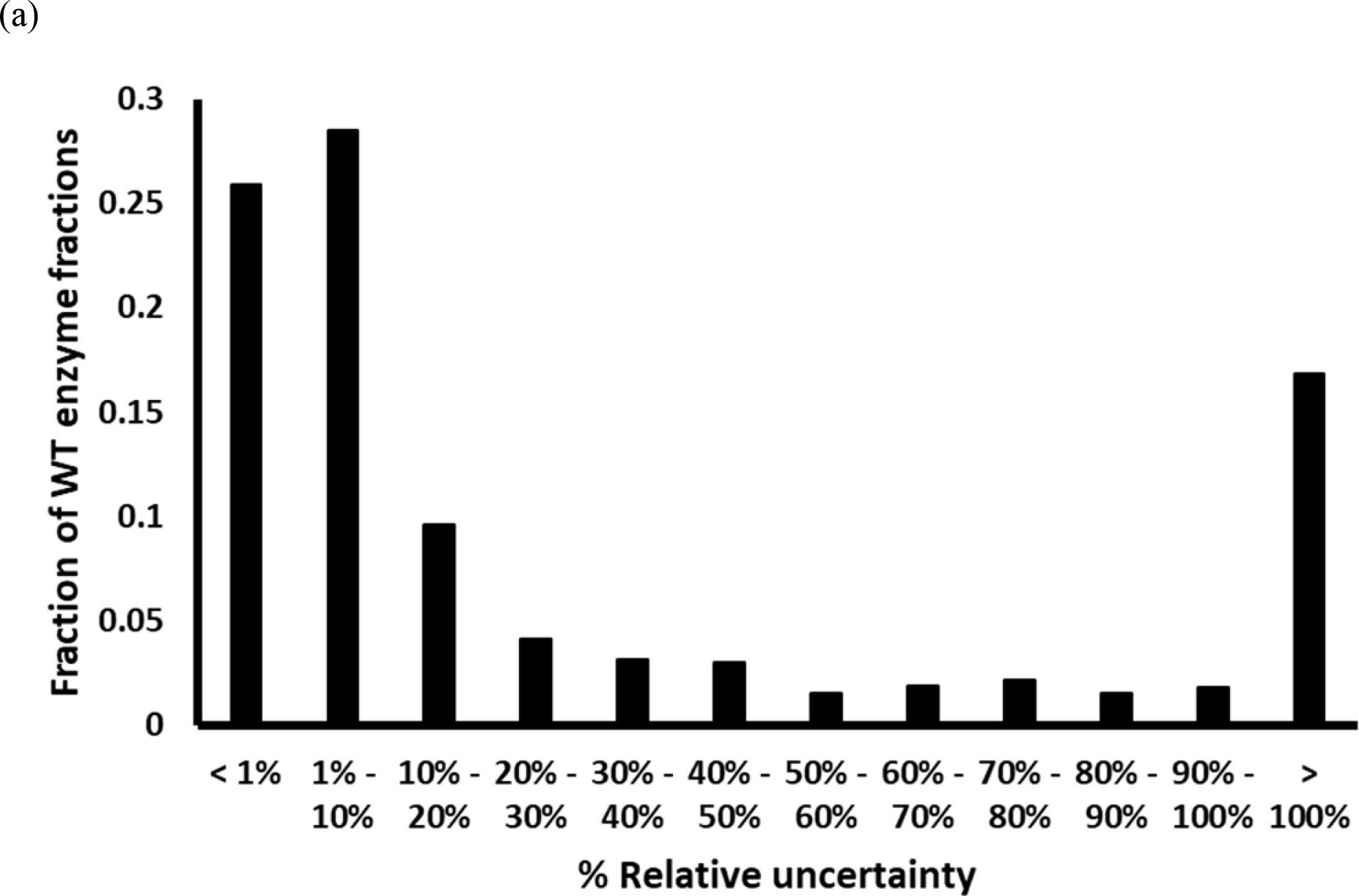
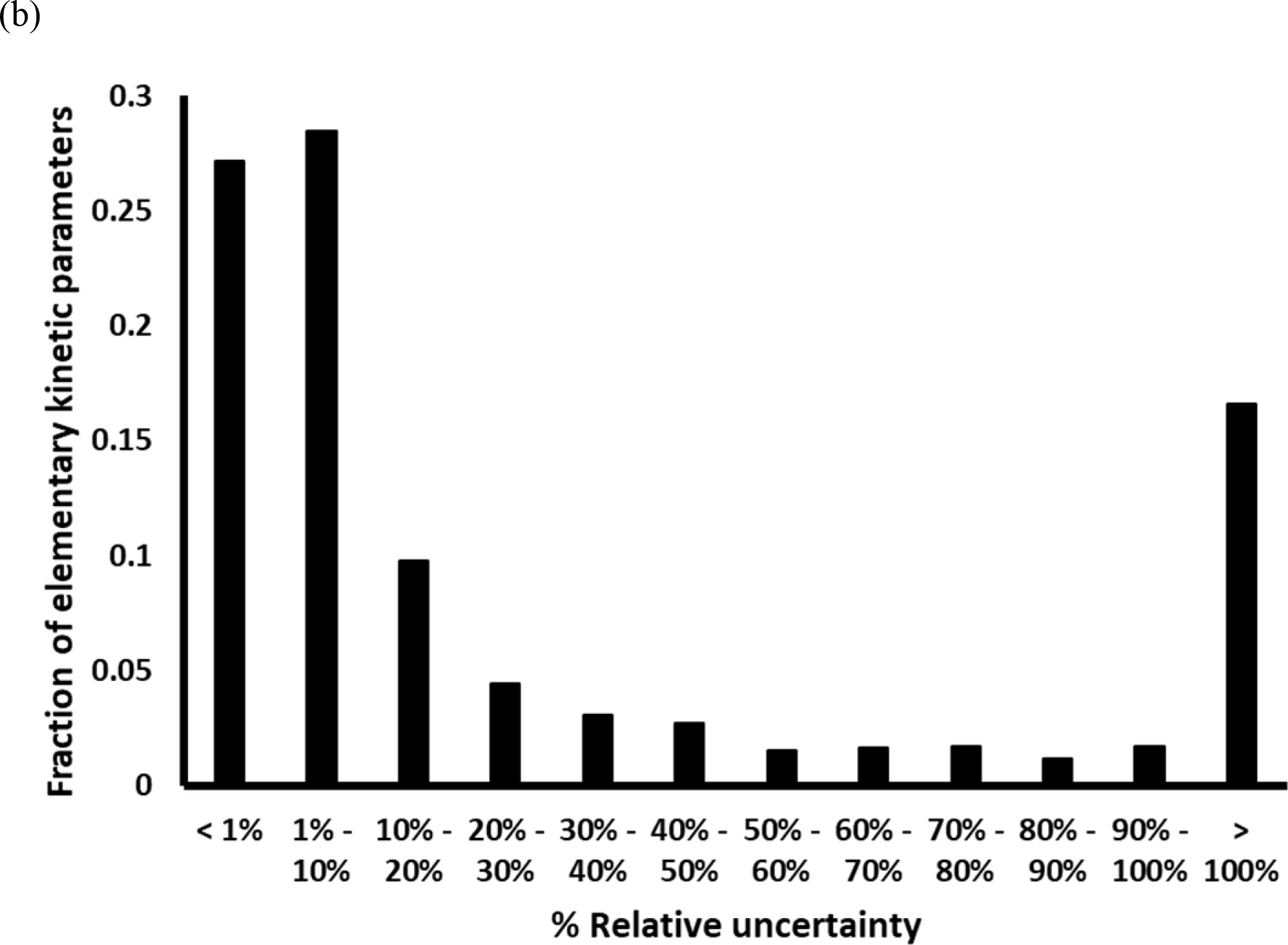
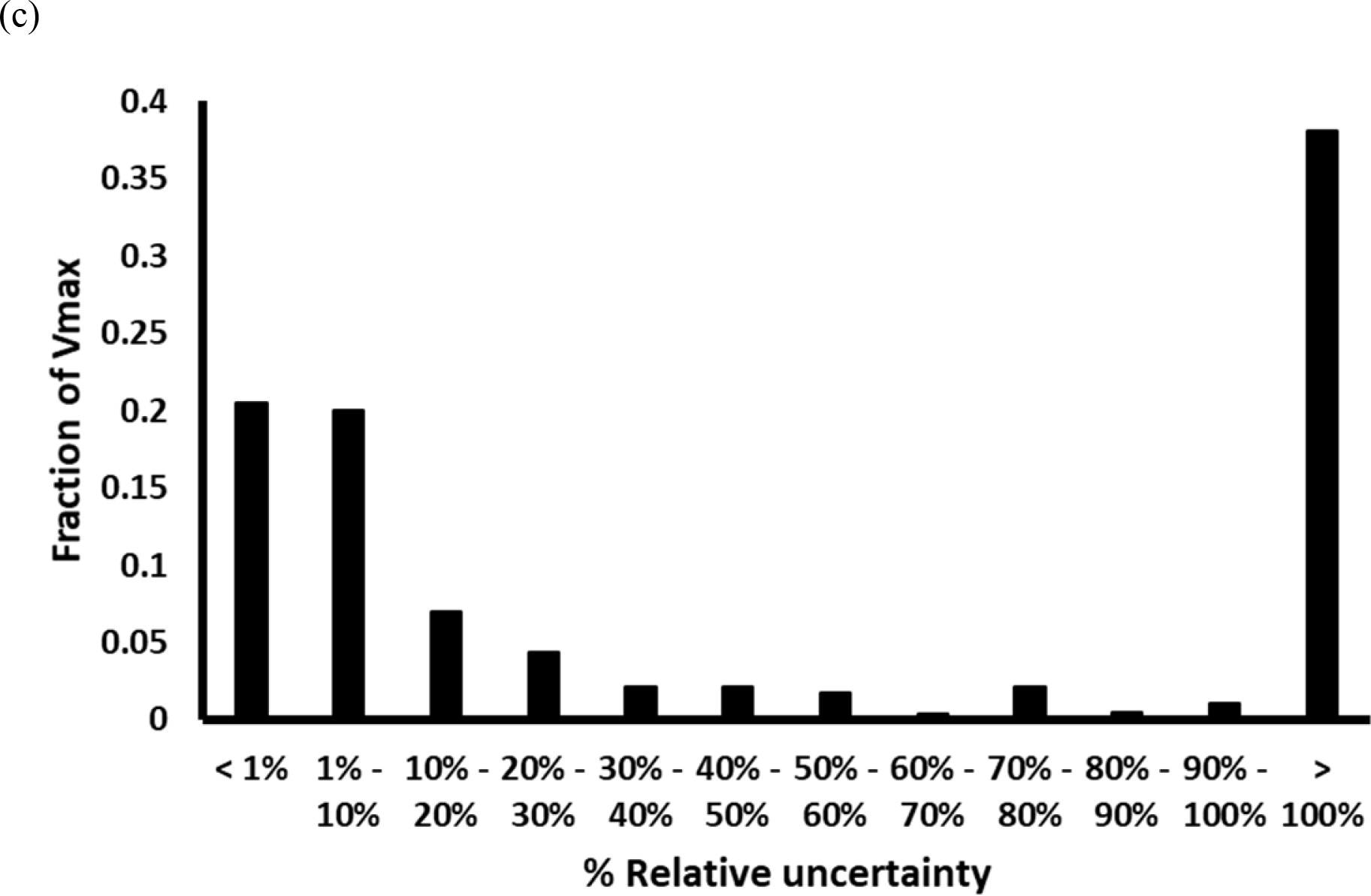
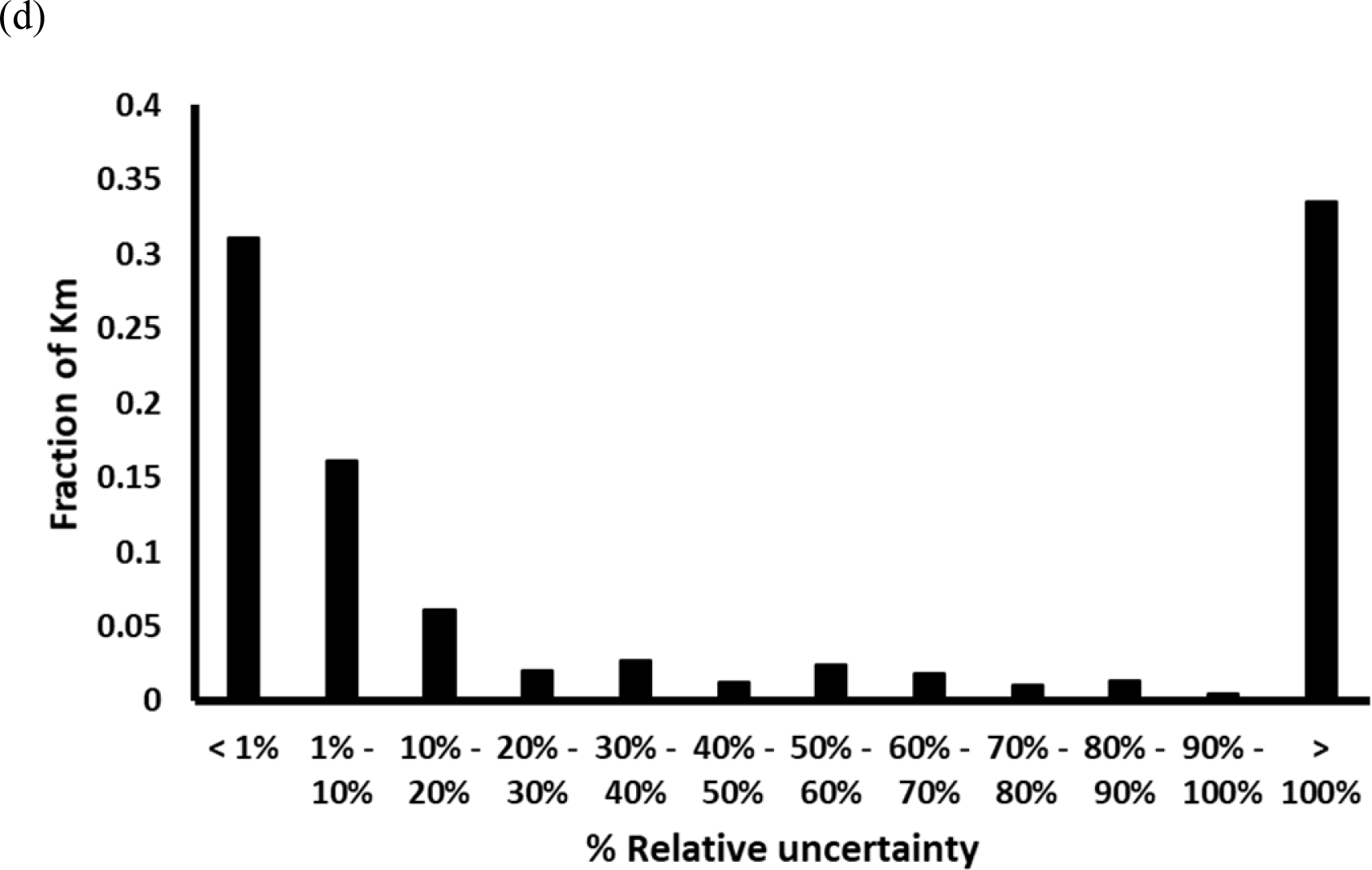
Distribution of relative uncertainty in estimation for (a) WT enzyme fractions, (b) elementary kinetic parameters, (c) *V*_*max*_, and (d) *K*_*m*_ in k-ecoli307. (a) Distribution of values assumed by WT enzyme fractions (red bars) and standard deviations (green bars) for estimated enzyme fractions.

Upon computing the uncertainty in estimation of Michaelis-Menten parameters, 231 of the 570 Vmax parameters from branched-chain amino acid and fatty acid biosynthetic pathways were resolved with a relative uncertainty (*σ*_*Vmax*_/*V*_*max*_) of less than 10% (Figure 2c). In contrast, 217 Vmax parameters from nucleotide biosynthesis, aspartate metabolism, serine metabolism, and the TCA cycle were resolved with a relative uncertainty exceeding 100%. Similarly, 893 of 1,891 Km parameters from the EMP pathway, pentose phosphate pathway, TCA cycle and fatty acid biosynthesis were resolved with a relative uncertainty (*σ*_*Km*_/*K*_*m*_) of less than 10% (Figure 2d), whereas 633 Km parameters from nucleotide biosynthesis, amino acid metabolism and the glyoxylate shunt were resolved with a relative uncertainty exceeding 100%. As expected, the estimation uncertainty for the elementary kinetic parameters propagated to Km and Vmax. However, the fraction of unresolved Michaelis-Menten parameters was substantially higher (33.4% of all Km and 38.1% of all Vmax) than the fraction of poorly resolved elementary kinetic parameters (16.6%). The higher fraction of unresolved Michaelis-Menten parameters stems from the fact that Michaelis-Menten parameters are assembled as a nonlinear combination of elementary kinetic parameters due to which the high relative uncertainty for any one elementary kinetic parameter can propagate to multiple Michaelis-Menten parameters. It is important to note that the approach used for computing standard deviations is based on statistical analysis of linear regression which only holds true in the neighborhood of the optimal solution. The analysis can be confounded by the nonlinear behavior of the kinetic model upon moving further away from the optimal solution. In addition, by looking at a single parameter at a time information of covariances between parameters is lost and the inferred uncertainty significantly overestimates the true parametric uncertainty. A more rigorous profile-likelihood method (Antoniewicz et al., 2006) can be employed to account for the nonlinear structure of the kinetic model that generally results in narrower confidence intervals as commonly seen in 13C-MFA.

The predictive capability of the model was evaluated by comparing the model prediction of product yields in engineered strains with the corresponding experimental yield. The genetic perturbation mutants considered for evaluation of predictive capability were not included in the training dataset for kinetic parameterization. Of the six over-producing strains evaluated, the kinetic model successfully predicted the yields of acetate, malate, and lactate to within 30% of the reported experimental yield (Table 1). This indicates that the genetic perturbations in the training dataset for parameterization and that the regulatory structure of the expanded kinetic model is sufficient to explain the phenotypic response of *E. coli* to perturbations in the EMP pathway. The yield predictions for acetate and malate were superior to those by k-ecoli457 due to the fact that both the training dataset for parameterization of the expanded model and cultivation of the engineered strains were at the same mid-exponential growth phase, whereas, the training dataset for k-ecoli457 was generated during late exponential growth phase. The transcriptomic and fluxomic differences between these two growth conditions limits the carbon flux through acetate metabolism in the late exponential growth phase (Ishii et al., 2007). Unlike predictions for core metabolism, product yields originating from peripheral metabolism were poorly predicted by both models. This is because, growth-coupling of peripheral metabolic pathways limits the flux through these pathways to less than 5 mmol/gdw-h in all mutants. The limited redirection of fluxes within peripheral metabolism when compared to central metabolism leads to insufficient fluxomic data for kinetic parameterization of peripheral metabolic pathways and therefore, adversely impacts the prediction fidelity of both kinetic models.

**Table 1:** Comparison of predicted product yields (mol/mol glucose) with experimental yields in engineered over-producing strains of *E. coli*. The experimental yields and predictions by k-ecoli457 were obtained from previously published data by Khodayari and Maranas (Khodayari and Maranas, 2016)

| Product | Perturbed Enzyme | Predicted Yield | Predicted Yield (k-ecoli457) | Experimental Yield |
| --- | --- | --- | --- | --- |
| Acetate | 0.1x RPI | 0.93 | 0.2 | 0.75 |
| L-Valine | 0.1x THRD | 0.03 | 0.02 | 0.34 |
| Lactate | 0x ACKr | 1.4 | 1.11 | 1.13 |
| Malate | 0.3x PTA;<br>10x PPCK | 0.16 | 0.84 | 0.15 |
| Artemisinin | 2x PDH | 0.17 | 0.03 | 0.38 |
| Naringenin | 2xACCOAC | 0.026 | 0.012 | 0.008 |

## Discussion

This manuscript details the development of K-FIT, an accelerated kinetic parameterization algorithm based on steady-state fluxomic data. The K-FIT algorithm estimates kinetic parameters by solving a nonlinear least-squares minimization problem to recapitulate experimentally measured steady-state metabolite concentrations and fluxes using an iterative loop comprised of three steps: K-SOLVE, SSF-Evaluator, and K-UPDATE. The computational savings afforded by bypassing ODE integration improves parameterization speed of K-FIT by over three orders of magnitude compared to the GA-based EM procedure for a core model of metabolism containing 953 kinetic parameters. These savings are likely to become even more pronounced for larger models. These computational savings enable the evaluation of reproducibility of kinetic parameterization within reasonable time even for large-scale models, which was previously not possible using meta-heuristic methods. The parallelizable architecture of SSF-Evaluator improves the scalability of the procedure while allowing compatibility with GPU-based computing architectures which affords significant improvements in computation speed. The iterative scheme presented in SSF-Evaluator is inherently numerically stable which allows it to handle stiff systems of equations with ease while permitting reliable calculation of first- and second-order gradients. Furthermore, the ability to calculate gradients enables local statistical analysis of the inferred kinetic parameters.

The applicability of K-FIT to large-scale models was demonstrated using an expanded kinetic model of *E. coli* containing 307 reactions, 258 metabolites, and 2,367 kinetic parameters, parameterized using fluxes elucidated using 13C-MFA. In order to avoid any error propagation arising from flux projection from simpler models, a recently developed two-step computational pipeline (Foster et al., 2019 (Under Review)) was used for kinetic parameterization using 13C-labeling data. First, fluxes were elucidated for the expanded model in the WT and six single gene-deletion strains using 13C-MFA. The elucidated fluxes were then used to parameterize the kinetic model corresponding to the same stoichiometric model. Although the expanded kinetic model recapitulated the fluxes better than a core model for *E. coli*, product yield predictions in engineered strains did not differ significantly compared to those predicted by the core model. Compared to k-ecoli457, yield predictions for acetate and malate were better recapitulated by k-ecoli307 because k-ecoli307 was parameterized using fluxomic data generated in the mid-exponential growth phase. This similarity in growth conditions between the training dataset and the engineered strains limits any proteome re-allocations that could lower the predictive capabilities of parameterized kinetic models. Yield predictions for metabolites from peripheral metabolism by both k-ecoli307 and k-ecoli457 differed from experimental yields by almost 100% (Table 1). This was traced back to insufficient fluxomic data from peripheral metabolism across the considered mutants due to growth coupling. Since most amino acids are not catabolized by *E. coli*, reliable parameterization of these pathways requires model expansion to amino acid pool turnover by protein synthesis and degradation. Additional fluxomic and metabolomic data from overproducing strains and auxotrophs will also help capture the link between genetic perturbations and increased flux through peripheral metabolism as the WT strain of *E. coli* does not secrete any amino acids during the mid-exponential growth phase.

A novel feature introduced by K-FIT is the rapid computation of local sensitivity of predicted fluxes and metabolite concentrations to kinetic parameters, which enables easy computation of the gradient and Hessian of the least-squares objective function for efficient traversal of the feasible kinetic parameter space. Since the Hessian represents the inverse of the covariance matrix in the neighborhood of the optimal solution, K-FIT is able to compute uncertainty in parameterization estimation with no added cost. Interestingly, reverse elementary fluxes in the WT strain were precisely estimated with a mean relative uncertainty of less than 1%. WT enzyme fractions were also resolved with a mean standard deviation of 0.04 mol/mol-total enzyme. However, the low abundance of 208 enzyme complexes relative to their corresponding standard deviation leads to a relative uncertainty exceeding 100%. This large relative uncertainty propagates to elementary kinetic parameters and Michaelis-Menten parameters due to nonlinear mapping of WT enzyme fractions to kinetic parameters. It is important to note that uncertainty analysis based on linear regression is accurate only near the optimal solution and a rigorous analysis accounting for the nonlinear behavior of the kinetic model will be required to compute the accurate confidence intervals further away from the optimal solution. Accurate confidence intervals will provide insights into resolvability of kinetic parameters for the set of experimental data and enable the identification of informative mutants (Zomorrodi et al., 2013) and design of experiments (Banga and Balsa-Canto, 2008) to pin down the poorly resolved kinetic parameters. Furthermore, using accurate confidence intervals, additional insights into reaction reversibility and importance of regulatory interactions can be gleaned. Currently, the statistical significance of regulatory interactions can only be evaluated using frameworks such as SIMMER (Hackett et al., 2016)

Overall, K-FIT highlights the data-demanding nature of the kinetic parameterization problem. Although kinetic parameterization was performed using only steady-state flux data, steady-state metabolite concentration data can also be included in the SSR objective function. In all studies enzyme levels were assumed to remain the same in mutants as in WT with the exception of enzymes that are associated with knock-out genes which were set to zero. Nevertheless, enzyme levels in mutant strains can be pre-specified in K-FIT if the information is known *a priori*. Ideally, one would want to integrate allosteric with transcriptional regulation so that the enzyme concentrations in the mutant networks can be related to the altered metabolite concentrations (Fuhrer et al., 2017). This would ultimately enable the integration of mutant network data generated under both genetic and environmental perturbations and improve its predictive capabilities. Furthermore, the local sensitivity of fluxes and metabolite concentrations with respect to kinetic parameters directly map to elasticity coefficients used in metabolic control analysis. They can thus be used to calculate flux and concentration control coefficients at minimal additional cost to inform metabolic engineering strategies.

## Methods

### Kinetic parameterization using K-FIT

K-FIT is a gradient-based kinetic parameterization algorithm that minimizes the least-squares objective function representing the weighted squared deviation between predicted and measured steady-state metabolic fluxes (and possibly metabolite concentrations) across multiple genetic perturbation mutants. The full mathematical description for the K-FIT algorithm is provided in the supplementary methods. The least-squares NLP is solved using the Levenberg-Marquardt algorithm (Madsen et al., 2004) in conjunction with the active-set method for enforcing linear inequality constraints (Gill et al., 1984). K-FIT is encoded and implemented in MATLAB^TM^ and run on an Intel-i7 (4-core processor, 2.6GHz, 12GB RAM) computer. K-FIT is tested using kinetic models at different size scales. The full source-code is made available on GitHub.

Computation of standard deviations for estimated kinetic parameters was performed using linear regression tools applied to a local linearization of mutant fluxes using Taylor series expansion (Wiechert et al., 1997). Briefly, the Covariance matrix ***C*** is computed by inverting the Hessian ***H*** computed by K-UPDATE. When the linear approximation holds, the diagonal of the covariance matrix represents the estimation variance of kinetic parameters. The approximate standard deviation of kinetic parameter *k*_*p*_ (*σ*_*p*_) is evaluated as 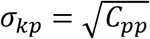. The approximate *1σ* confidence interval is computed as *k*_*p*_ ± *σ*_*kp*_. Note that this method of analysis does not capture covariance between parameters which is expected to significantly lower uncertainty.

### Construction of the expanded kinetic model of *E. coli*, k-ecoli307

The expanded metabolic model is constructed by de-lumping the central and peripheral metabolic pathways in the core model (Foster et al., 2019 (Under Review)) based on the reported biomass composition (Neidhardt and Curtiss, 1996). The expanded model contains 307 reactions and 258 metabolites. Atom mapping for the additional reactions were obtained from the previously published genome-scale carbon mapping model for *E. coli* (Gopalakrishnan and Maranas, 2015). The amino acid labeling data for flux elucidation was obtained from the published work by Long *et al* (2018). Metabolic fluxes and 95% confidence intervals were elucidated using 13C-metabolic flux analysis as described earlier (Antoniewicz et al., 2006; Gopalakrishnan and Maranas, 2015). The mechanism and allosteric regulation of enzyme-catalyzed reactions in the model were obtained from k-ecoli457, the near-genome-scale kinetic model for *E. coli* (Khodayari and Maranas, 2016). The standard deviation *σ*_*j*_ corresponding to the estimated flux *V*_*j*_ to be used as a weighting factor in the K-FIT algorithm is computed from the lower and upper bounds of the confidence interval 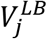 and 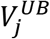 reported by 13C-MFA as 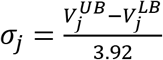. The input files containing the model, reaction mechanism descriptions, and experimental data used for kinetic parameterization are provided in Supplementary Tables ST13 – ST16. Computed kinetic parameters were also packaged into Michaelis-Menten parameters as described earlier (Khodayari and Maranas, 2016). The estimated kinetic parameters and Michaelis-Menten parameters are reported in Supplementary Tables ST21 and ST22.

## ACKNOWLEDGEMENT

This work was supported by the National Science Foundation at the Pennsylvania State University, University Park, under grant NSF/MCB-1615646, by the US Department of Energy (DOE) under grants DE-SC0018260, and by the Center for Bioenergy Innovation under the US Department of Energy (DOE) under grant DE-AC05-000R22725.

---

**Figure S1:**
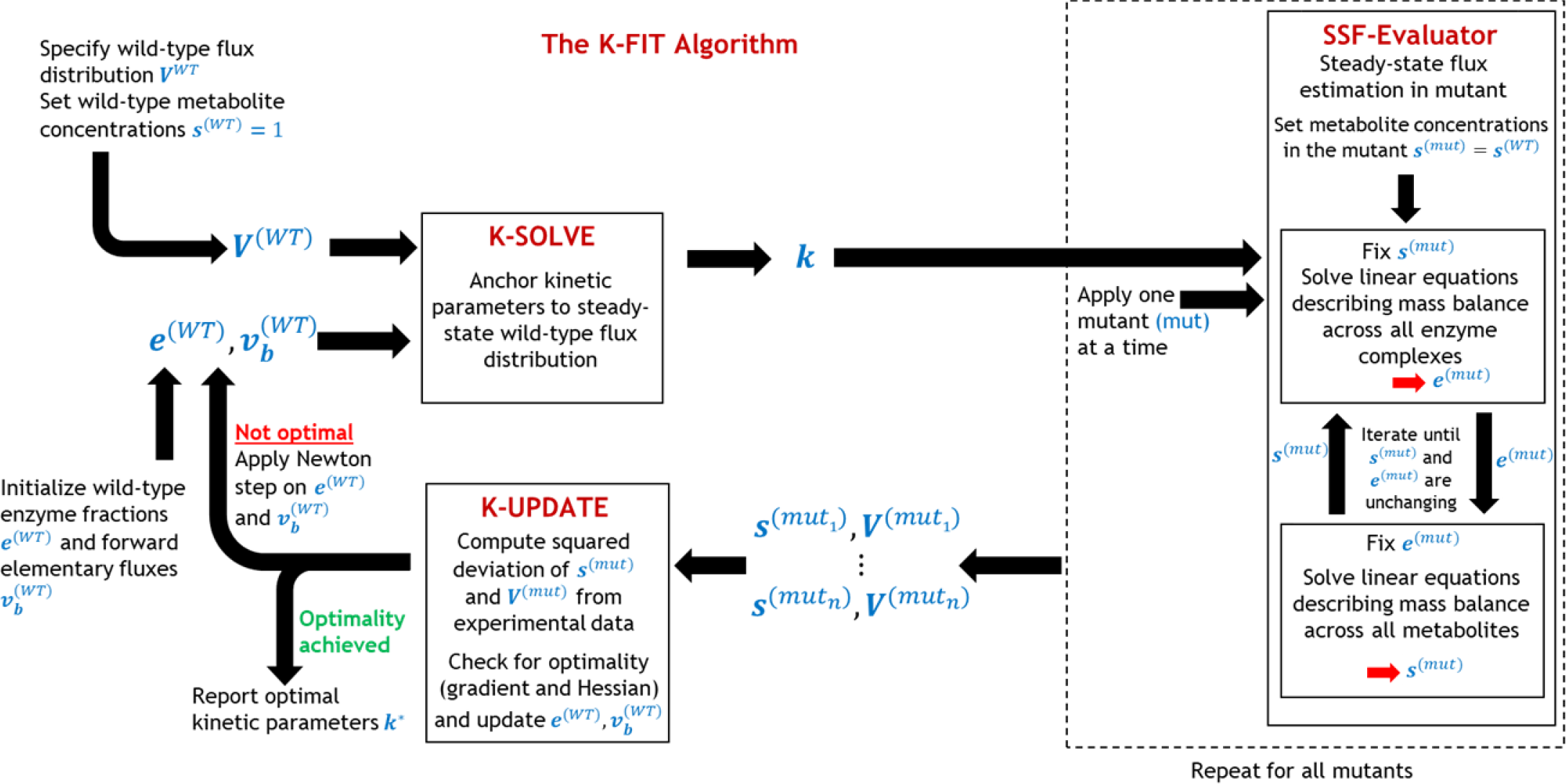
Overview of the K-FIT algorithm showing the flow of information between various components.

**Figure S2:**
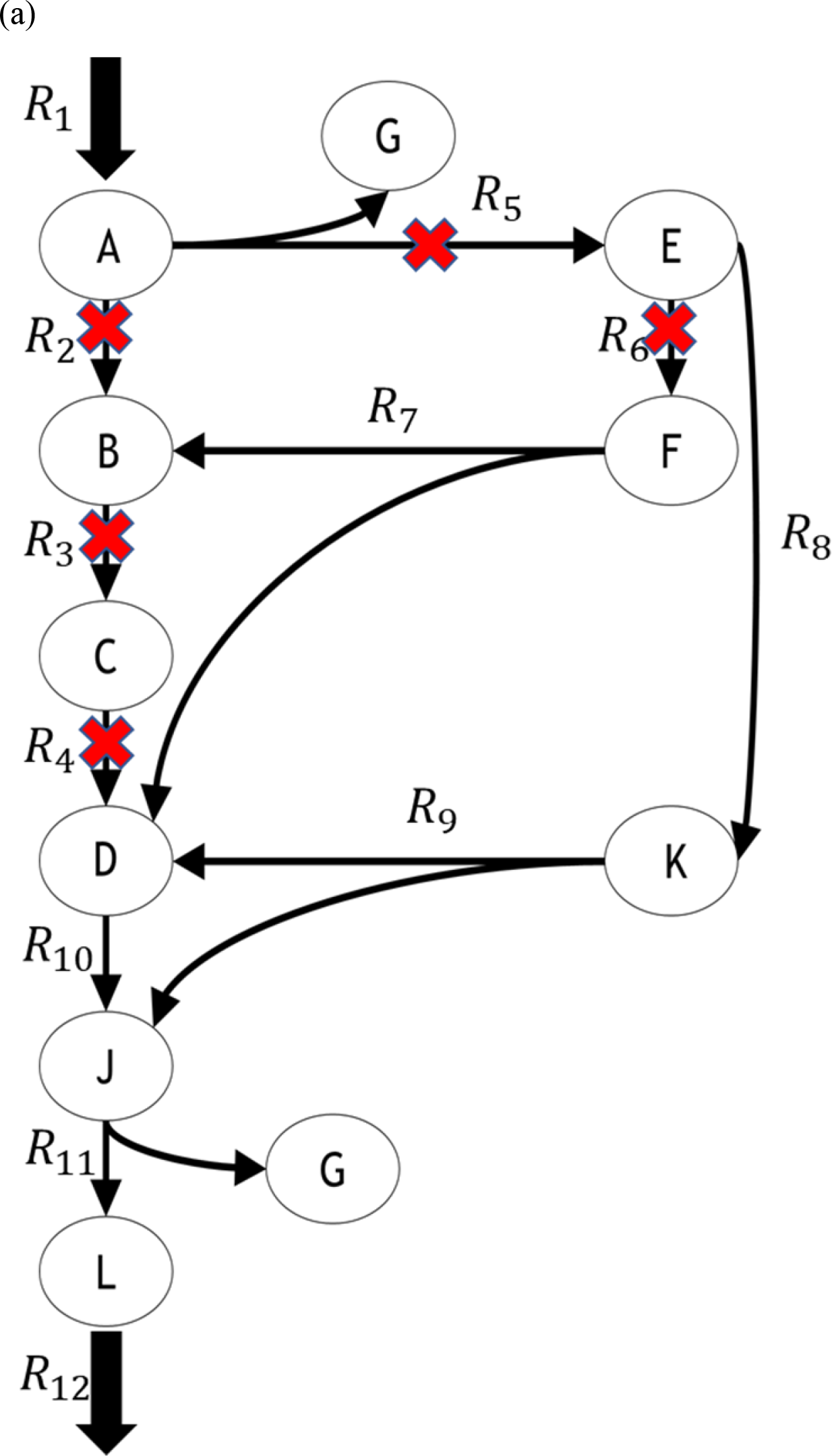
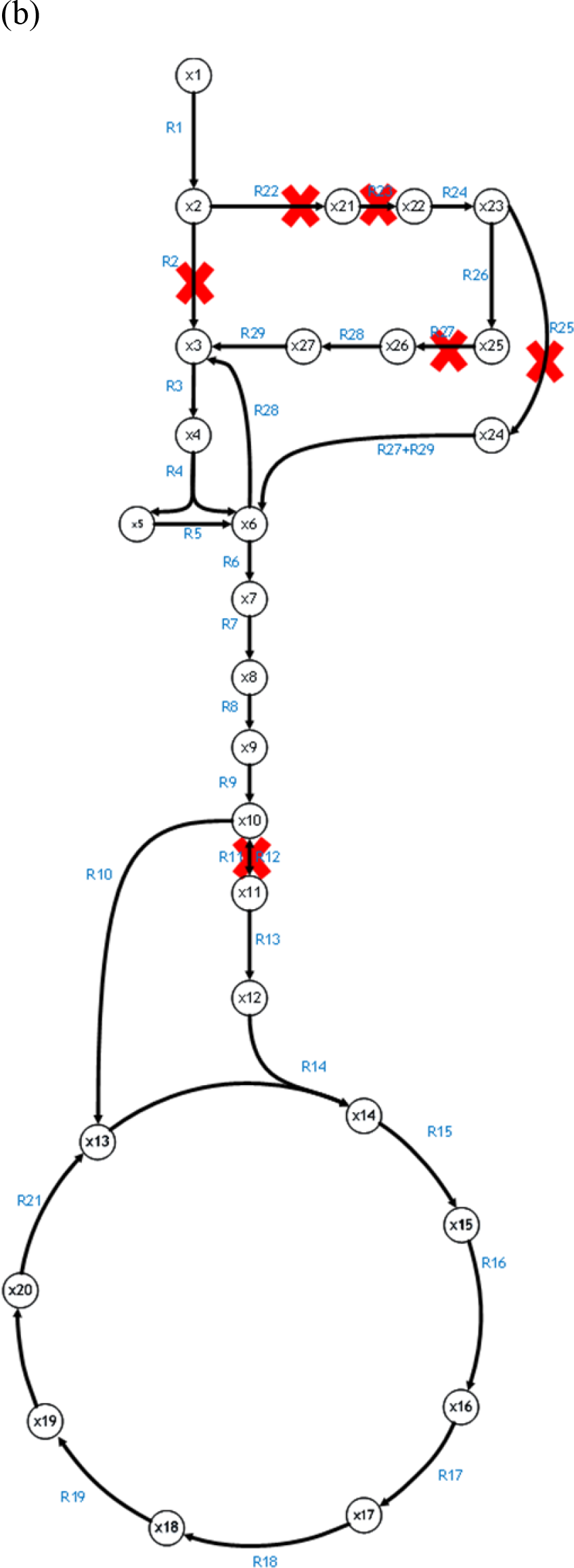
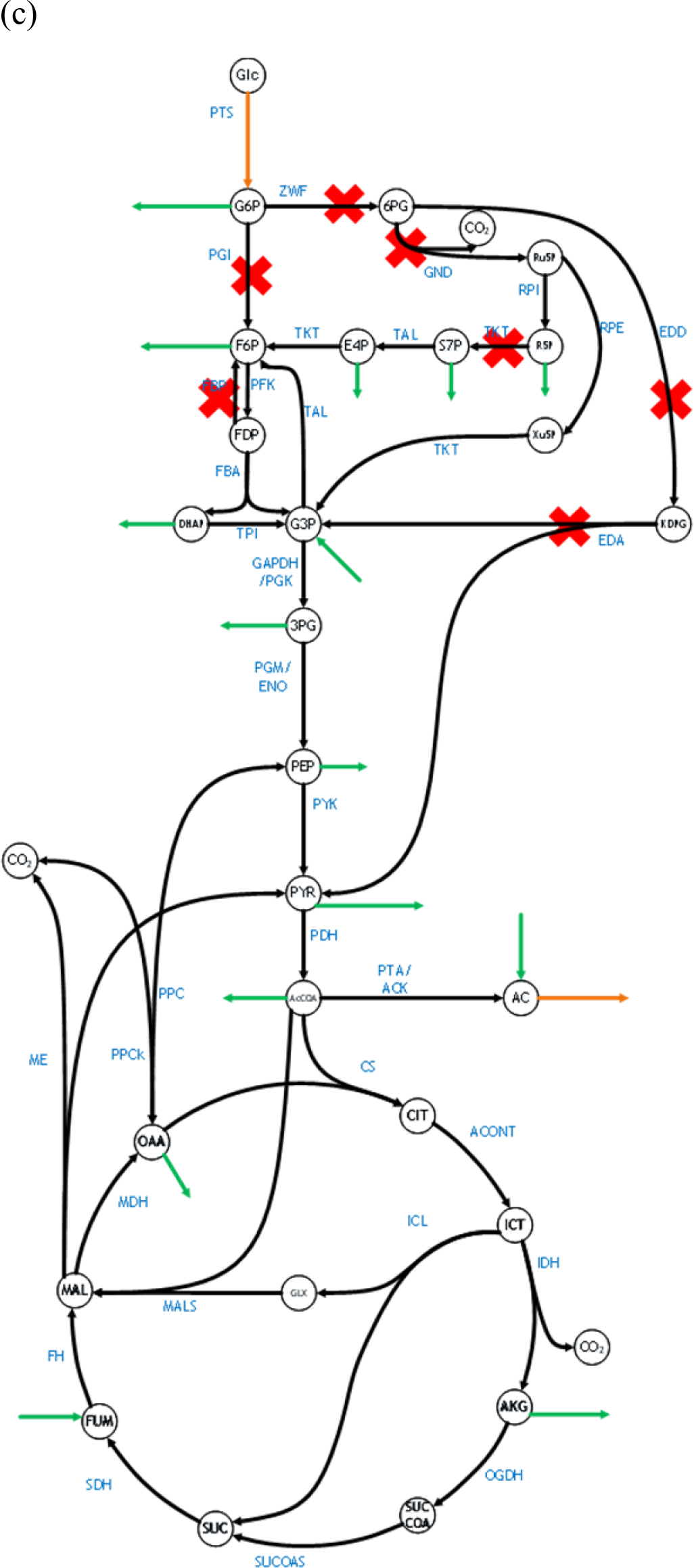
Test models used to benchmark the performance of K-FIT against GA-based EM procedure. (a) Small model containing 14 reactions and 11 metabolites. (b) Medium-sized model containing 33 reactions and 28 metabolites. (c) Core model containing 108 reactions and 65 metabolites. Reactions knocked out in the single gene-deletion mutants are indicated using a red X.

**Figure S3:**
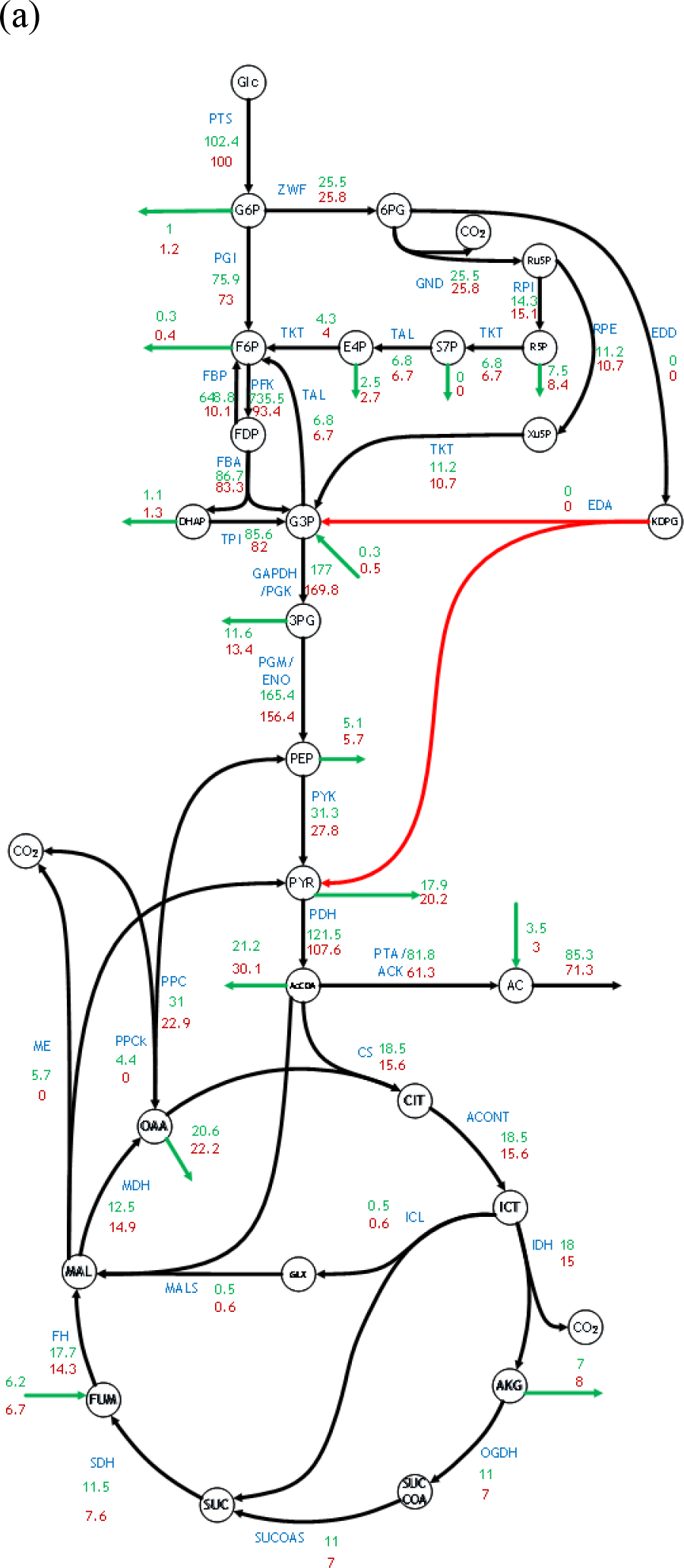
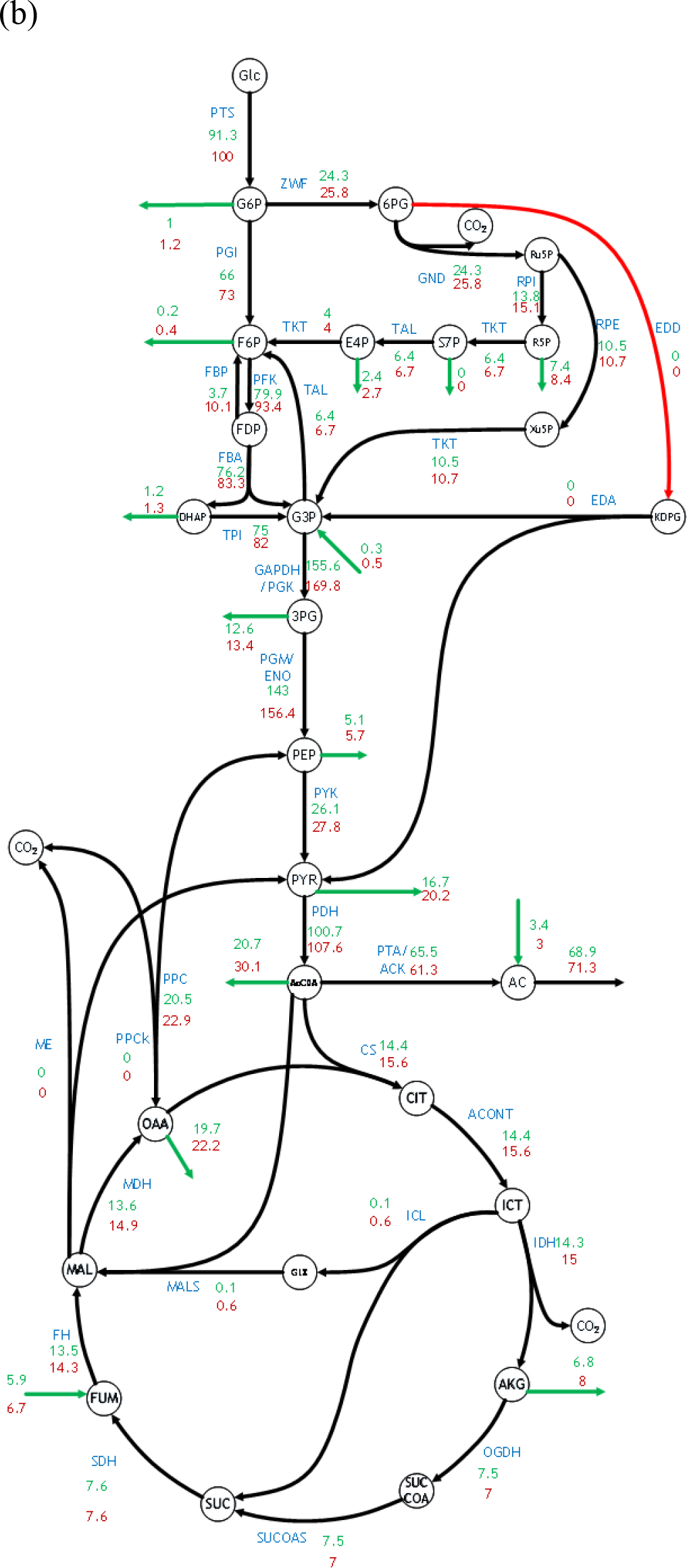
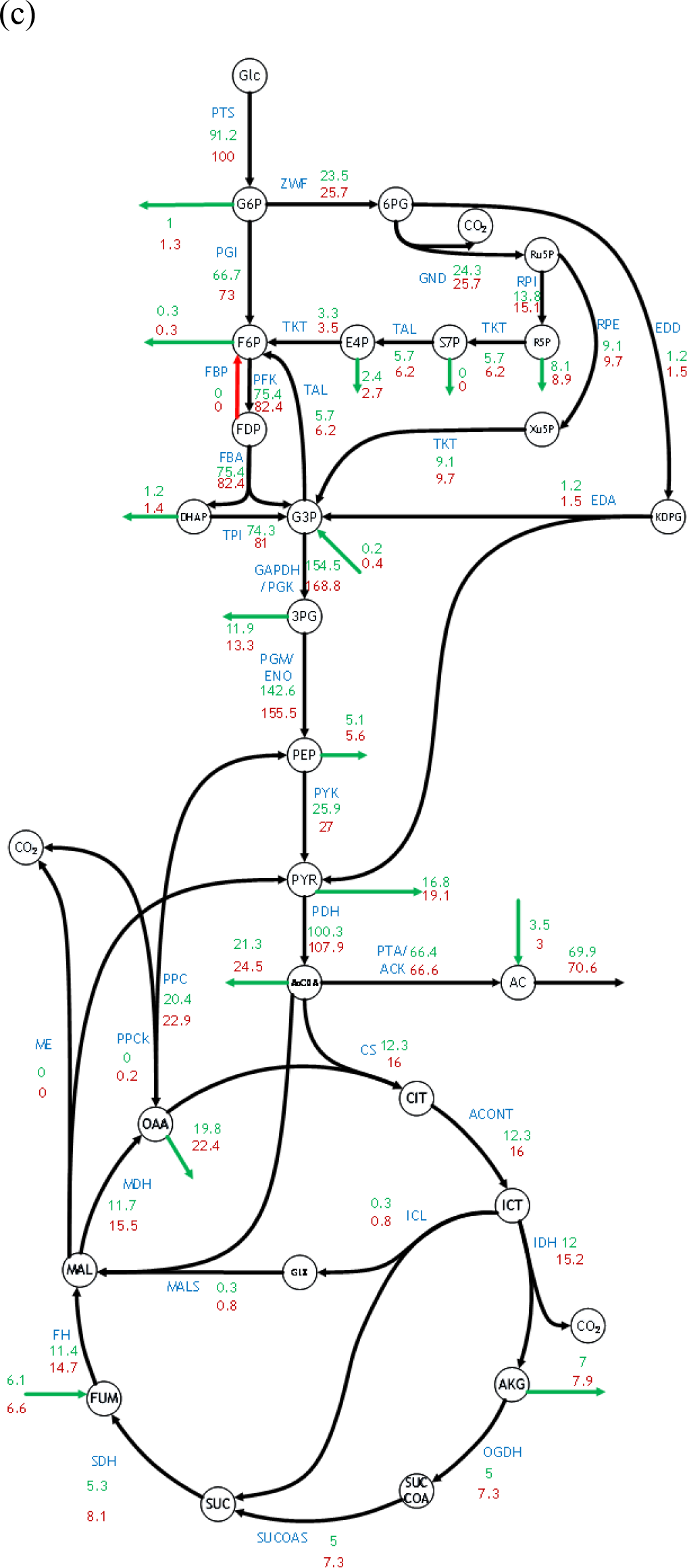
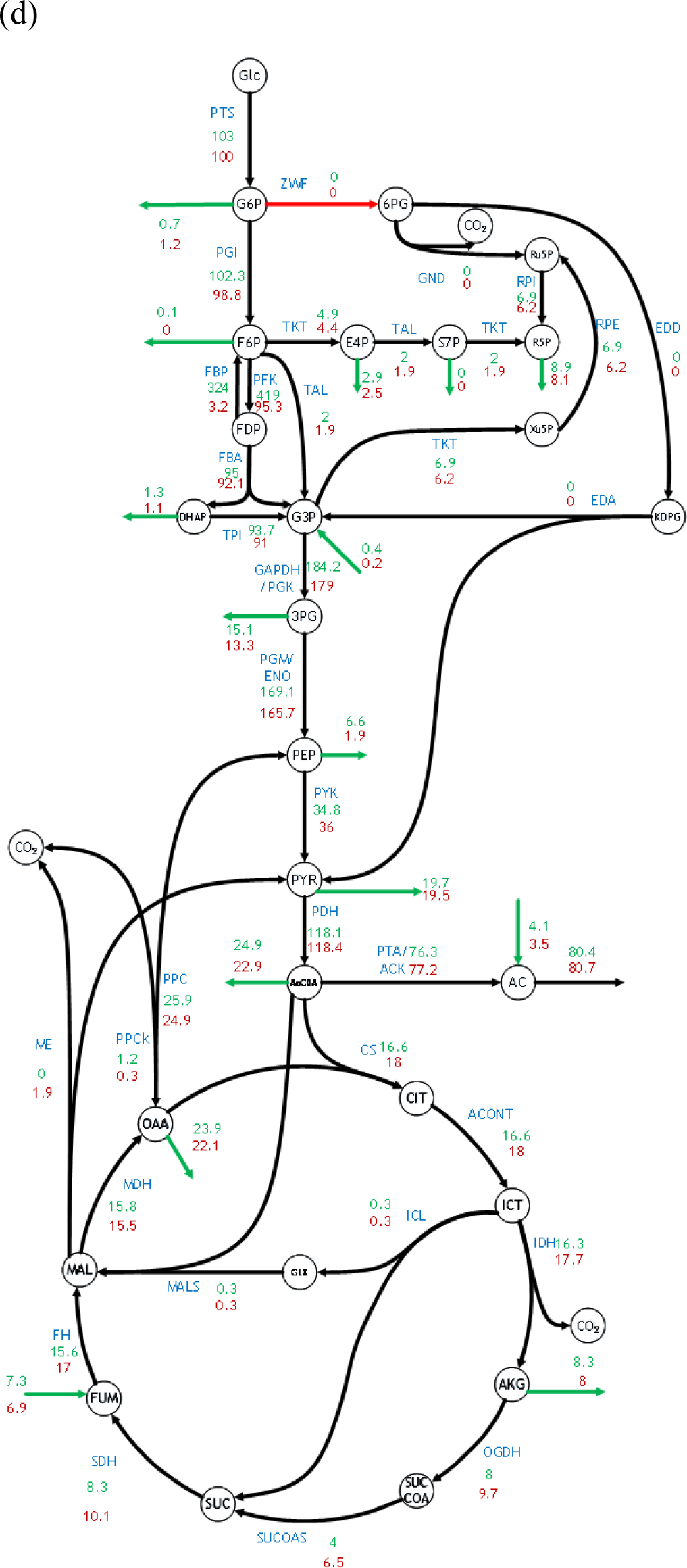
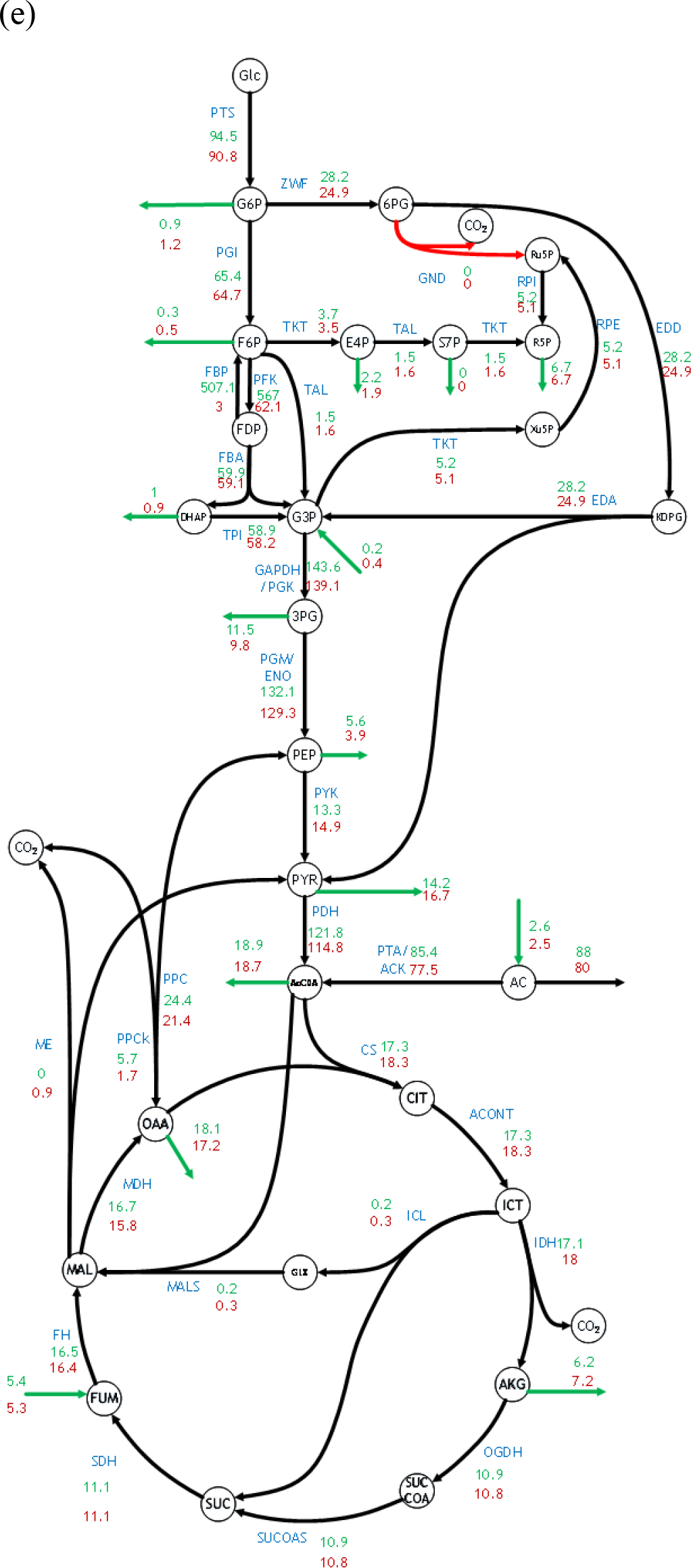
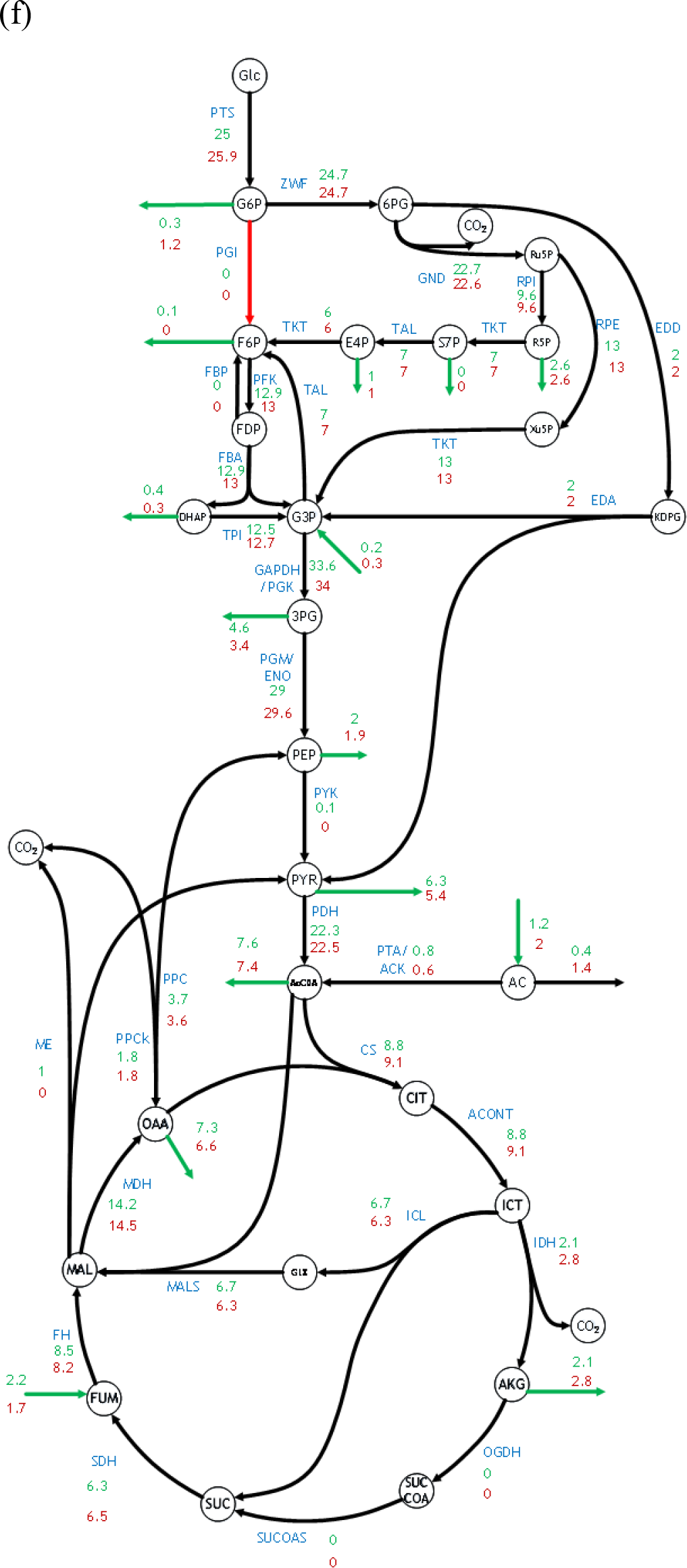
Flux distribution through central metabolism of the expanded model for *E. coli* in (a) Δ*eda*, (b) Δ*edd*, (c) *Δfbp*, (d) *Δzwf*, (e) Δ*gnd*, and (f) *Δpgi* mutant strains. Reactions representing metabolite flows between central and peripheral metabolism are indicated using green arrows. Fluxes elucidated using 13C-MFA are shown in green and the corresponding flux prediction by the expanded kinetic model is shown in brown. Reactions corresponding to the knocked-out genes in each mutant strain are indicated using red arrows. Flux measurements for PFK and FBP were not fitted due to poor resolution 13C-MFA

**Figure S4:**
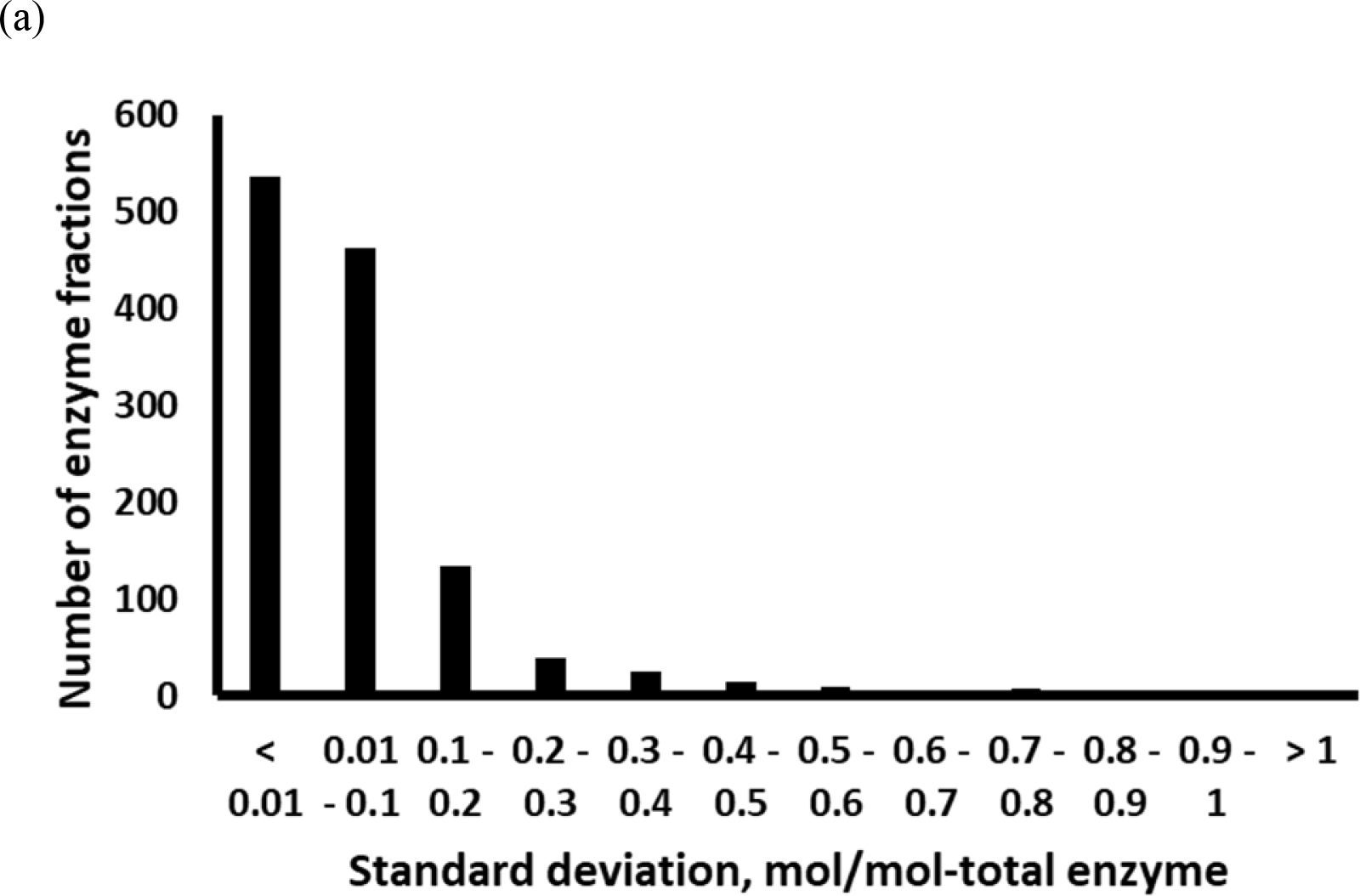
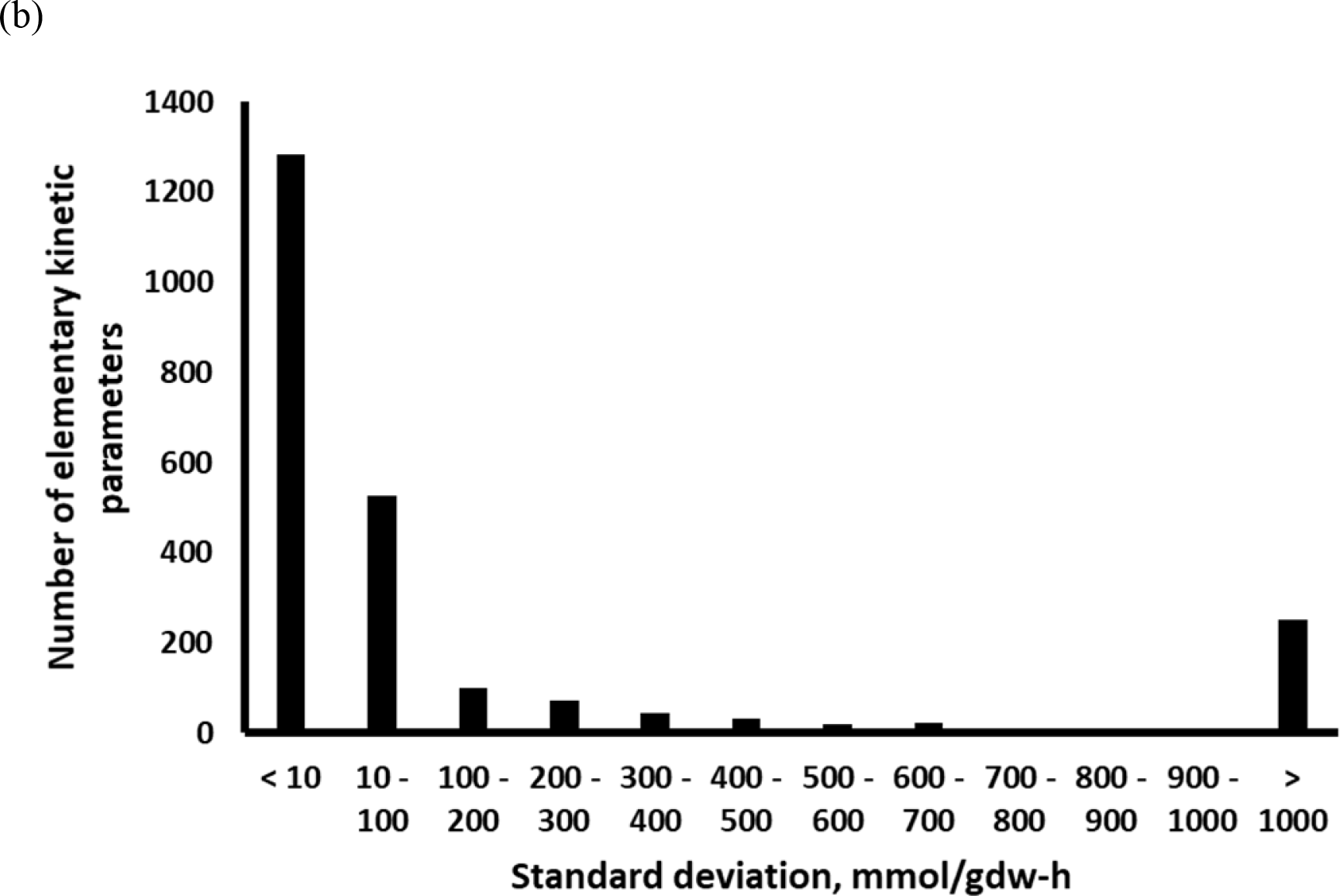
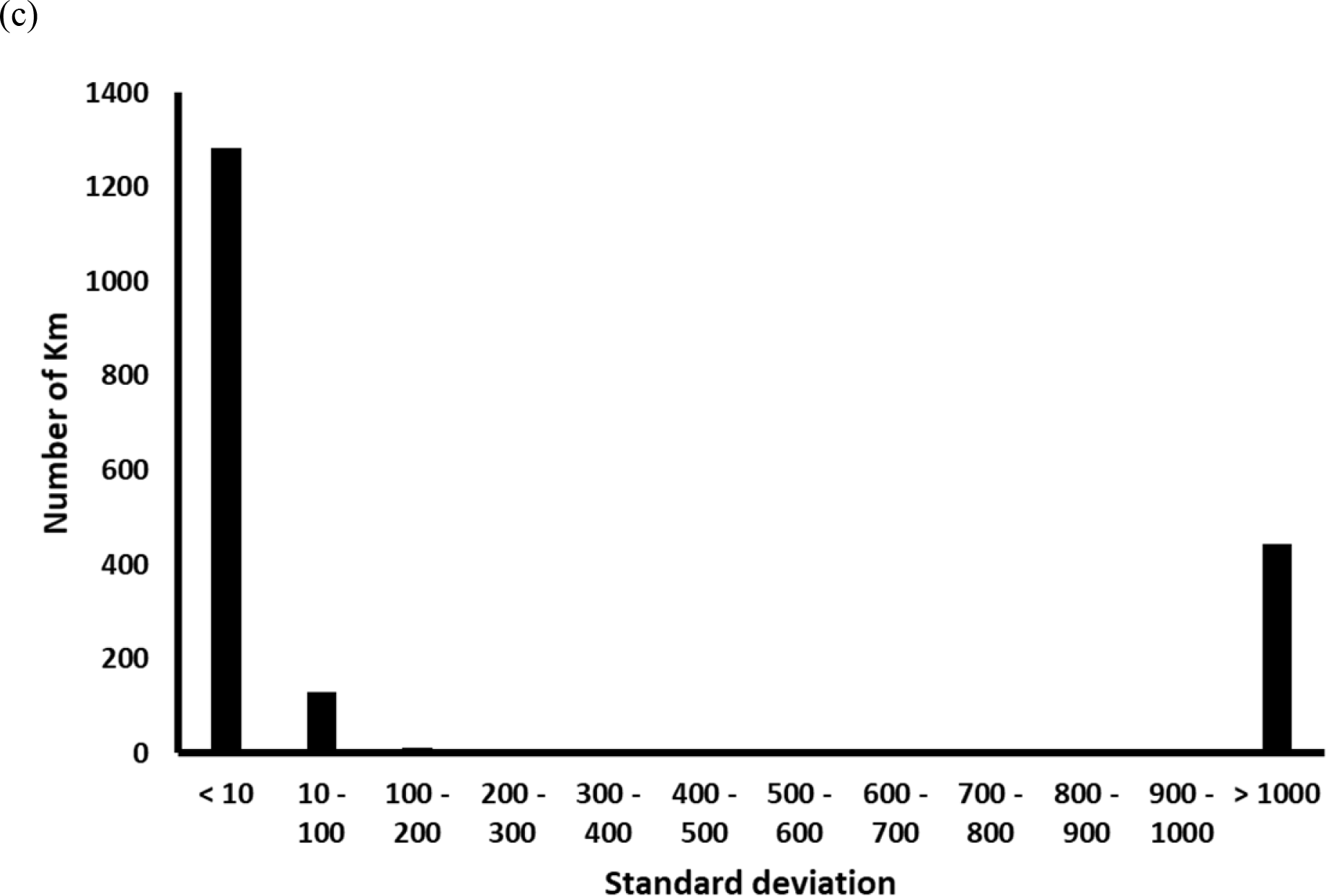
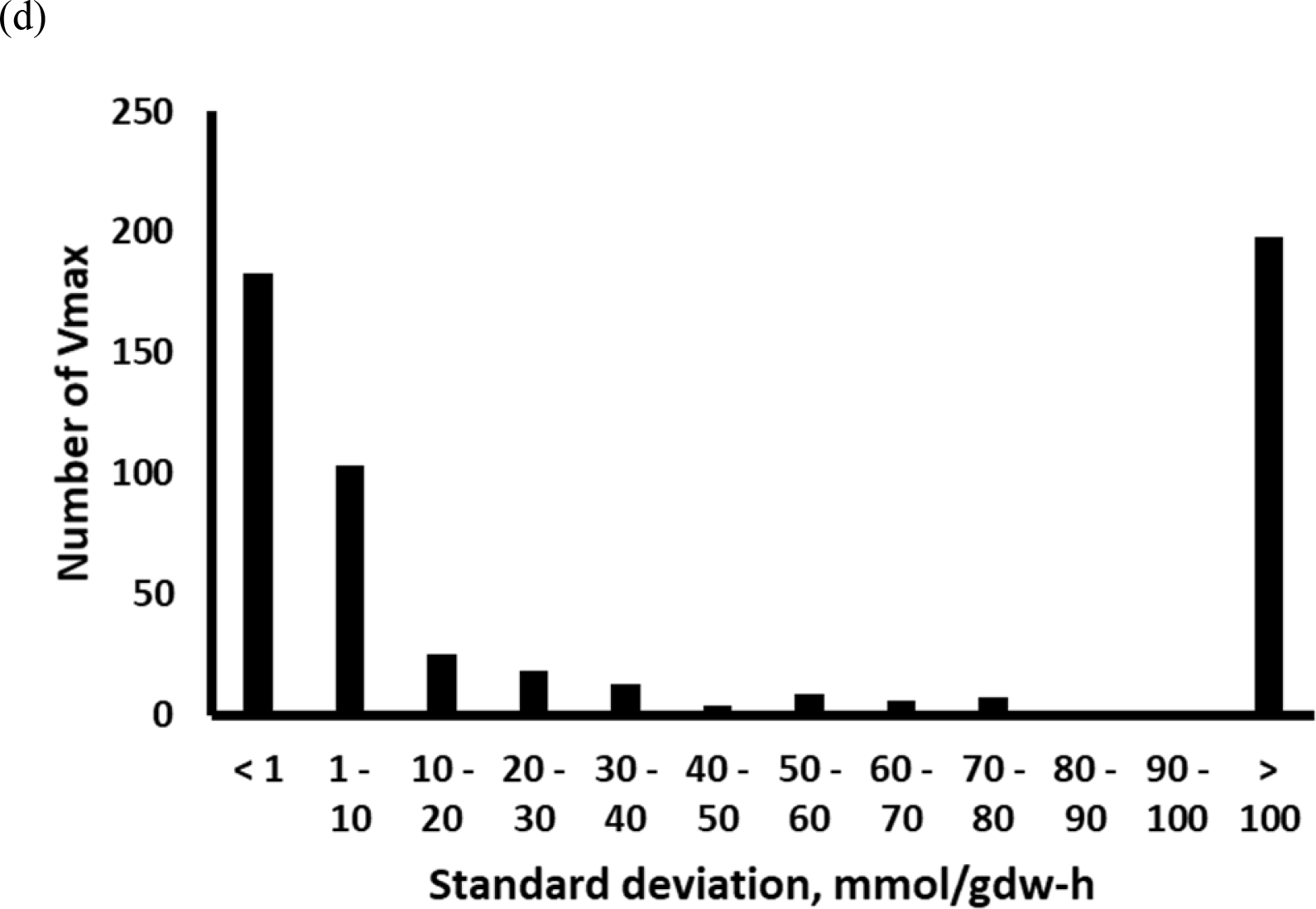
Distribution of standard deviations for estimated (a) WT enzyme fractions, (b) elementary kinetic parameters, (c) *K*_*m*_, and (d) *V*_*max*_ in k-ecoli307

**Figure S5:**
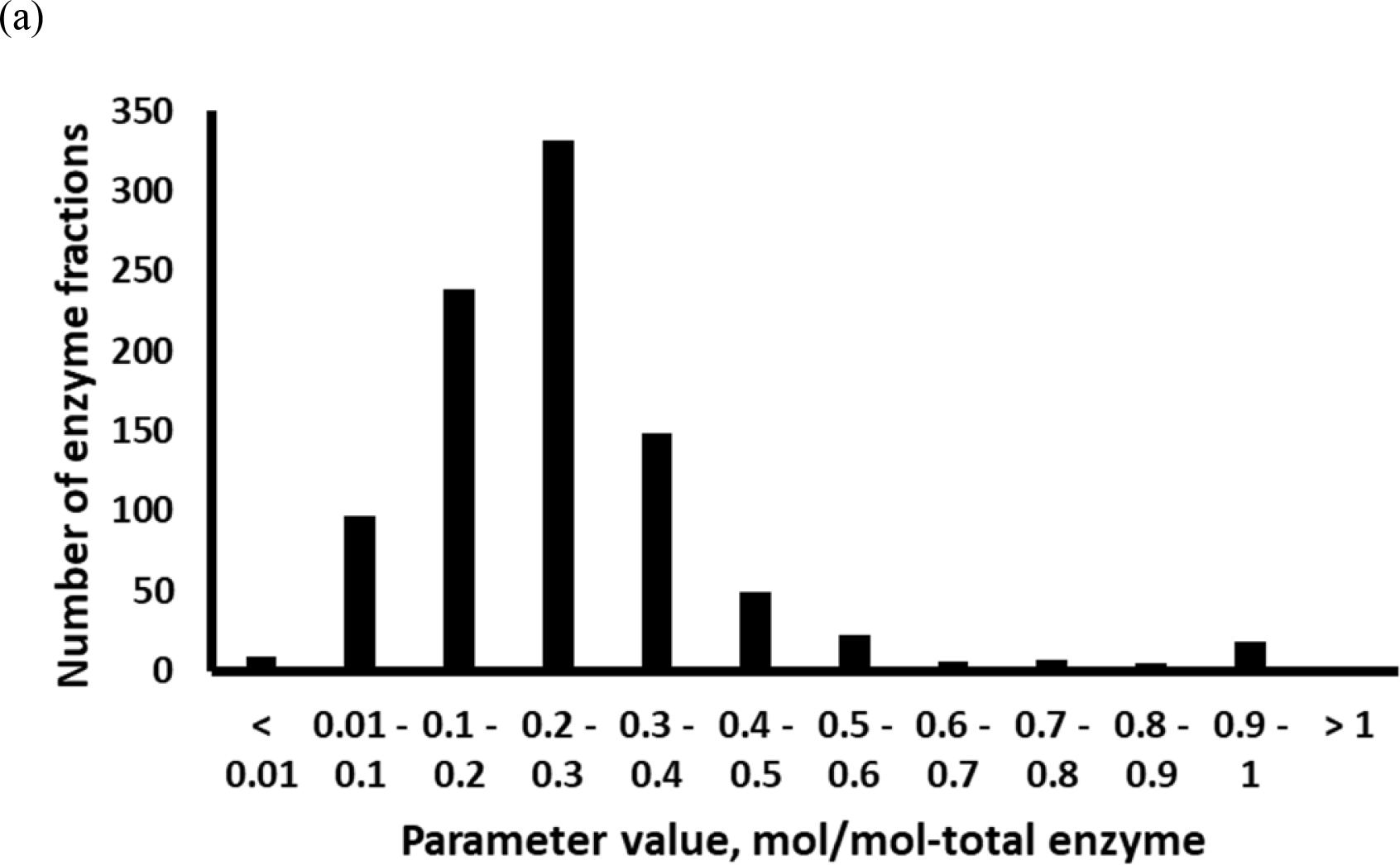
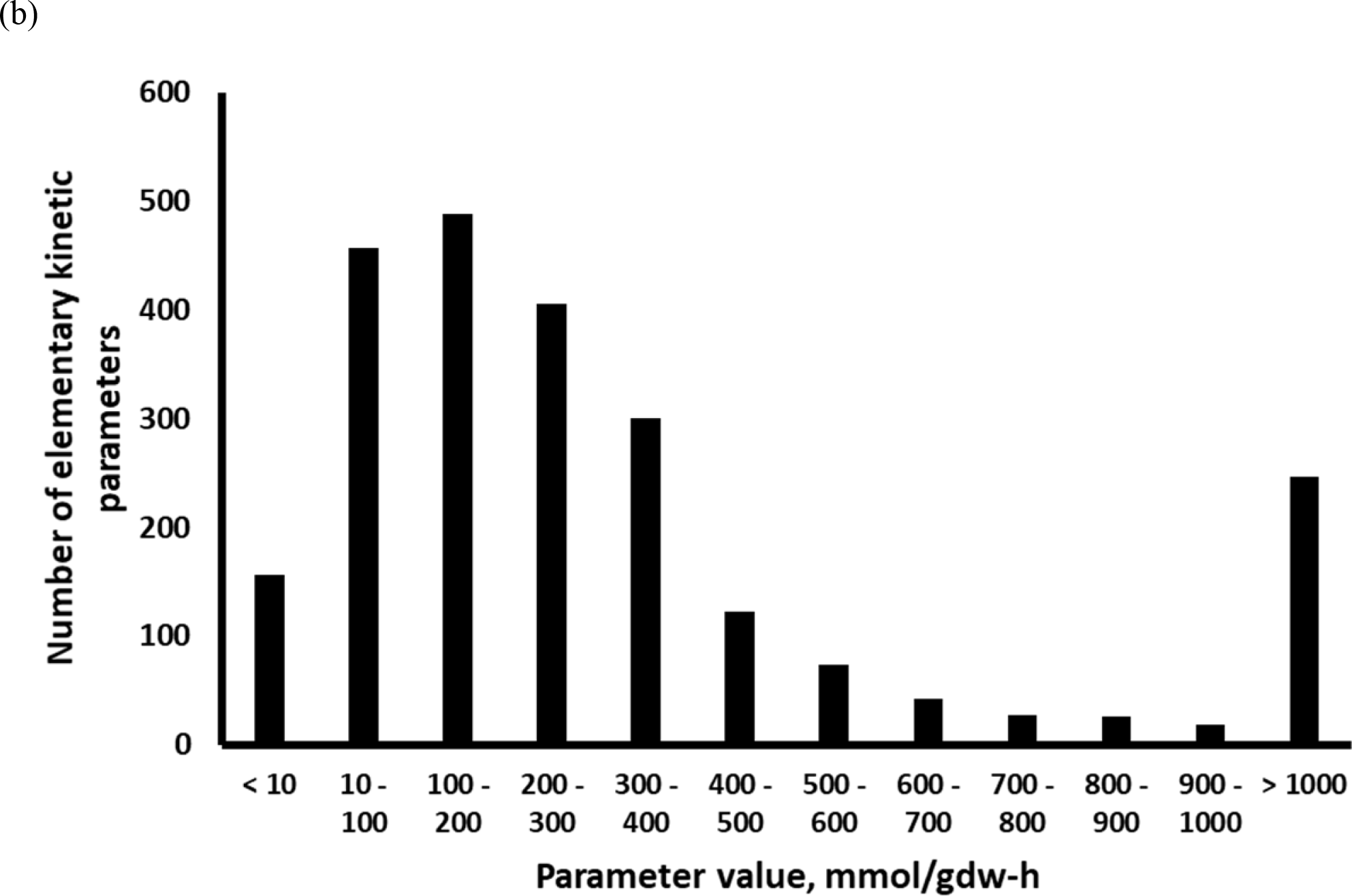
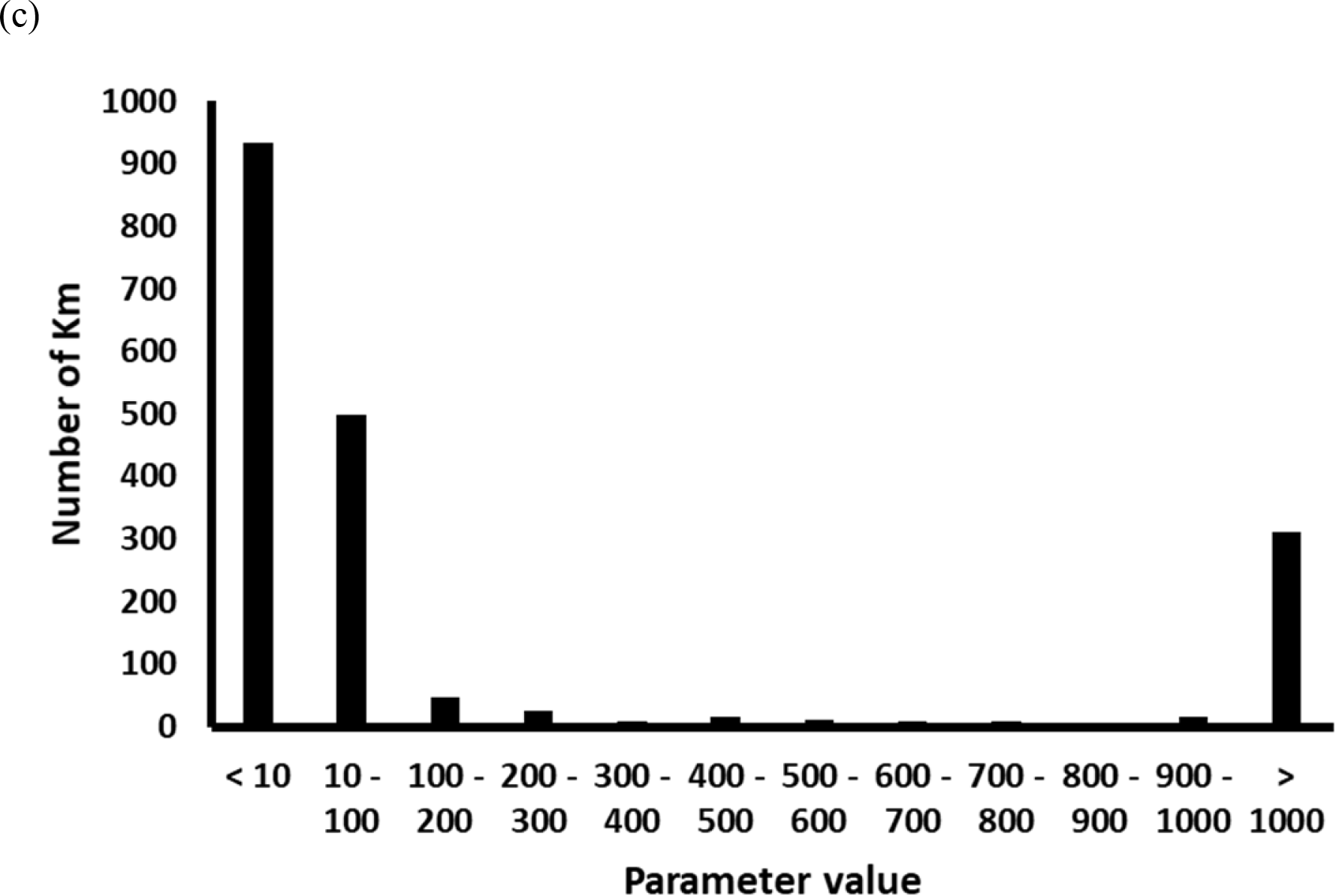
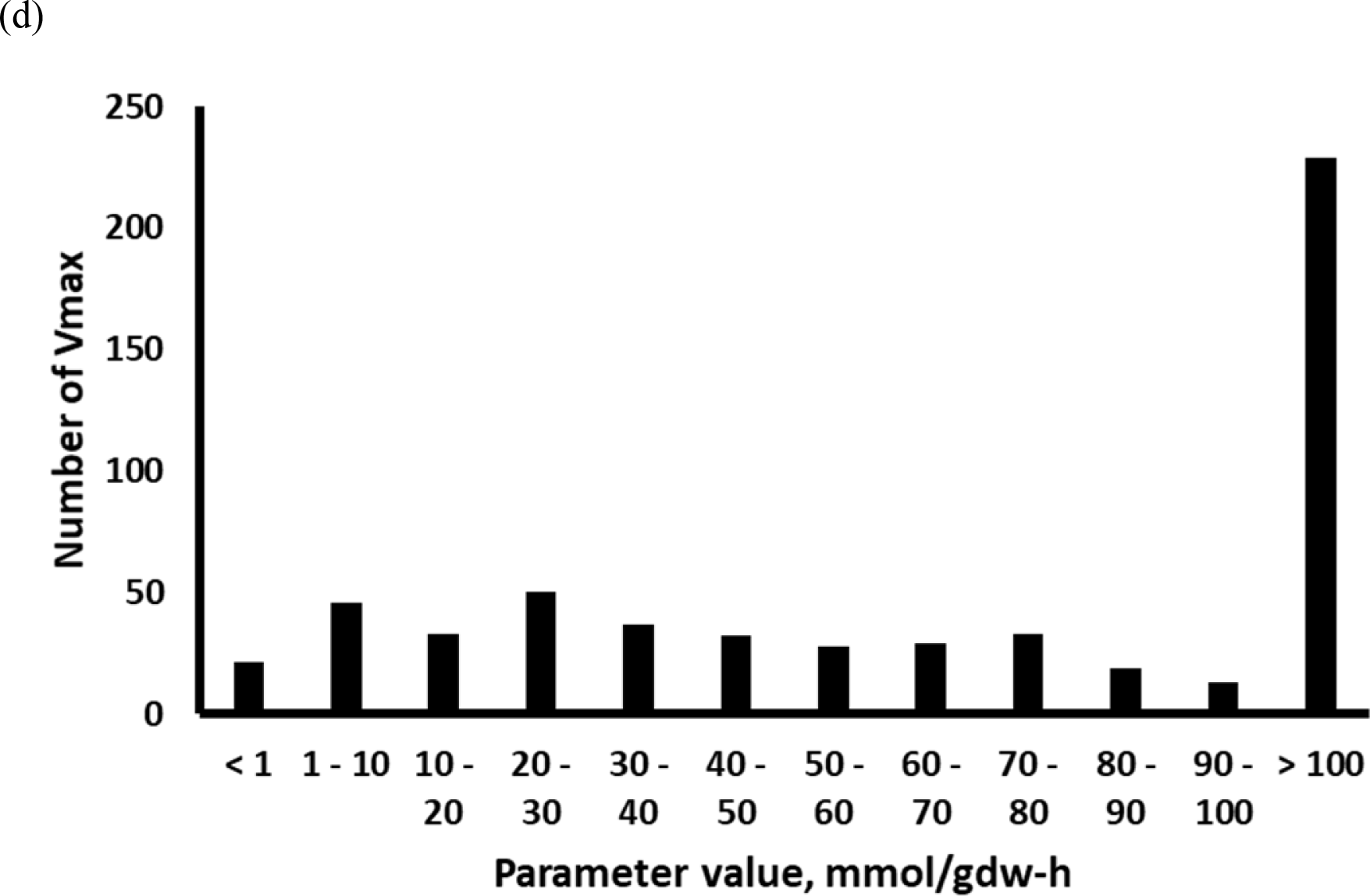
Distribution of values assumed by (a) WT enzyme fractions, (b) elementary kinetic parameters, (c) *K*_*m*_, and (d) *V*_*max*_ in k-ecoli307

