## Supplementary methods for "K-FIT: An accelerated kinetic parameterization algorithm using steady-state fluxomic data"

### Overview of elementary reaction step decomposition

The complex catalytic mechanism of enzyme catalysis can be decomposed into a series of *elementary steps* that are modeled using mass action kinetics. Each elementary step is treated as reversible with one forward and one reverse *elementary reaction*. Each elementary reaction is associated with one of three types of events: (i) binding of one metabolite with an enzyme complex, (ii) release of one metabolite from an enzyme complex, or (iii) conversion of the enzyme-reactant complex to the enzyme-product complex. The flux through each elementary reaction is termed *elementary flux* and is related to the concentration of metabolites and enzyme complexes using mass-action kinetics. The following example details the decomposition of an enzyme-catalyzed reaction into elementary steps and establishes the basic terms used in the kinetic parameterization algorithm.

Consider the conversion of a metabolite A to B catalyzed by an enzyme E, regulated by a non-competitive inhibitor C, and an activator D. The reaction mechanism can be decomposed into six elementary steps as shown in Table 1. The set of elementary steps $L$ is defined as $L = \{1,2,3,4,5,6\}$. Elementary steps 1, 2, and 3 describe the conversion of A to B and are therefore termed *catalytic elementary steps*. Elementary steps 4 and 5 model the inhibition of enzyme catalysis by metabolite C and step 6 denotes the activation of the inactive enzyme complex for catalysis by metabolite D. Steps 4, 5, and 6 do not participate in the reaction; instead they regulate enzyme function and are thus referred to as *regulatory elementary steps*. The set of catalytic elementary steps, denoted by $L^{cat}$, is defined here as $L^{cat} = \{1,2,3\}$. The corresponding set of regulatory elementary steps $L^{reg}$ is defined as $L^{reg} = \{4,5,6\}$. From Table 1, we see that the number of unique enzyme complexes formed over the course of the reaction is equal to the number of elementary steps required to model the catalytic and regulatory functions of the enzyme.

Each elementary step is modeled to be reversible with two separate elementary reactions in the forward and reverse directions. Thus, an enzyme-catalyzed reaction that decomposes into $n_L$ elementary steps will involve $n_P = 2n_L$ elementary reactions. The index of any elementary step $l \in L$ is related to the corresponding indices of its forward and reverse elementary reactions ($fwd$ and $rev$, respectively) as follows:

$$fwd = 2l - 1 \quad (1)$$
$$rev = 2l \quad (2)$$

| Type of Elementary Step | Elementary step # | Elementary Step | Description of elementary step |
|---|---|---|---|
| Catalytic Steps | 1 | $A + E \underset{\hat{k}_2}{\overset{\hat{k}_1}{\rightleftharpoons}} EA$ | Reactant binding to free enzyme |
| | 2 | $AE \underset{\hat{k}_4}{\overset{\hat{k}_3}{\rightleftharpoons}} EB$ | Conversion of reactant to product |
| | 3 | $EB \underset{\hat{k}_6}{\overset{\hat{k}_5}{\rightleftharpoons}} E + B$ | Product release from bound complex |
| Regulatory Steps | 4 | $C + E \underset{\hat{k}_8}{\overset{\hat{k}_7}{\rightleftharpoons}} EC$ | Inhibition of free enzyme |
| | 5 | $C + EA \underset{\hat{k}_{10}}{\overset{\hat{k}_9}{\rightleftharpoons}} EAC$ | Inhibition of enzyme-substrate complex |
| | 6 | $D + E^* \underset{\hat{k}_{12}}{\overset{\hat{k}_{11}}{\rightleftharpoons}} E$ | Activation of inactive enzyme form |

Table 1: List of elementary steps describing the catalytic mechanism and regulation of enzyme $E$

Based on this, the set of elementary reactions is defined as $P = \{p|p = 1,2, \ldots ,2L\}$. This implies that there is a sequence of alternating forward and reverse elementary reactions contained within set P. Each elementary step is associated with its own *kinetic rate constant* $\hat{k}_p \forall p \in P$. We define $[A]$, $[B]$, $[C]$, and $[D]$ to be the concentrations of metabolite $A$, $B$, $C$, and $D$, respectively, and $[E^*]$ $[E]$, $[EA]$ and $[EB]$ to denote the concentrations of the un-activated enzyme $E^*$, active free enzyme $E$, substrate-bound complex $EA$ and the product-bound complex $EB$, respectively. $[EC]$ and $[EAC]$ denote concentrations of the inhibitor-bound complexes $EC$ and $EAC$, respectively. As stated earlier, flux through elementary steps is referred to as *elementary flux*. For the example reaction, the elementary flux through the twelve elementary steps, $v_p$ $\forall p \in P$ can be computed by expressing the reaction rate of each elementary reaction using mass-action kinetics as described by Tran *et al* [1] and is shown in Equations (3):

$$\begin{aligned} v_1 &= \hat{k}_1[A][E] & v_2 &= \hat{k}_2[EA] \\ v_3 &= \hat{k}_3[EA] & v_4 &= \hat{k}_4[EB] \\ v_5 &= \hat{k}_5[EB] & v_6 &= \hat{k}_6[B][E] \\ v_7 &= \hat{k}_7[C][E] & v_8 &= \hat{k}_8[EC] \\ v_9 &= \hat{k}_9[C][EA] & v_{10} &= \hat{k}_{10}[EAC] \\ v_{11} &= \hat{k}_{11}[D][E^*] & v_{12} &= \hat{k}_{12}[E] \end{aligned} \tag{3}$$

Consistent with the convention introduced by Tran *et al* [1], the concentration of metabolites $A$, $B$, $C$, and $D$ are normalized with respect to the concentrations in the Wild-Type (WT) strain $A_{WT}$, $B_{WT}$, $C_{WT}$, and $D_{WT}$, respectively. The corresponding *relative concentrations* $a$, $b$, $c$, and $d$ are defined as:

$$\begin{aligned} a &= [A]/[A_{WT}] \\ b &= [B]/[B_{WT}] \\ c &= [C]/[C_{WT}] \\ d &= [D]/[D_{WT}] \end{aligned} \tag{4}$$

The total concentration $[E_0]$ of the enzyme catalyzing the conversion of $A$ to $B$ is related to the concentration of various enzyme forms/complexes as:

$$[E_0] = [E] + [EA] + [EB] + [EC] + [EAC] + [E^*] \tag{5}$$

*Enzyme fractions* are defined as the fractional abundance of each enzyme form relative to the total enzyme $[E_0]$.

$$\begin{aligned} e &= [E]/[E_0] & e_c &= [EC]/[E_0] \\ e_a &= [EA]/[E_0] & e_{ac} &= [EAC]/[E_0] \\ e_b &= [EB]/[E_0] & e^* &= [E^*]/E_0] \end{aligned} \tag{6}$$

Metabolite and total enzyme concentrations in the WT strain are often unavailable and are therefore, lumped together with kinetic rate constants yielding the following aggregated *kinetic parameters*:

$$\begin{aligned} k_1 &= \hat{k}_1[A_{WT}][E_0] & k_2 &= \hat{k}_2[E_0] \\ k_3 &= \hat{k}_3[E_0] & k_4 &= \hat{k}_4[E_0] \\ k_5 &= \hat{k}_5[E_0] & k_6 &= \hat{k}_6[B_{WT}][E_0] \\ k_7 &= \hat{k}_7[C_{WT}][E_0] & k_8 &= \hat{k}_8[E_0] \end{aligned} \tag{7}$$

$k_9 = \hat{k}_9[C_{WT}][E_0]$ $\qquad$ $k_{10} = \hat{k}_{10}[E_0]$
$k_{11} = \hat{k}_{11}[D_{WT}][E_0]$ $\qquad$ $k_{12} = \hat{k}_{12}[E_0]$

Upon substituting the definitions from Equations (4), (6) and (7) in Equation (3) expressions are derived for all fluxes as a function of aggregated kinetic parameters, relative metabolite concentrations and fractional enzyme abundances:

$$\begin{array}{ll} v_1 = k_1 a e & v_2 = k_2 e_a \\ v_3 = k_3 e_a & v_4 = k_4 e_b \\ v_5 = k_5 e_b & v_6 = k_6 b e \\ v_7 = k_7 c e & v_8 = k_8 e_c \\ v_9 = k_9 c e_a & v_{10} = k_{10} e_{ac} \\ v_{11} = k_{11} d e^* & v_{12} = k_{12} e \end{array} \tag{8}$$

Conservation of mass across all enzyme fractions at pseudo-steady-state yields the following linear equalities:

$$\frac{de}{dt} = v_2 + v_5 + v_8 + v_{11} - v_1 - v_6 - v_7 - v_{12} = 0 \tag{9}$$

$$\frac{de_a}{dt} = v_1 + v_4 + v_{10} - v_2 - v_3 - v_9 = 0 \tag{10}$$

$$\frac{de_b}{dt} = v_3 + v_6 - v_4 - v_5 = 0 \tag{11}$$

$$\frac{de_c}{dt} = v_7 - v_8 = 0 \tag{12}$$

$$\frac{de_{ac}}{dt} = v_9 - v_{10} = 0 \tag{13}$$

$$\frac{de^*}{dt} = v_{12} - v_{11} \tag{14}$$

Upon substituting the flux expressions from Equations (8) in Equations (9) - (14), an $[n_L \times n_L]$ square system of linear algebraic equations with the enzyme fractions as the only variables is obtained assuming that the relative metabolite concentrations ($a$, $b$, $c$, and $d$) and kinetic parameters ($k_p \forall p \in P$) are specified.

$$\frac{de}{dt} = k_2 e_a + k_5 e_a + k_8 e_c + k_{11} d e^* - (k_1 a + k_6 b + k_7 c + k_{12}) e = 0 \tag{15}$$

$$\frac{de_a}{dt} = k_1 a e + k_4 e_b + k_{10} e_{ac} - (k_2 + k_3 + k_9 c) e_a = 0 \tag{16}$$

$$\frac{de_b}{dt} = k_3 e_a + k_6 be - (k_4 + k_5)e_b = 0 \quad (17)$$

$$\frac{de_c}{dt} = k_7 ce - k_8 e_c = 0 \quad (18)$$

$$\frac{de_{ac}}{dt} = k_9 ce_a - k_{10} e_{ac} = 0 \quad (19)$$

$$\frac{de^*}{dt} = k_{12} e - k_{11} de^* = 0 \quad (20)$$

Note that Equation (15) can be reconstituted as a linear combination of Equations (16) - (20) because the free enzyme must be regenerated at the end of the catalytic cycle to maintain steady-state. This results in a rank-deficiency in this system of equations that can be rectified by appending Equation (21) which ensures that the total enzyme concentration is maintained constant at metabolic steady-state. Equation (21) is obtained by substituting Equations (6) in Equation (5):

$$e + e_a + e_b + e_c + e_{ac} = 1 \quad (21)$$

Equation (21) replaces Equation (15) resulting in an $[n_L \times n_L]$ system of equations of full-rank for computing enzyme fractions given kinetic parameters and relative metabolite concentrations. This means that given WT-normalized concentrations $a$, $b$, $c$, and $d$, and kinetic parameters $k_p \forall p \in P$, solving the system of linear equations yields a unique assignment for the enzyme fractions $e$, $e_a$, $e_b$, $e_c$, $e_{ac}$ and $e^*$. Fluxes through the elementary reactions are computed by substituting the newly computed enzyme fractions in Equations (8). Using the mapping of elementary flux indices to elementary step indices described in Equations (1) and (2), the net flux through any elementary step $l = \{1,2,3,4,5,6\}$ can be recovered as follows:

$$v_l^{(net)} = v_{2l-1} - v_{2l} \quad (22)$$

The net flux through all the catalytic steps ($l = \{1,2,3\}$) is equal to the net flux through the overall reaction $V$.

$$v_l^{(net)} = V \qquad l = \{1,2,3\} \quad (23)$$

From the steady-state conditions on the "dead-end" complexes formed via substrate-level regulation (see Equations (12), (13), and (14)), it can be derived that the net flux through the regulatory elementary steps is always equal to zero.

$$v_l^{(net)} = 0 \qquad l = \{4,5,6\} \qquad (24)$$

The automated calculation of the net flux through a reaction given elementary kinetic parameters and relative metabolite concentrations is facilitated by deriving generalized expressions for Equations (8) - (14) using the following quantities:

$\boldsymbol{v}$ is the $[n_P \times 1]$ vector of elementary fluxes whose elements $v_p$ denote the flux through elementary reaction $p \in P$

$\boldsymbol{e}$ is the $[n_L \times 1]$ vector of enzyme fractions whose elements $e_l$ represent the fractional abundance of enzyme complex $l \in L$

$I = \{i | i = 1,2, \ldots, n_M\}$ is the set of all metabolites. In the above example $n_M = 4$.

$\boldsymbol{s}$ is the $[n_M \times 1]$ matrix of relative metabolite concentrations whose elements $s_i$ represent the fold-change in concentration of metabolite $i \in I$ relative to WT.

$\boldsymbol{E}$ is the enzyme complex stoichiometry matrix of dimensions $[n_L \times n_P]$ whose elements $E_{lp}$ represent the stoichiometric coefficient of enzyme complex $l \in L$ in elementary reaction $p \in P$

$\boldsymbol{E}$ is defined as follows for the above example:

$$\boldsymbol{E} = \begin{array}{c} \\ \left[ \begin{array}{cccccccccccc} v_1 & v_2 & v_3 & v_4 & v_5 & v_6 & v_7 & v_8 & v_8 & v_{10} & v_{11} & v_{12} \\ -1 & 1 & 0 & 0 & 0 & 0 & -1 & 1 & 0 & 0 & 1 & -1 \\ 1 & -1 & -1 & 1 & 0 & 0 & 0 & 0 & -1 & 1 & 0 & 0 \\ 0 & 0 & 1 & -1 & -1 & 1 & 0 & 0 & 0 & 0 & 0 & 0 \\ 0 & 0 & 0 & 0 & 0 & 0 & 1 & -1 & 0 & 0 & 0 & 0 \\ 0 & 0 & 0 & 0 & 0 & 0 & 0 & 0 & 1 & -1 & 0 & 0 \\ 0 & 0 & 0 & 0 & 0 & 0 & 0 & 0 & 0 & 0 & -1 & 1 \end{array} \right] \end{array} \begin{array}{l} \\ e \\ e_a \\ e_b \\ e_c \\ e_{ac} \\ e^* \end{array}$$

Note that all the elements in $\boldsymbol{E}$ can assume only a value of -1, 0, or 1 and that $\boldsymbol{E}$ has exactly one negative and one positive entry per column. This is because, by definition, elementary reactions operate on a single enzyme form (either metabolite-bound of free) which is converted into another form but never destroyed. In contrast, the same enzyme form can participate in multiple elementary reactions and there exists at least one elementary reaction that consumes it (entry of -1) and at least one that produces it (i.e. entry of 1).

$\boldsymbol{S}$ is the metabolite stoichiometry matrix of dimensions $[n_M \times n_P]$ whose elements $S_{ip}$ represent the stoichiometric coefficient of metabolite $i \in I$ in elementary reaction $p \in P$

$\boldsymbol{S}$ is defined as follows for the above example:

$$\boldsymbol{S} = \begin{array}{c} \\ \left[\begin{array}{cccccccccccc} v_1 & v_2 & v_3 & v_4 & v_5 & v_6 & v_7 & v_8 & v_8 & v_{10} & v_{11} & v_{12} \\ -1 & 1 & 0 & 0 & 0 & 0 & 0 & 0 & 0 & 0 & 0 & 0 \\ 0 & 0 & 0 & 0 & 1 & -1 & 0 & 0 & 0 & 0 & 0 & 0 \\ 0 & 0 & 0 & 0 & 0 & 0 & -1 & 1 & -1 & 1 & 0 & 0 \\ 0 & 0 & 0 & 0 & 0 & 0 & 0 & 0 & 0 & 0 & -1 & 1 \end{array}\right] \begin{array}{c} \\ a \\ b \\ c \\ d \end{array} \end{array}$$

As was the case for matrix E, all elements of $\boldsymbol{S}$ are equal to either -1, 0 or 1. In addition, $\boldsymbol{S}$ has at most one non-zero entry per column. This is because an elementary reaction can represent only a single binding, release, or catalysis event [2]. Catalytic elementary reactions do not involve metabolites, whereas binding and release events either consume or produce a metabolite, respectively.

The flux $v_p$ through elementary reaction $p$ ($v_p$) is related to the concentration of metabolites and enzyme complexes using mass-action kinetics [3]:

$$v_p = k_p \left( \prod_{\substack{l \\ E_{lp}<0}} e_{l,c} \right) \left( \prod_{\substack{i \\ S_{ip}\leq 0}} s_i^{-S_{ip}} \right) \qquad \forall p \in P \qquad (25)$$

In Equation (25), the product operator in $\left( \prod_{\substack{l \\ E_{lp}<0}} e_l \right)$ serves to identify the only reactant enzyme complex participating in elementary reaction $p$. Recall that matrix E has a single element equal to -1 per column. Likewise, the product operator in $\left( \prod_{\substack{i \\ S_{ip}\leq 0}} s_i^{-S_{ip}} \right)$ serves to identify the only reactant metabolite (if any) in elementary reaction $p$. Recall that matrix $\boldsymbol{S}$ has at most one non-zero element per column equal to -1 or 1. Therefore, elementary reactions representing catalysis or product release do not involve a metabolite on the reactant side thus yielding a zero exponent. Elementary reactions modeling binding of a metabolite with an enzyme complex always involve a single reacting metabolite which yield an exponent of 1 (negative of -1 stoich. coeff.). This implies that in Equation (25) the exponent on the metabolite concentration is always equal to either 0 or 1. Equation (25) thus captures either a linear relation between $v_p$ and $e_l$ when there is no participating metabolite or a bilinear relation when a metabolite is a co-reactant in the elementary reaction. The kinetic parameter $k_p$ for elementary reaction $p \in P$ is a lumped parameter expressed as the product of the kinetic rate constant $\hat{k}_p$, the total enzyme concentration $E_0$, and the metabolite concentration in the WT as described by Equation (7).

Conservation of mass across the $l^{th}$ enzyme complex is mathematically represented as:

$$\frac{de_l}{dt} = \sum_{p=1}^{n_P} E_{lp} v_p \qquad \forall p \in P \qquad (26)$$

At pseudo-steady-state Equation (26) simplifies to:

$$\sum_{p=1}^{n_P} E_{lp} v_p = 0 \qquad \forall p \in P \qquad (27)$$

The net flux through the $l^{th}$ elementary step ($v_l^{(net)}$) is computed as the difference between the flux through the corresponding forward and reverse elementary reactions as described by Equation (22). The net flux through all catalytic elementary steps is equal to the net overall flux through the reaction. As a convention, we assign the "net" flux through the last catalytic elementary step as an index indicator of the net flux ($V$) through the overall reaction. This information is stored in the set $L^{(net)}$ which is $L^{(net)} = \{3\}$ for the above example. This index mapping the last catalytic step to the net flux through the overall reaction is accomplished using a $[1 \times n_L]$ indicator vector $\boldsymbol{N}$ whose elements are as follows:

$$N_j = \begin{cases} 1, & if\ l \in L^{(net)} \\ 0, & \text{Otherwise} \end{cases} \qquad (28)$$

In reference to the above example, $\boldsymbol{N}$ is a $[1 \times 6]$ vector ***vector would be [6 x 1] *** defined as $\boldsymbol{N} = [0 \quad 0 \quad 1 \quad 0 \quad 0 \quad 0]$. The net flux ($V$) through the overall reaction is recovered by the summation operator in Equation (29). Only a single term in the sum is non-zero.

$$V = \sum_{l=1}^{n_L} N_l v_l^{(net)} \qquad (29)$$

Even though the above treatment refers to a reversible uni-molecular reaction with non-competitive inhibition the same concepts can be generalized to any ordered or ping-pong mechanism of enzyme catalysis involving $n_{subs}$ substrates, $n_{pdt}$ products, activators, competitive inhibitors and uncompetitive inhibitors. Examples of elementary step decomposition for various reaction mechanisms is shown in Table 2. The above definitions and concepts form the foundation for the K-FIT procedure for estimating kinetic parameters given flux distributions.

Table 2: Elementary step decomposition for various reactions

| Reaction | Reaction Mechanism | Elementary Step Decomposition | Example from Central Metabolism |
|---|---|---|---|
| $A \rightleftharpoons B$ | Uni-Uni | $E + A \rightleftharpoons EA$<br>$EA \rightleftharpoons EB$<br>$EB \rightleftharpoons E + B$ | Phosphoglucose isomerase |
| $A \rightleftharpoons B + C$ | Uni-Bi | $E + A \rightleftharpoons EA$<br>$EA \rightleftharpoons EBC$<br>$EBC \rightleftharpoons EC + B$<br>$EC \rightleftharpoons E + C$ | Fructose bisphosphate aldolase |
| $A + B \rightleftharpoons C + D$ | Ordered Bi-Bi | $E + A \rightleftharpoons EA$<br>$EA + B \rightleftharpoons EAB$<br>$EAB \rightleftharpoons ECD$<br>$ECD \rightleftharpoons ED + C$<br>$ED \rightleftharpoons E + D$ | Phosphoglycerate kinase |
| $A + B \rightleftharpoons C + D$ | Bi-substrate Ping-Pong | $E + A \rightleftharpoons EA$<br>$EA \rightleftharpoons EC$<br>$EC \rightleftharpoons E^* + C$<br>$E^* + B \rightleftharpoons E^*B$<br>$E^*B \rightleftharpoons ED$<br>$ED \rightleftharpoons E + D$ | Transketolase |

**Nonlinear least-squares regression-based procedure for kinetic parameterization**

The K-FIT kinetic parameterization procedure is designed to make use of steady-state flux measurements for multiple genetic perturbations to parameterize a single kinetic model of metabolism. Kinetic parameters values are estimated by solving a least-squares problem that minimizes the deviation between predicted and experimentally measured steady-state flux distributions across all perturbed networks. The formal description of this least-squares optimization problem requires the definition of the following sets, parameters, variables and constraints:

**Sets**

Set of metabolites $I = \{i|i = 1,2, \dots, n_M\}$

Set of reactions $J = \{j|j = 1,2, \dots, n_R\}$

Set of elementary steps $L = \{l|l = 1,2, \dots, n_L\}$

$L_j^{cat} \subseteq L$ is the subset of all catalytic elementary steps for reaction $j$

$L_j^{reg} \subset L$ is the subset of all regulatory elementary steps for reaction $j$

Set of elementary reactions $P = \{p|p = 1,2, \dots, n_P\}$

Set of perturbation mutants $C = \{c|c = 1,2, \dots, n_C\}$ with $c = 1$ denoting wild-type WT (or reference) network

$J_c^{meas} \subseteq J$ is the subset of all reactions with available flux measurements under perturbation mutant $c \in C$. The cardinality of $J_c^{meas}$ is $n_c^{meas}$.

**Parameters**

$\boldsymbol{S}$ is the metabolite stoichiometry matrix of dimensions $[n_M \times n_P]$ whose elements $S_{ip}$ represent the stoichiometric coefficient of metabolite $i \in I$ in elementary reaction $p \in P$

$\boldsymbol{E}$ is the enzyme complex stoichiometry matrix of dimensions $[n_L \times n_P]$ whose elements $E_{lp}$ represent the stoichiometric coefficient of enzyme complex (or free enzyme) $l \in L$ in elementary reaction $p \in P$

$\boldsymbol{V}_c^{(meas)}$ is the $[n_c^{meas} \times 1]$ vector of flux measurements in mutant $c \in C$ whose elements $V_{j,c}^{(meas)}$ represent the measured flux through reaction $j \in J_c^{meas}$ with standard deviation $\sigma_{j,c}^{(meas)}$

$\boldsymbol{L}^{(net)}$ is the $[n_R \times 1]$ net flux mapping vector whose elements ($L_j^{(net)}$) store the index of the last catalytic elementary step $l \in L$ that quantifies the net flux through the overall reaction $j \in J$.

**Variables**

$\boldsymbol{k}$ is the $[n_P \times 1]$ vector of kinetic parameters whose elements $k_p$ denote the kinetic parameter for elementary reaction $p \in P$

$\boldsymbol{s}$ is the $[n_M \times n_C]$ matrix of relative metabolite concentrations whose elements $s_{ic}$ represent the fold-change in concentration of metabolite $i \in I$ in mutant $c \in C$ relative to WT. The $c^{th}$ column representing the $[n_M \times 1]$ vector of relative metabolite concentrations in mutant $c \in C$ is denoted as $\boldsymbol{s}_c$.

$\boldsymbol{e}$ is the $[n_L \times n_C]$ matrix of enzyme fractions whose elements $e_{lc}$ represent the fractional abundance of enzyme complex $l \in L$ in mutant $c \in C$. The $c^{th}$ column representing the $[n_L \times 1]$ vector of enzyme fractions for mutant $c \in C$ is denoted as $\boldsymbol{e}_c$. The number of enzyme complexes is equal to the number of elementary steps as discussed earlier.

$\boldsymbol{v}$ is the $[n_P \times n_C]$ matrix of elementary fluxes whose elements $v_{p,c}$ denote the flux through elementary reaction $p \in P$ in mutant $c \in C$. The $c^{th}$ column representing the $[n_p \times 1]$ vector of elementary fluxes in mutant $c \in C$ is denoted as $\boldsymbol{v}_c$.

$\boldsymbol{v}^{net}$ is the $[n_L \times n_C]$ matrix of net elementary fluxes whose elements $v_{l,c}^{net}$ represent the net flux through elementary step $l \in L$ in mutant $c \in C$. The $c^{th}$ column representing the $[n_L \times 1]$ vector of net elementary fluxes in mutant $c \in C$ is denoted as $\boldsymbol{v}_c^{net}$.

$\boldsymbol{V}$ is the $[n_R \times n_C]$ matrix of reaction fluxes whose elements $V_{j,c}$ denote the flux through reaction $j \in J$ in mutant $c \in C$. The $c^{th}$ column representing the $[n_R \times 1]$ vector of elementary fluxes in mutant $c \in C$ is denoted as $\boldsymbol{V}_c$.

In addition to these variable declarations the following three matrices are defined:

$\boldsymbol{R}$ is an $[n_R \times n_L]$ grouping matrix that indicates which enzyme complexes $l \in L$ participate in reaction $j \in J$. It is defined as:

$$R_{jl} = \begin{cases} 1 \text{ if } l \in \{L_j^{cat} \cup L_j^{reg}\} \\ 0, \text{otherwise} \end{cases}$$

$\boldsymbol{N}$ is an $[n_R \times n_L]$ indicator matrix that is used to map net flux through elementary steps $l$ to flux through the overall reaction $j \in J$. Based on the convention established in the Introduction section (Equation (28)), the last catalytic step serves as a measure of flux through the overall reaction. It is defined as:

$$N_{jl} = \begin{cases} 1\ if\ l = L_j^{(net)} \\ 0, \text{otherwise} \end{cases}$$

$\boldsymbol{Z}$ is an $[n_R \times n_C]$ indicator matrix that maps the abundance of the enzyme catalyzing reaction $j$ in mutant $c \in C$ relative to its abundance in the WT strain. It is defined as:

$$Z_{j,c} = \begin{cases} 0 \text{ if reaction } j \in J \text{ is eliminated under condition } c \in C \\ 1, \text{otherwise} \end{cases}$$

The definition of matrix $\boldsymbol{Z}$ implies that the mutant networks are derived by eliminating one or more reactions from the metabolic network of the reference strain. This definition can be generalized to incorporate other genetic perturbations such as over-expression and down-regulation of gene expression. In the absence of proteomic data in mutant strains, we assume that the enzymes maintain levels as in the WT.

**Least-squares minimization problem P1**

Using the definitions introduced above the least-squares minimization problem for kinetic parameterization is formulated for the general case as the following nonlinear optimization problem:

$$\min_{\boldsymbol{k},\boldsymbol{e},\boldsymbol{s},\boldsymbol{v},\boldsymbol{V}} \phi = \sum_{c=1}^{n_C} \sum_{j \in J_c^{meas}} \left( \frac{V_{jc} - V_{jc}^{(meas)}}{\sigma_{jc}} \right)^2$$

subject to:

$$v_{p,c} = k_p \left( \prod_{\substack{l \\ E_{lp}<0}} e_{l,c} \right) \left( \prod_{\substack{i \\ S_{ip}\le 0}} s_{i,c}^{-S_{ip}} \right) \qquad \forall p \in P,\ \forall c \in C \qquad (30)$$

$$v_{l,c}^{(net)} = v_{(2l-1),c} - v_{2l,c} \qquad \forall l \in L,\ \forall c \in C \qquad (31)$$

$$\sum_{p=1}^{P} (E_{lp} v_{pc}) = 0 \qquad \forall l \in L,\ \forall c \in C \qquad (32)$$

$$\sum_{l=1}^{n_L} (R_{jl} e_{l,c}) = Z_{jc} \qquad \forall j \in J,\ \forall c \in C \qquad (33)$$

$$\sum_{p=1}^{P} (S_{ip} v_{pc}) = 0 \qquad \forall i \in I,\ \forall c \in C \qquad (34)$$

$$V_{j,c} = \sum_{l=1}^{n_L} (N_{jl} v_{l,c}^{(net)}) \qquad \forall j \in J,\ \forall c \in C \qquad (35)$$

$$s_{i,c} \ge 0 \qquad \forall i \in I,\ \forall c \in C \qquad (36)$$

$$s_{i,1} = 1 \qquad \forall i \in I \qquad (37)$$

$$0 \le e_{l,c} \le 1 \qquad \forall l \in L,\ \forall c \in C \qquad (38)$$

$$k_p \ge 0 \qquad \forall p \in P \qquad (39)$$

Equation (30) in the above formulation represents the rate law for any elementary reaction governed by mass-action kinetics. It is a generalized form of Equation (25) accounting for reaction rates across all mutants $c \in C$. As discussed before, the role of the product operators is to select the single enzyme complex and (possibly) metabolite participating in the elementary reaction rate equation. Therefore, Equation (30) involves either a bilinear term (product of enzyme fraction times a relative metabolite concentration) or linear term (enzyme fraction term) in the right-hand side. Equation (32) and (34) enforce conservation of mass across all enzyme complexes and metabolites, respectively. Equation (32) is an extension of Equation (27) to include enzyme complex balances across all mutants. Equation (33) ensures that the total amount of the enzyme in all of its forms catalyzing reaction $j$ remains constant. It is a generalization of Equations (21) to account for enzyme presence or absence in different mutants. Thus, the presence or absence of reaction $j$ in mutant $c$ is captured by Equation (33). Equation (31) computes the net flux through any elementary step based on the mapping of elementary reactions to elementary steps established with Equations (1), (2) and (22). Equation (35) links the net flux through the elementary steps (i.e., last catalytic step) of a reaction to the overall flux through reaction $j$. Equation (38) ensures that the enzyme fractional abundances are bounded between zero and one. Equation (36) and (39) enforce non-negativity of relative metabolite concentrations and kinetic parameters, respectively. Since all metabolite concentrations are normalized with respect to the corresponding concentrations in the WT strain as described in Equations (4), Equation (37) sets all relative concentrations for the WT strain ($c = 1$) equal to one.

Equation (30) involving (at most) bilinear terms is the only set of nonlinear constraints in NLP problem P1. This constraint renders the optimization formulation nonconvex making even the identification of a feasible point challenging let alone convergence to the optimum value. Therefore, any attempt to solve problem P1 using an off-the-self NLP solver such MINOS [4], CONOPT [5], or *fmincon* from the Optimization Toolbox in MATLAB™ is unlikely to succeed due to difficulties in maintaining feasibility and progressively reduced step-length in the line-search. Conceptually, this can be remedied by integrating Equations (32) and (34) to steady-state after substituting the expression for elementary flux from Equation (30). However, this tends to be rather time consuming (i.e., order of minutes) due to the stiffness of the differential equations and the loss of accuracy arising from taking large time steps. Furthermore, the inability to integrate Equations (32) and (34) to steady-state for some sets of kinetic parameters results in the premature termination of any gradient-based optimization algorithm. Therefore, past efforts in kinetic parameterization have relied on meta-heuristic optimization algorithms such as Genetic Algorithm [6] and particle swarm optimization [7]. The lack of gradient information in this class of methods limits efficient traversal of the kinetic space in search of an acceptable solution which may or may not be optimal or even near-optimal for the least squares objective function. This computational inefficiency in performing kinetic model parameterization prevents any follow up calculations to assess uncertainties in kinetic parameters due to experimental errors or internal kinetic parameter dependencies. This computational inefficiency is one of the contributing factors that have so far throttled back the parameterization of large-scale and wide application of kinetic models in strain design. Faced with these challenges, we put forth a customized procedure that can reliably identify optimal or near optimal kinetic model parameterizations while achieve orders of magnitude improvement in computational time over stochastic approaches. The following subsections will describe strategies to transform problem P1 into a successive sequence of easier-to-solve subproblems. These strategies form the basis of the kinetic parameterization algorithm, K-FIT. K-FIT allows for the efficient solution of NLP problem P1 using three main tasks/procedures:

I. Procedure K-SOLVE anchors kinetic parameters $k_p$ to the specified steady-state flux distribution in the WT network $\boldsymbol{V}_1$ such that such that conservation of mass across metabolites (Equation (34)), pseudo-steady-state condition across all enzyme complexes (Equation (32)), and normalization of metabolite concentrations (Equation (37)) are simultaneously satisfied for the WT network. This is accomplished by rearranging Equation (30) to express $\boldsymbol{k}$ as a function of the WT enzyme fractions $\boldsymbol{e}_1$ and the flux through the reverse elementary reactions $\boldsymbol{v}_r \subset \boldsymbol{v}_1$ while maintaining the relative metabolite concentrations $s_{i,1} = 1 \, \forall i \in I$ (i.e., $\boldsymbol{k} = f(\boldsymbol{v}_r, \boldsymbol{e}_1)$).

II. Procedure SSF-Evaluator computes the steady-state fluxes $\boldsymbol{V}_c$ and relative metabolite concentrations $\boldsymbol{s}_c$ across all mutants ($c > 1$) using the kinetic parameters $\boldsymbol{k}$ computed in procedure K-SOLVE. Procedure SSF-Evaluator decomposes the system of bilinear equations in $\boldsymbol{s}_c$ and $\boldsymbol{e}_c$ defined by Equations (30), (32), (33), and (34) into two blocks of equations representing conservation of mass across enzyme complexes and metabolites, respectively. The bilinear equations become linear when one of either ($\boldsymbol{s}_c$ or $\boldsymbol{e}_c$) is specified. When $\boldsymbol{s}_c$ is specified, Equations (32) and (33) form an exactly determined $[n_L \times n_L]$ system of linear algebraic equations in $\boldsymbol{e}_c$. Similarly, Equation (34) represents an exactly determined $[n_M \times n_M]$ system of linear algebraic equations in $\boldsymbol{s}_c$ when $\boldsymbol{e}_c$ is specified. SSF-Evaluator iterates between these two blocks using originally a fixed-point iteration (FPI) scheme (or Newton / Richardson extrapolation if needed) until a steady-state is found. This strategy allows for the direct evaluation of both fluxes and concentration across all mutants that automatically satisfy all the nonlinear equality constraints from problem P1 and leaves only linear (in)equalities in the constraint set.

III. Procedure K-UPDATE computes the sensitivity of net flux through all reactions $\boldsymbol{V}$ to WT enzyme fractions $\boldsymbol{e}_1$ and reverse elementary fluxes $\boldsymbol{v}_r$, which is then used to compute the approximate gradient $\boldsymbol{G}$ and the approximate $\boldsymbol{H}$ for the objective function $\phi$. $\boldsymbol{G}$ and $\boldsymbol{H}$ are then used to check for optimality and update $\boldsymbol{e}_1$ and $\boldsymbol{v}_r$ using a Newton step if optimality is not achieved. The updated values for $\boldsymbol{e}_1$ and $\boldsymbol{v}_r$ are then fed to the K-SOLVE procedure which evaluates updated kinetic parameters $\boldsymbol{k}$ and the calculation sequence described above is repeated.

The K-FIT procedure is illustrated with Figure 1 and Supplementary Figure S1 which details the overall information flow loop, variable designations and calculation schema. The mathematical details and implementation of all the component subroutines of K-FIT are described in the following subsections.

### I. KSOLVE: Anchoring kinetic parameters to the WT flux distribution

K-SOLVE computes a set of kinetic parameters $\boldsymbol{k}$ that satisfy Equations (30) - (39) for the WT network ($c = 1$) when the WT flux distribution $\boldsymbol{V}_1$, enzyme fractions $\boldsymbol{e}_1$ and non-negative elementary fluxes $\boldsymbol{v}_1$ are specified. This anchoring is required because conservation of mass across all enzyme fractions, mass balance across metabolites, and normalization of metabolite concentrations may not be simultaneously satisfied. To demonstrate this, we recast Equations (32), (33), and (34) after substituting the expression for elementary fluxes in terms of mass-action kinetics described in Equation (30) and setting $s_{i,1} = 1 \forall i \in I$ based on Equation (37).

$$\sum_{l=1}^{n_L} (R_{jl} e_{l,1}) = 1 \qquad \begin{matrix} \forall l \in L \\ \forall c \in C \end{matrix} \qquad (40)$$

$$\sum_{p=1}^{n_P} \left( E_{lp} k_p \left( \prod_{\substack{l' \\ E_{l'p} < 0}} e_{l,c} \right) \right) = 0 \qquad \begin{matrix} \forall l \in L \\ \forall c \in C \\ \forall l' \in L \end{matrix} \qquad (41)$$

$$\sum_{p=1}^{n_P} \left( S_{ip} k_p \left( \prod_{\substack{l \\ E_{lp} < 0}} e_{l,c} \right) \right) = 0 \qquad \begin{matrix} \forall i \in I \\ \forall c \in C \end{matrix} \qquad (42)$$

Equations (40), (41), and (42) form an overdetermined system of $(n_L + n_M)$ linear algebraic equations in $n_L$ unknown enzyme fractions $\boldsymbol{e}_1$ when kinetic parameters $\boldsymbol{k}$ are specified. This system of equations for arbitrary values of $\boldsymbol{k}$ will likely be infeasible indicating that not possible values for kinetic parameters $\boldsymbol{k}$ simultaneously satisfy conservation of mass across all metabolites and enzyme complexes. This necessitates the development of the K-SOLVE procedure which derives a link between $\boldsymbol{k}$, and $\boldsymbol{e}_1$ so that conservation of mass is always satisfied. This is achieved by rearranging Equation (30) for the WT network and exploiting the property that the product term containing relative metabolite concentrations $\left( \prod_{\substack{i \\ S_{ip} \le 0}} s_{i,1}^{-S_{ip}} \right)$ will always be equal to one because the metabolite concentrations are scaled with respect to WT (i.e. $s_{i,1} = 1$):

$$k_p = v_{p,1} \left( \prod_{\substack{l \\ E_{lp} < 0}} e_{l,1} \right)^{-1} \qquad \forall p \in P \qquad (43)$$

Note that Equation (43) reveals that $k_p$ can be uniquely determined when both $\boldsymbol{v}_1$ and $\boldsymbol{e}_1$ are specified. Of these variables, $\boldsymbol{e}_1$ is bounded between 0 and 1, and further constrained by the following relations:

$$\sum_{l=1}^{n_L}(R_{jl}e_{l,1}) = 1 \qquad \forall l \in L \quad (40)$$

$$0 \leq e_{l,1} \leq 1 \qquad \forall l \in L \quad (44)$$

Equation (43) relates $2n_L$ kinetic parameters to $n_L$ enzyme fractions and $2n_L$ elementary fluxes. Enzyme fractions are further constrained by Equation (40) and bounded as shown in Equation (44) implying that there exists multiple value assignments for the $2n_L$ elementary fluxes that could yield the same $k_p$ values. This implies that the assignment of values for the elementary fluxes $v_p$ is not unique and that there exist unsatisfied degrees of freedom as only a subset of $v_p$ are independent variables. The reason for this dependency is the presence of pairs of forward and reverse fluxes that can assume an infinity of possibly combinations of values with the same net flux $v_l^{(net)}$ through the elementary step. We extract an independent subset of $\boldsymbol{v}_1$ by arbitrarily selecting the reverse flux as the independent variables and relating the forward fluxes as a function of the reverse and net fluxes. This requires the definition of two separate $[n_L \times 1]$ vectors $\boldsymbol{v}_f$ and $\boldsymbol{v}_r$ denoting fluxes through forward and reverse elementary reactions, respectively in the WT strain. Elements of vectors $\boldsymbol{v}_f$ and $\boldsymbol{v}_r$ are mapped to the $[2n_L \times 1]$ vector of elementary fluxes in the WT network ($\boldsymbol{v}_1$) using Equations (45) and (46).

$$v_{f,l} = v_{2l-1,1} \quad (45)$$

$$v_{r,l} = v_{2l,1} \quad (46)$$

Because the net flux through an elementary step $v_{l,1}^{(net)}$ is the difference between the forward and reverse elementary fluxes we obtain

$$v_{f_l} = v_{l,1}^{(net)} + v_{r_l} \qquad \forall l \in L \quad (47)$$

The net flux through all elementary steps of an enzyme-catalyzed reaction in the WT strain is related to the net flux through the reaction in the WT ($c = 1$) by Equations (48) and (49).

$$v_{l,1}^{(net)} = V_{j,1} \qquad \forall l \in L_j^{cat} \; \forall j \in J \quad (48)$$

$$v_{l,1}^{(net)} = 0 \qquad \forall l \in L_j^{reg} \; \forall j \in J \quad (49)$$

When $\boldsymbol{V}_1$ is specified then the values of the net fluxes $v_l^{(net)}$ through all elementary steps (both catalytic and regulatory) can be recovered from Equations (48) and (49). These values can then be plugged into Equation (47) to calculate $\boldsymbol{v}_f$ for a given assignment of value of the independent variables $\boldsymbol{v}_r$. Since vector $\boldsymbol{V}_1$ stores the steady-state fluxes in the WT, equality constraints representing conservation of mass across metabolites in the WT in problem P1 are inherently satisfied.

Therefore when $\boldsymbol{v}_1^{(net)}$, $\boldsymbol{v}_r$, and $\boldsymbol{e}_1$ are specified, a unique set of kinetic parameters $\boldsymbol{k}$ can be obtained by solving the following $[n_P \times n_P]$ system of linear algebraic equations.

$$v_{r,l} + v_l^{(net)} = k_{(2l-1)} \left( \prod_{\substack{l' \\ E_{l'p}<0}} e_{l',1} \right)$$

$$v_{r,l} = k_{(2l)} \left( \prod_{\substack{l' \\ E_{l'p}<0}} e_{l',1} \right) \qquad \forall p \in P \quad \forall l \in L \quad \forall l' \in L \qquad (50)$$

Note that the rate law expressions in Equations (50) are derived by setting the relative metabolite concentrations in Equation (30) for WT to one. The vector of kinetic parameters $\boldsymbol{k}$ is recovered from Equation (51) as the following explicit relations

$$k_{(2l-1)} = \left( v_{r_l} + v_l^{(net)} \right) \left( \prod_{\substack{l' \\ E_{l'p}<0}} e_{l',1} \right)^{-1}$$

$$k_{(2l)} = \left( v_{r_l} \right) \left( \prod_{\substack{l' \\ E_{l'p}<0}} e_{l',1} \right)^{-1} \qquad \forall p \in P \quad \forall l \in L \quad \forall l' \in L \qquad (51)$$

Values assumed by $\boldsymbol{e}_1$ and $\boldsymbol{v}_r$ are constrained by the following (in)equalities:

$$\sum_{l=1}^{n_L} R_{jl} e_{l,1} = 1 \qquad \forall j \in J \qquad (40)$$

$$0 \leq e_{l,1} \leq 1 \qquad \forall l \in L \qquad (44)$$

$$v_{r,l} \geq 0 \qquad \forall l \in L \qquad (52)$$

$$v_{r,l} + v_{l,1}^{(net)} \geq 0 \qquad \forall l \in L \qquad (53)$$

Since all elementary fluxes and enzyme fractions are non-negative, non-negativity of the kinetic parameters $\boldsymbol{k}$ computed in Equation (51) is always guaranteed. The steps for computing this feasible set of kinetic parameters is provided in the following algorithmic description. K-SOLVE accepts WT enzyme fractions $\boldsymbol{e}_1$ and reverse elementary fluxes $\boldsymbol{v}_r$ as inputs and returns kinetic parameters $\boldsymbol{k}$ as the output.

**Algorithm procedure K-SOLVE**

**Begin**

Specify and fix flux distribution in the WT strain $\boldsymbol{V}_1$.

Specify and fix $\boldsymbol{e}_1$ and $\boldsymbol{v}_r$ satisfying Equation (40), (44), (52), and (53).

Set $v_{l,1}^{(net)}\ \forall l \in L_j^{cat}$ to $V_{j,1} \forall j \in J$

Set $v_{l,1}^{(net)}\ \forall l \in L_j^{reg}$ to 0

Compute kinetic parameters $\boldsymbol{k}$ by substituting $\boldsymbol{e}_1$, $\boldsymbol{v}_r$, and $\boldsymbol{v}_1^{(net)}$ in Equation (51)

**return $\boldsymbol{k}$**

**end**

### II. SSF-Evaluator: Evaluation of steady-state fluxes for the mutant networks using the kinetic parameter assignments of K-SOLVE

Having computed a set of kinetic parameters $\boldsymbol{k}$ satisfying Equations (30) - (39) for the WT strain ($c = 1$) using K-SOLVE, the objective of SSF-Evaluator is to compute the flux distributions in the mutant strains. Typically, this is achieved by integrating the ODEs describing conservation of mass across all metabolites and enzyme complexes. To circumvent the unreliability and high computational cost associated with numerical integration, we put forth a decomposition-based approach that leverages the bilinear structure of the underlying system of equations. In this section, we derive updating formulae for the metabolite concentrations (Equations (57), (59), and (82)) in response to the altered enzyme concentrations compared to WT in the mutant networks (see Equation (33)). These update formulae are then fed into the SSF-Evaluator procedure that evaluates fluxes and metabolite concentrations in mutants when the kinetic parameters $\boldsymbol{k}$ are provided.

Substituting the expression for $v_{p,c}$ from Equation (30) that pose metabolite and enzyme mass balances as functions of enzyme fractions $\boldsymbol{e}_c$ and relative metabolite concentrations $\boldsymbol{s}_c$ across all mutant networks $c \in C$ into Equations (32) and (34) yields Equations (54) and (55), respectively:

$$\sum_{p=1}^{n_P}\left(E_{lp}k_p\left(\prod_{\substack{l' \\ E_{l'p}<0}} e_{l,c}\right)\left(\prod_{\substack{i \\ S_{ip}\le 0}} s_{i,c}^{-S_{ip}}\right)\right) = 0 \qquad \forall l \in L \quad \forall c \in C \qquad (54)$$

$$\sum_{p=1}^{n_P}\left(S_{ip}k_p\left(\prod_{\substack{l \\ E_{lp}<0}} e_{l,c}\right)\left(\prod_{\substack{i' \\ S_{i'p}\le 0}} s_{i,c}^{-S_{i'p}}\right)\right) = 0 \qquad \forall i \in I \quad \forall c \in C \qquad (55)$$

Equations (54) and (55) must be supplemented by Equation (33) that imposes that the sum of the fractional abundance of all enzyme complexes of a particular enzyme must be equal to the fold-change in the total enzyme level relative to WT. Thus, for every mutant network $c$, the enzyme fractions $\boldsymbol{e}_c$ encode any changes to enzyme level by means of upregulation, downregulation or absence as described by Equation (33).

$$\sum_{l=1}^{n_L}(R_{jl}e_{l,c}) = Z_{jc} \qquad \forall j \in J \quad \forall c \in C \qquad (33)$$

Equations (33) and (54) form a $[n_L \times n_L]$ system of linear algebraic equations of full rank in $\boldsymbol{e}_c$ that can efficiently be solved for the fractional enzyme complex abundances in all mutant networks given the values for the relative metabolite concentrations $\boldsymbol{s}_c$ and kinetic parameters $\boldsymbol{k}$ [8]. It is important to note that the steady-state enzyme fractions $\boldsymbol{e}_c$ encode any changes to enzyme presence in mutant $c$ through Equation (33).

Elementary binding and release steps bind only one reactant or release only one product at a time. This ensures that the only possible exponent for the metabolite concentration term in Equations (54) and (55) is equal to one. Equations (55) therefore simplifies to a system of linear algebraic equations in $\boldsymbol{s}_c$ when $\boldsymbol{e}_c$ is specified and can be recast as:

$$\sum_{\substack{p\in P\\ S_{ip}>0}}\left(S_{ip}k_p\left(\prod_{\substack{l\\ E_{lp}<0}} e_{l,c}\right)\right)+\sum_{\substack{p\in P\\ S_{ip}<0}}\left(S_{ip}k_p\left(\prod_{\substack{l\\ E_{lp}<0}} e_{l,c}\right)s_{i,c}\right)=0 \qquad \begin{matrix}\forall i\in I\\ \forall c\in C\end{matrix} \quad (56)$$

The relative metabolite concentrations can then be directly calculated from the following explicit expression:

$$s_{i,c}=-\frac{\sum_{\substack{p\in P\\ S_{ip}>0}}\left(S_{ip}k_p\left(\prod_{\substack{l\\ E_{lp}<0}} e_{l,c}\right)\right)}{\sum_{\substack{p\in P\\ S_{ip}<0}}\left(S_{ip}k_p\left(\prod_{\substack{l\\ E_{lp}<0}} e_{l,c}\right)\right)} \qquad \begin{matrix}\forall i\in I\\ \forall c\in C\end{matrix} \quad (57)$$

Equation (57) relates relative metabolite concentrations $\boldsymbol{s}_c$ to enzyme fractions $\boldsymbol{e}_c$ at metabolic steady-state for a given set of elementary step kinetic parameters $\boldsymbol{k}$. When $\boldsymbol{s}_c$ and $\boldsymbol{e}_c$ do not represent steady-state relative metabolite concentrations and enzyme fractions, the left hand-side of Equation (56) quantifies the mass imbalance of metabolite $i$ in network $c$ as shown in Equation (58).

$$\frac{ds_{i,c}}{dt}=\sum_{\substack{p\in P\\ S_{ip}>0}}\left(S_{ip}k_p\left(\prod_{\substack{l\\ E_{lp}<0}} e_{l,c}\right)\right)+\sum_{\substack{p\in P\\ S_{ip}<0}}\left(S_{ip}k_p\left(\prod_{\substack{l\\ E_{lp}<0}} e_{l,c}\right)s_{i,c}\right) \qquad \begin{matrix}\forall i\in I\\ \forall c\in C\end{matrix} \quad (58)$$

#### i. Fixed-Point Iteration (FPI)

In summary, the enzyme fractions $\boldsymbol{e}_c$ can be computed from the kinetic parameters $\boldsymbol{k}$ and metabolite concentrations $\boldsymbol{s}_c$ by solving the system of linear equations (33) and (54) and in turn the computed enzyme fractions $\boldsymbol{e}_c$ can be used to update metabolite concentrations $\boldsymbol{s}_c$. This establishes the following fixed-point iteration (FPI) procedure to solve for the unknown concentrations $\boldsymbol{s}_c$ and enzyme fractions $\boldsymbol{e}_c$ given kinetic parameters $\boldsymbol{k}$:

**Algorithmic Implementation of FPI**

**Begin**

Specify and fix $\boldsymbol{k}$

set $stol := 10^{-6}$, $iter := 1$

Initialize ${s_{i,c}}^{(0)} := 1, \forall i \in I$ and $c \in C$

Compute $\boldsymbol{e}_c^{(iter)}$ by solving Equations (33) and (54) with $\boldsymbol{s}_c := \boldsymbol{s}_c^{(0)}$

Compute $\boldsymbol{s}_c^{(iter)}$ by solving Equation (57) with $\boldsymbol{e}_c := \boldsymbol{e}_c^{(iter)}$

Compute $\frac{ds_c}{dt}$ by solving Equation (58) with $\boldsymbol{s}_c := \boldsymbol{s}_c^{(iter)}$ and $\boldsymbol{e}_c := \boldsymbol{e}_c^{(iter)}$

**While** $\left\|\frac{ds_c}{dt}\right\|_\infty > stol$ **or** $\left\|\boldsymbol{s}_c^{(iter+1)} - \boldsymbol{s}_c^{(iter)}\right\|_\infty > 10^{-4}$

$iter \coloneqq iter + 1$

Compute $\boldsymbol{e}_c^{(iter)}$ by solving Equations (33) and (54) with $\boldsymbol{s}_c := \boldsymbol{s}_c^{(iter)}$

Compute $\boldsymbol{s}_c^{(iter)}$ by solving Equation (57) with $\boldsymbol{e}_c := \boldsymbol{e}_c^{(iter)}$

Compute $\frac{ds_c}{dt}$ by solving Equation (58) with $\boldsymbol{s}_c := \boldsymbol{s}_c^{(iter)}$ and $\boldsymbol{e}_c := \boldsymbol{e}_c^{(iter)}$

**return** $\boldsymbol{s}_c^{(FPI)} \coloneqq \boldsymbol{s}_c^{(iter)}$ and $\boldsymbol{e}_c^{(FPI)} \coloneqq \boldsymbol{e}_c^{(iter)}$

**end**

It is important to note that the FPI algorithm has linear convergence which causes the method to slow down as we approach metabolic steady-state. This can be accelerated by switching to Newton's method which has quadratic convergence. We switch to Newton's method when either the mass imbalance is within the specified threshold of $stol$ or the progress towards steady-state becomes too slow. This happens when the change in metabolite concentrations between iterations falls below a pre-specified threshold of $10^{-4}$.

#### ii. Newton's method for accelerating convergence

Let $\boldsymbol{s}_c^{(FPI)}$ and $\boldsymbol{e}_c^{(FPI)}$ be current iterates that do not represent steady-state relative metabolite concentrations and enzyme fractions, respectively. They can be used as starting points for Newton's method where relative metabolite concentrations are updated in the $n^{th}$ iteration as:

$$\boldsymbol{s}_c^{(n+1)} = \boldsymbol{s}_c^{(n)} - \left( \frac{\partial \left( \frac{d\boldsymbol{s}_c}{dt} \right)}{\partial \boldsymbol{s}_c} \right)^{-1} \frac{d\boldsymbol{s}_c}{dt} \tag{59}$$

In Equation (59), $\frac{d\boldsymbol{s_c}}{dt}$ is computed using Equation (58). The quantity $\left(\frac{\partial\left(\frac{d\boldsymbol{s_c}}{dt}\right)}{\partial \boldsymbol{s_c}}\right)$ represents the Jacobian $\boldsymbol{J}$ of the function $\frac{d\boldsymbol{s_c}}{dt}$ described in Equation (59) and can be recast in terms of elementary fluxes as:

$$\frac{ds_{i,c}}{dt} = \sum_{p=1}^{P}\left(S_{ip}v_{pc}\right) = 0 \qquad \begin{matrix}\forall i \in I \\ \forall c \in C\end{matrix} \qquad (60)$$

The Jacobian $J$ obtained by differentiating Equation (60) with respect to $\boldsymbol{s_c}$ yields:

$$J_{ii\prime,c} = \frac{\partial}{\partial s_{i\prime,c}}\left(\frac{ds_{i,c}}{dt}\right) = \sum_{p=1}^{P} S_{ip}\left(\frac{\partial v_{pc}}{\partial s_{i',c}}\right) \qquad \begin{matrix}\forall i \in I \\ \forall c \in C \\ \forall i' \in I\end{matrix} \qquad (61)$$

Recall that $v_{pc}$ is related to kinetic parameters $\boldsymbol{k}$, enzyme fractions $\boldsymbol{e_c}$, and relative metabolite concentrations $\boldsymbol{s_c}$ using the mass-action kinetics of Equation (30). The sensitivity of $v_{pc}$ to the relative metabolite concentrations is obtained by differentiating Equation (30) with respect to $\boldsymbol{s_c}$:

$$v_{p,c} = k_p\left(\prod_{\substack{l \\ E_{lp}<0}} e_{l,c}\right)\left(\prod_{\substack{i \\ S_{ip}<0}} s_{i,c}^{-S_{ip}}\right) \qquad \begin{matrix}\forall p \in P \\ \forall c \in C\end{matrix} \qquad (30)$$

$$\frac{\partial v_{pc}}{\partial s_{i\prime,c}} = \sum_{\substack{l \\ E_{lp}<0}}\left(k_p\left(\left(\prod_{\substack{i \\ S_{ip}<0}} s_{q,c}^{-S_{ip}}\right)\frac{\partial}{\partial s_{i,c}}\left(\prod_{\substack{l \\ E_{lp}<0}} e_{l,c}\right) + \left(\prod_{\substack{l \\ E_{lp}<0}} e_{l,c}\right)\frac{\partial}{\partial s_{i\prime,c}}\left(\prod_{\substack{i \\ S_{ip}<0}} s_{q,c}^{-S_{ip}}\right)\right)\right) \qquad \begin{matrix}\forall p \in P \\ \forall c \in C \\ \forall i \in I \\ \forall i' \in I\end{matrix} \qquad (62)$$

Since only one enzyme complex and (at most) one metabolite participates in any elementary reaction, the derivatives in Equation (62) can be simplified as:

$$\frac{\partial}{\partial s_{i\prime,c}}\left(\prod_{\substack{l \\ E_{lp}<0}} e_{l,c}\right) = -\sum_{\substack{l \\ E_{lp}\leq 0}} E_{lp}\left(\frac{\partial e_{l,c}}{\partial s_{i',c}}\right) \qquad \begin{matrix}\forall i' \in I \\ \forall p \in P \\ \forall c \in C\end{matrix} \qquad (63)$$

$$\frac{\partial}{\partial s_{i\prime,c}}\left(\prod_{\substack{i \\ S_{ip}\leq 0}} s_{i,c}^{-S_{ip}}\right) = -\sum_{\substack{i \\ S_{ip}\leq 0}} S_{ip}\left(\frac{\partial s_{i,c}}{\partial s_{i',c}}\right) \qquad \begin{matrix}\forall i' \in I \\ \forall p \in P \\ \forall c \in C\end{matrix} \qquad (64)$$

Equation (62) can therefore be simplified by substituting the expressions for the derivatives in Equations (63) and (64) as:

$$\frac{\partial v_{pc}}{\partial s_{i',c}} = -k_p\left(\left(\prod_{\substack{i\\S_{ip}<0}} s_{i,c}^{-S_{ip}}\right)\left(\sum_{\substack{l\\E_{lp}\leq 0}} E_{lp}\frac{\partial e_{l,c}}{\partial s_{i',c}}\right)+\left(\prod_{\substack{l\\E_{lp}<0}} e_{l,c}\right)\left(\sum_{\substack{i\\S_{ip}\leq 0}} S_{ip}\frac{\partial s_{i,c}}{\partial s_{i',c}}\right)\right) \quad \begin{array}{l}\forall p\in P\\ \forall c\in C\\ \forall i'\in I\end{array} \tag{65}$$

The partial derivatives $\frac{\partial e_{l,c}}{\partial s_{i',c}}$ must be computed to quantify the sensitivity of elementary fluxes to substrate concentrations. This is achieved by differentiating Equations (33) and (54) with respect to $s_{i',c}$:

$$\sum_{l=1}^{n_L}\left(R_{jl}\frac{\partial e_{l,c}}{\partial s_{i',c}}\right) = 0 \quad \begin{array}{l}\forall l\in L\\ \forall c\in C\\ \forall i'\in I\end{array} \tag{66}$$

$$\sum_{p=1}^{P}\left(E_{lp}k_p\left(\prod_{\substack{i\\S_{ip}\leq 0}} s_{i,c}\right)\sum_{\substack{l\\E_{lp}<0}}\left(\frac{\partial e_{l,c}}{\partial s_{i',c}}\right)\right)-\sum_{p=1}^{P}\left(S_{i'p}E_{lp}k_p\left(\prod_{\substack{l\\E_{lp}<0}} e_{l,c}\right)\right) = 0 \quad \begin{array}{l}\forall l\in L\\ \forall c\in C\\ \forall i'\in I\end{array} \tag{67}$$

$\frac{\partial e_{l,c}}{\partial s_{i',c}}$ is computed by solving an exactly determined $[n_L \times n_L]$ system of linear algebraic equations formed by Equations (66) and (67). The computed $\frac{\partial e_{l,c}}{\partial s_{i',c}}$ is then substituted in Equation (65) to compute $\frac{\partial v_{pc}}{\partial s_{i,c}}$ which is subsequently substituted in Equation (61) to compute all elements in the Jacobian $\boldsymbol{J}$. Having computed $\boldsymbol{J}$, metabolite concentrations can be updated using Equation (59) until the steady-state concentrations are reached or $\boldsymbol{J}$ becomes singular. An alternative updating scheme for when $\boldsymbol{J}$ become singular is detailed in the following subsection. The following algorithm details the steps involved in the identification of steady-state metabolite concentrations using Newton's method.

**Algorithmic Implementation of Newton's Method**

**Begin**

Specify and fix $\boldsymbol{k}$

Set $stol := 10^{-6}$, $iter := 1$

Initialize $s_{i,c}{}^{(iter)} := s_{i,c}^{FPI}, \forall i \in I$ and $c \in C$

Compute $\boldsymbol{e}_c^{(iter)}$ by solving Equations (33) and (54) with $\boldsymbol{s}_c := \boldsymbol{s}_c^{(iter)}$

Compute $\frac{d\boldsymbol{s}_c}{dt}$ by substituting $\boldsymbol{s}_c := \boldsymbol{s}_c^{(iter)}$ and $\boldsymbol{e}_c := \boldsymbol{e}_c^{(iter)}$ into Equation (58)

Compute $\frac{\partial \boldsymbol{e}_c}{\partial \boldsymbol{s}}$ by solving Equations (66) and (67) with $\boldsymbol{s}_c := \boldsymbol{s}_c^{(iter)}$ and $\boldsymbol{e}_c := \boldsymbol{e}_c^{(iter)}$

Compute $\frac{\partial \boldsymbol{v}_c}{\partial \boldsymbol{s}}$ by substituting $\frac{\partial \boldsymbol{e}_c}{\partial \boldsymbol{s}}$ into Equation (65)

Compute $\boldsymbol{J}$ by substituting $\frac{\partial \boldsymbol{v}_c}{\partial \boldsymbol{s}}$ Equation (61)

**While** $\left(\left\|\frac{d\boldsymbol{s}_c}{dt}\right\|_\infty > stol\right)$ **and** $\boldsymbol{J}$ is not singular

$iter \coloneqq iter + 1$

Update $\boldsymbol{s}_c^{(iter)}$ by substituting $\frac{\partial\left(\frac{d\boldsymbol{s}_c}{dt}\right)}{\partial \boldsymbol{s}_c} = \boldsymbol{J}$ and $\boldsymbol{s}_c^{(n)} = \boldsymbol{s}_c^{(iter-1)}$ into Equation (59)

Compute $\boldsymbol{e}_c^{(iter)}$ by solving Equations (33) and (54) with $\boldsymbol{s}_c := \boldsymbol{s}_c^{(iter)}$

Compute $\frac{d\boldsymbol{s}_c}{dt}$ by substituting $\boldsymbol{s}_c := \boldsymbol{s}_c^{(iter)}$ and $\boldsymbol{e}_c := \boldsymbol{e}_c^{(iter)}$ into Equation (58)

Compute $\frac{\partial \boldsymbol{e}_c}{\partial \boldsymbol{s}}$ by solving Equations (66) and (67) with $\boldsymbol{s}_c := \boldsymbol{s}_c^{(iter)}$ and $\boldsymbol{e}_c := \boldsymbol{e}_c^{(iter)}$

Compute $\frac{\partial \boldsymbol{v}_c}{\partial \boldsymbol{s}}$ by substituting $\frac{\partial \boldsymbol{e}_c}{\partial \boldsymbol{s}}$ into Equation (65)

Compute $\boldsymbol{J}$ by substituting $\frac{\partial \boldsymbol{v}_c}{\partial \boldsymbol{s}}$ Equation (61)

**return** $\boldsymbol{s}_c^{(NM)} \coloneqq \boldsymbol{s}_c^{(iter)}$ and $\boldsymbol{e}_c^{(NM)} \coloneqq \boldsymbol{e}_c^{(iter)}$

**end**

On average we find $\boldsymbol{J}$ becomes singular in only approximately 5% of the all mutant flux evaluations using SSF-Evaluator, thus requiring a different updating formula.

#### iii. Richardson's Extrapolation when J becomes singular

If singularity for the Jacobian is detected then we switch to a semi-implicit first-order integrator [9] using Richardson's extrapolation by initializing the relative metabolite concentrations at the current point ($\boldsymbol{s}_c^{(NM)}$). The update formula for the metabolite concentrations (Equation (72)) is derived using the following procedure. The initial value problem described by Equation (60) can be expressed in matrix form as:

$$\frac{d\boldsymbol{s}_c}{dt} = \boldsymbol{S}.\boldsymbol{v}_c = \boldsymbol{f}(\boldsymbol{s}_c) \qquad \forall c \in C \qquad (68)$$

Equation (68) is integrated starting from the initial condition $\boldsymbol{s}(0) = \boldsymbol{s}_c^{(NM)}$ where $\boldsymbol{s}_c^{(NM)}$ is the vector of relative metabolite concentrations when Newton's method fails (J becomes singular). We use the implicit Euler's method to update substrate concentrations $\boldsymbol{s}$ upon taking a time step of $h$. This is due to the stiffness of the system of equations that precludes the use of a less costly explicit method. The update formula for the $n^{th}$ iteration is:

$$\frac{\boldsymbol{s}_c^{(n+1)}-\boldsymbol{s}_c^{(n)}}{h} = \boldsymbol{f}(\boldsymbol{s}_c^{(n+1)}) \qquad \forall c \in C \qquad (69)$$

Since $\boldsymbol{s}_c^{(n+1)}$ is unknown, $\boldsymbol{f}(\boldsymbol{s}_c^{(n+1)})$ cannot be evaluated *a priori* and must be approximated using Taylor series expansion.

$$\boldsymbol{f}\left(\boldsymbol{s}_c^{(n+1)}\right) = \boldsymbol{f}\left(\boldsymbol{s}_c^{(n)}\right) + \frac{\partial \boldsymbol{f}}{\partial \boldsymbol{s}_c}\left(\boldsymbol{s}_c^{(n+1)} - \boldsymbol{s}_c^{(n)}\right) \qquad \forall c \in C \qquad (70)$$

Equation (70) is substituted back in Equation (69) to yield:

$$\boldsymbol{s}_c^{(n+1)} - \boldsymbol{s}_c^{(n)} = h\boldsymbol{f}\left(\boldsymbol{s}_c^{(n)}\right) + h\frac{\partial \boldsymbol{f}}{\partial \boldsymbol{s}}\left(\boldsymbol{s}_c^{(n+1)} - \boldsymbol{s}_c^{(n)}\right) \qquad \forall c \in C \qquad (71)$$

Equation (71) is rearranged to obtain the semi-implicit update formula for $\boldsymbol{s}_c$ :

$$\boldsymbol{s}_c^{(n+1)} = \boldsymbol{s}_c^{(n)} + \left(\boldsymbol{I} - h\frac{\partial \boldsymbol{f}}{\partial \boldsymbol{s}_c}\right)^{-1} h\boldsymbol{f}\left(\boldsymbol{s}_c^{(n)}\right) \qquad \forall c \in C \qquad (72)$$

$\frac{\partial \boldsymbol{f}}{\partial \boldsymbol{s}_c}$ in Equation (72) is the Jacobian matrix $\boldsymbol{J}$ also present in Equation (61) and is calculated as described earlier. Equation (72) is integrated using the error-controlled integration algorithm Richardson extrapolation until either the time step $h$ exceeds a maximum time step of $h_{max}$ or the desired threshold on Equation (68) is reached ($\left\|\frac{d\boldsymbol{s}_c}{dt}\right\|_\infty \leq stol$). If $h$ exceeds $h_{max}$, Newton's method is reinitialized using concentrations at the termination point of the semi-implicit integration procedure ($\boldsymbol{s}_c^{(INT)}$) and solved until $\left\|\frac{d\boldsymbol{s}_c}{dt}\right\|_\infty \leq stol$ is achieved.

**Algorithmic Implementation of Semi-implicit integration using Richardson's extrapolation**

**Begin**

Specify and fix $\boldsymbol{k}$

Set $stol := 10^{-6}$, $iter := 1$, $h \coloneqq 2 \times 10^{-6}$, $h_{max} \coloneqq 10^{10}$, $tol \coloneqq 10^{-4}$

Initialize $s_{i,c}{}^{(iter)} := s_{i,c}^{(NM)}, \forall i \in I$ and $c \in C$

Compute $\boldsymbol{e}_c^{(iter)}$ by solving Equations (33) and (54) with $\boldsymbol{s}_c := \boldsymbol{s}_c^{(iter)}$

Compute $\frac{d\boldsymbol{s}_c}{dt}$ by substituting $\boldsymbol{s}_c := \boldsymbol{s}_c^{(iter)}$ and $\boldsymbol{e}_c := \boldsymbol{e}_c^{(iter)}$ into Equation (58)

Compute $\boldsymbol{J}$ by solving Equations (61), (65), (66) and (67) with $\boldsymbol{s}_c := \boldsymbol{s}_c^{(iter)}$ and $\boldsymbol{e}_c := \boldsymbol{e}_c^{(iter)}$

Set $iter \coloneqq iter + 1$

**While** $\left(\left\|\frac{ds_c}{dt}\right\|_\infty > stol\right)$ **or** $h < h_{max}$

Compute $\boldsymbol{s}_c^{(n)}$ by substituting $\boldsymbol{s}_c^{(n-1)} := \boldsymbol{s}_c^{(iter-1)}$, $\boldsymbol{f}\left(\boldsymbol{s}_c^{(n-1)}\right) \coloneqq \frac{d\boldsymbol{s}_c}{dt}$, $\frac{\partial \boldsymbol{f}}{\partial \boldsymbol{s}_c} \coloneqq \boldsymbol{J}$, and $h \coloneqq h$ into Equation (72)

Set $\boldsymbol{s}_c^{(one-step)} \coloneqq \boldsymbol{s}_c^{(n)}$

Compute $\boldsymbol{s}_c^{(n)}$ by substituting $\boldsymbol{s}_c^{(n-1)} := \boldsymbol{s}_c^{(iter-1)}$, $\boldsymbol{f}\left(\boldsymbol{s}_c^{(n-1)}\right) \coloneqq \frac{d\boldsymbol{s}_c}{dt}$, $\frac{\partial \boldsymbol{f}}{\partial \boldsymbol{s}_c} \coloneqq \boldsymbol{J}$, and $h \coloneqq \frac{h}{2}$ into Equation (72).

Compute $\boldsymbol{e}_c^{(iter)}$ by solving Equations (33) and (54) with $\boldsymbol{s}_c := \boldsymbol{s}_c^{(n)}$

Compute $\frac{d\boldsymbol{s}_c}{dt}$ by substituting $\boldsymbol{s}_c := \boldsymbol{s}_c^{(n)}$ and $\boldsymbol{e}_c := \boldsymbol{e}_c^{(iter)}$ into Equation (58).

Compute $\boldsymbol{J}$ by solving Equations (61), (65), (66) and (67) with $\boldsymbol{s}_c := \boldsymbol{s}_c^{(n)}$ and $\boldsymbol{e}_c := \boldsymbol{e}_c^{(iter)}$

Compute $\boldsymbol{s}_c^{(n)}$ by substituting $\boldsymbol{s}_c^{(n-1)} := \boldsymbol{s}_c^{(n)}$, $\boldsymbol{f}\left(\boldsymbol{s}_c^{(n-1)}\right) \coloneqq \frac{d\boldsymbol{s}_c}{dt}$, $\frac{\partial \boldsymbol{f}}{\partial \boldsymbol{s}_c} \coloneqq \boldsymbol{J}$, and $h \coloneqq \frac{h}{2}$ into Equation (72).

Set $\boldsymbol{s}_c^{(two-step)} \coloneqq \boldsymbol{s}_c^{(n)}$

**if** $\left(\left\|\boldsymbol{s}_c^{(two-step)} - \boldsymbol{s}_c^{(one-step)}\right\|_\infty < tol\right)$

Set $\boldsymbol{s}_c^{(iter)} \coloneqq \boldsymbol{s}_c^{(two-step)}$

Compute $\boldsymbol{e}_c^{(iter)}$ by solving Equations (33) and (54) with $\boldsymbol{s}_c := \boldsymbol{s}_c^{(iter)}$

Compute $\frac{d\boldsymbol{s}_c}{dt}$ by substituting $\boldsymbol{s}_c := \boldsymbol{s}_c^{(iter)}$ and $\boldsymbol{e} := \boldsymbol{e}_c^{(iter)}$ into Equation (58).

Compute $\boldsymbol{J}$ by solving Equations (61), (65), (66) and (67) with $\boldsymbol{s}_c := \boldsymbol{s}_c^{(iter)}$ and $\boldsymbol{e} := \boldsymbol{e}_c^{(iter)}$

Set $h \coloneqq \frac{h \times \sqrt{tol}}{\sqrt{\left\|\boldsymbol{s}^{(two-step)} - \boldsymbol{s}^{(one-step)}\right\|_\infty}}$

Set $iter \coloneqq iter + 1$

**else**

Set $h \coloneqq \frac{h}{2}$

**return** $\boldsymbol{s}_c^{(INT)} \coloneqq \boldsymbol{s}_c^{(iter)}$ and $\boldsymbol{e}_c^{(INT)} \coloneqq \boldsymbol{e}_c^{(iter)}$

**end**

For the large-scale kinetic model (k-ecoli307) parameterized in this study (see Results section in the main manuscript), the average computation time required to evaluate steady-state fluxes in mutants by FPI, Newton's method, and semi-implicit integration was 10 seconds, 4 seconds, and 37 seconds, respectively. In contrast, steady-state flux evaluation using numerical integration alone required over 6 minutes to achieve the same mass imbalance of $10^{-3}$ mol%. CPU times are reported are reported for an Intel-i7 (4-core processor, 2.6GHz, 12GB RAM) computer using a single core implementation.

The three separate methods of updating metabolite concentrations (i) FPI, (ii) Newton's method and (iii) semi-implicit integration and switching criteria are integrated into the SSF-Evaluator procedure. SSF-Evaluator initially solves for steady-state concentrations using FPI and switches to Newton's method when the change in metabolite concentrations between successive iterations falls below a pre-specified threshold of $10^{-4}$. Newton's method fails when the Jacobian $\boldsymbol{J}$ becomes singular, which prompts the switch to semi-implicit integration using Richardson's extrapolation. The following summarizes in detail the algorithmic steps involved:

**Algorithmic Implementation of Steady-State Flux Estimator (SSF-Evaluator)**

**Begin**

Specify and fix $\boldsymbol{k}$

Specify $mutant\ c$

Set $stol := 10^{-6}$

Initialize ${s_{i,c}}^{(init)} := 1, \forall i \in I$

Compute $\boldsymbol{e}_c^{(init)}$ by solving Equations (33) and (54)with $\boldsymbol{s}_c := \boldsymbol{s}_c^{(init)}$

Compute $\frac{ds_c}{dt}$ by substituting $\boldsymbol{s}_c := \boldsymbol{s}_c^{(init)}$ and $\boldsymbol{e}_c := \boldsymbol{e}_c^{(init)}$ into Equation (58).

**While** $\left(\left\|\frac{ds_c}{dt}\right\|_\infty > stol\right)$

Compute $\boldsymbol{s}_c^{(FPI)}$ by using the **FPI** algorithm using $\boldsymbol{s}_c^{(0)} = \boldsymbol{s}_c^{(init)}$

Compute $\boldsymbol{e}_c^{(FPI)}$ by solving Equations (33) and (54) with $\boldsymbol{s}_c := \boldsymbol{s}_c^{(FPI)}$

Compute $\frac{ds_c}{dt}$ by substituting $\boldsymbol{s}_c := \boldsymbol{s}^{(FPI)}$ and $\boldsymbol{e}_c := \boldsymbol{e}^{(FPI)}$ into Equation (58).

**if** $\left(\left\|\frac{ds_c}{dt}\right\|_\infty \le stol\right)$

Set $\boldsymbol{s}_c^{(SS)} \coloneqq \boldsymbol{s}_c^{(FPI)}$

**else**

Set $\boldsymbol{s}_c^{(INT)} \coloneqq \boldsymbol{s}_c^{(FPI)}$

**while** $\left(\left\|\frac{ds_c}{dt}\right\|_\infty > stol\right)$

Compute $\boldsymbol{s}_c^{(NM)}$ by solving the **Newton's method** using $\boldsymbol{s}_c^{(0)} = \boldsymbol{s}_c^{(INT)}$

Compute $\boldsymbol{e}_c^{(NM)}$ by solving Equations (33) and (54) with $\boldsymbol{s}_c := \boldsymbol{s}^{(NM)}$

Compute $\frac{d\boldsymbol{s}_c}{dt}$ by substituting $\boldsymbol{s}_c := \boldsymbol{s}^{(NM)}$ and $\boldsymbol{e}_c := \boldsymbol{e}^{(NM)}$ into Equation (58).

**if** $\left(\left\|\frac{d\boldsymbol{s}_c}{dt}\right\|_\infty > stol\right)$

Compute $\boldsymbol{s}_c^{(INT)}$ using **Semi-implicit integration** using $\boldsymbol{s}_c^{(0)} = \boldsymbol{s}_c^{(NM)}$

Compute $\boldsymbol{e}_c^{(INT)}$ by solving Equations (33) and (54) with $\boldsymbol{s}_c := \boldsymbol{s}^{(INT)}$

Compute $\frac{d\boldsymbol{s}_c}{dt}$ by substituting $\boldsymbol{s}_c := \boldsymbol{s}^{(INT)}$ and $\boldsymbol{e}_c := \boldsymbol{e}^{(INT)}$ into Equation (58).

**if** $\left(\left\|\frac{d\boldsymbol{s}_c}{dt}\right\|_\infty \leq stol\right)$

Set $\boldsymbol{s}_c^{(SS)} \coloneqq \boldsymbol{s}_c^{(INT)}$

**else**

Set $\boldsymbol{s}_c^{(SS)} \coloneqq \boldsymbol{s}_c^{(NM)}$

Compute $\boldsymbol{e}_c^{(SS)}$ by solving Equations (33) and (54) with $\boldsymbol{s}_c := \boldsymbol{s}_c^{(SS)}$

Compute steady-state fluxes $\boldsymbol{V}_c^{(SS)}$ by solving Equations (30), (31), and (35) with $\boldsymbol{s}_c := \boldsymbol{s}_c^{(SS)}$ and $\boldsymbol{e}_c := \boldsymbol{e}_c^{(SS)}$

**return** $\boldsymbol{V}_c^{(SS)}$, $\boldsymbol{s}_c^{(SS)}$ and $\boldsymbol{e}_c^{(SS)}$

**end**

Overall, SSF-Evaluator provides an integrated procedure for calculating steady-state relative metabolite concentrations and enzyme fractions across all mutant networks given a set of kinetic parameters bypassing integration in almost all cases. Steady-state elementary fluxes are then computed by substituting the known $\boldsymbol{k}$, $\boldsymbol{s}^{(SS)}$ and $\boldsymbol{e}^{(SS)}$ into Equation (30). Elementary fluxes are then related to the net flux through the reaction using Equations (31) and (35).

It is important to note that SSF-Evaluator is parallelizable across all mutant networks as reactions fluxes in any particular mutant are independent of metabolite concentrations and enzyme abundances in any other mutant. Based on this, K-SOLVE and SSF-Evaluator generate steady-state reaction fluxes $\boldsymbol{V}^{(SS)}$, relative metabolite concentrations $\boldsymbol{s}^{(SS)}$, and enzyme fractions $\boldsymbol{e}^{(SS)}$ across all mutants given enzyme fractions $\boldsymbol{e}_1$ and reverse elementary fluxes $\boldsymbol{v}_r$ for the WT.

### III. K-UPDATE: Checking for convergence and updating the current WT enzyme fractions and WT reverse elementary fluxes using an implicit Newton step

Procedure K-UPDATE performs an implicit Newton step in the space of of $e_1$ and $v_r$. The updated values for $e_1$ and $v_r$ then yield a new vector of kinetic parameter values $k$ by procedure K-SOLVE. The SSF-Evaluator procedure, in turn, allows for the calculation of the relative metabolite concentrations and enzyme fractions using as input the kinetic parameters estimated by K-SOLVE. This implies that metabolic fluxes $\boldsymbol{V}_c$ in the mutant networks can be expressed as implicit functions of $\boldsymbol{e}_1$ and $\boldsymbol{v}_r$. Executing procedure K-SOLVE and SSF-Evaluator allows for the calculation of the value of these implicit functions $\boldsymbol{V}_c = \boldsymbol{V}_c(\boldsymbol{e}_1, \boldsymbol{v}_r)$. This means that NLP problem P1 can be recast as the following NLP problem with only linear constraints described below. No equality constraints describing conservation of mass across all metabolites and enzyme complexes in the WT need to be explicitly imposed in formulation K-FIT as they are implicitly enforced by K-SOLVE. This limits kinetic parameter values to only those that simultaneously satisfy conservation of mass (for both enzymes and metabolites) and concentration scaling with respect to WT. By propagating the calculated $\boldsymbol{k}$, SSF-Evaluator identifies fluxes $\boldsymbol{V}_c$ across all mutants that automatically satisfy conservation of mass constraints. The objective function $\phi$ as defined in K-FIT below includes the sum of squared errors between metabolic fluxes <u>for only the mutant networks</u>. This is because the measured metabolic fluxes for WT are treated as equality constraints and used for expressing the forward elementary fluxes as a function of the reverse ones (see Equation 47). Note that relative metabolite concentration data for the mutant networks, whenever available, can be appended in the sum of least squares errors objective function in a similar manner.

$$\min_{\boldsymbol{e}_1, \boldsymbol{v}_r} \phi(\boldsymbol{e}_1, \boldsymbol{v}_r) = \sum_{c=2}^{C} \sum_{j \in J_c^{meas}} \left( \frac{V_{jc}(\boldsymbol{e}_1, \boldsymbol{v}_r) - v_{jc}^{(meas)}}{\sigma_{jc}} \right)^2 \qquad \text{(K-FIT)}$$

Subject to:

$$v_{l,1}^{(net)} = V_{j,1} \qquad \forall l \in L_j^{cat} \;\; \forall j \in J \qquad (48)$$

$$v_{l,1}^{(net)} = 0 \qquad \forall l \in L_j^{reg} \;\; \forall j \in J \qquad (49)$$

$$\sum_{l=1}^{n_L} (R_{jl} e_{l,1}) = 1 \qquad \forall j \in J \qquad (40)$$

$$0 \leq e_{l,1} \leq 1 \qquad \forall l \in L \qquad (44)$$

$$v_{r,l} \geq 0 \qquad \forall l \in L \qquad (52)$$

$$v_{l,1}^{(net)} + v_{r,l} \geq 0 \qquad \forall l \in L \qquad (53)$$

Since all constraints in formulation K-FIT are linear, K-FIT can efficiently be solved using a gradient-based method that requires as inputs the first- and second-order gradients of the objective function with respect to the variables $\boldsymbol{e_1}$ and $\boldsymbol{v_r}$ to construct the update formula and check for convergence. However, it is important to note that the nonlinear coupling between $\boldsymbol{e}_1, \boldsymbol{v}_r$ and $V_{jc}$ remains in the objective function as an implicit relation. The expressions that relate the approximate gradient and Hessian to the sensitivity of the predicted steady-state fluxes can be derived by constructing a quadratic approximation for the objective function $\phi$. The following steps describe the construction of the quadratic approximation of $\phi$ used to update $\boldsymbol{e}_1$ and $\boldsymbol{v}_r$ at each iteration of K-FIT.

The variables $\boldsymbol{e}_1$ and $\boldsymbol{v}_r$ are first assembled for convenience into a single $[2n_L \times 1]$ vector $\boldsymbol{x}$

$$\boldsymbol{x} = [(\boldsymbol{e}_1)^{\boldsymbol{T}} | (\boldsymbol{v_r}\,)^{\boldsymbol{T}}]^{\boldsymbol{T}} \tag{73}$$

The objective function $\boldsymbol{\phi}$ and flux through reaction $j$ in mutant $c$ are recast as implicit functions of $\boldsymbol{x}$ as $\phi(\boldsymbol{x})$ and $V_{j,c}(\boldsymbol{x})$**.** The objective function is expressed in vector form as

$$\phi(\boldsymbol{x}) \; = \left(\boldsymbol{V}(\boldsymbol{x}) - \boldsymbol{V}^{(\boldsymbol{meas})}\right)^{T} \boldsymbol{W}^{-\boldsymbol{1}} \left(\boldsymbol{V}(\boldsymbol{x}) - \boldsymbol{V}^{(\boldsymbol{meas})}\right) \tag{74}$$

$\boldsymbol{V}(\boldsymbol{x})$ is the $[n_{meas} \times 1]$ vector of the calculated steady-state fluxes in mutants.

$$n_{meas} = \sum_{c} \left(\text{cardinality } of \ J_c^{(meas)}\right)$$

$\boldsymbol{V}^{(\boldsymbol{meas})}$ is the $[n_{meas} \times 1]$ vector of measured fluxes.

$\boldsymbol{W}$ is the $[n_{meas} \times n_{meas}]$ diagonal matrix storing the variance of the flux measurements, thus

$W_{ii} = \sigma_i^{-2} \; \forall i = \{1,2, \dots, n_{neas}\}$

Upon defining the residual $\boldsymbol{r}(\boldsymbol{x}) = \left(\boldsymbol{V}(\boldsymbol{x}) - \boldsymbol{V}^{(\boldsymbol{meas})}\right)$, the objective function is expressed more compactly as

$$\phi(\boldsymbol{x}) \; = \left(\boldsymbol{r}(\boldsymbol{x})\right)^{T} \boldsymbol{W}^{-\boldsymbol{1}} \boldsymbol{r}(\boldsymbol{x}) \tag{75}$$

For a small perturbation $\Delta\boldsymbol{x}$ to the parameter vector $\boldsymbol{x}$, the objective function at $\boldsymbol{x} + \Delta\boldsymbol{x}$ becomes equal to

$$\phi(\boldsymbol{x} + \Delta\boldsymbol{x}) \; = \left(\boldsymbol{r}(\boldsymbol{x} + \Delta\boldsymbol{x})\right)^{T} \boldsymbol{W}^{-\boldsymbol{1}} \boldsymbol{r}(\boldsymbol{x} + \Delta\boldsymbol{x}) \tag{76}$$

Equation (73) is identical to the least squares representation of isotope tracer-based flux elucidation using 13C-MFA [10]. A popular and successful solution strategy involves constructing a quadratic approximation of the objective function described by Equation (76). Using Taylor series expansion, $\boldsymbol{r}(\boldsymbol{x}+\Delta\boldsymbol{x})$ linearized about $\boldsymbol{x}$ as described by Antoniewicz *et al*. [10] as:

$$r(\boldsymbol{x}+\Delta\boldsymbol{x}) = \boldsymbol{r}(\boldsymbol{x}) + \frac{\partial\boldsymbol{r}}{\partial\boldsymbol{x}}\Delta\boldsymbol{x} \tag{77}$$

$\frac{\partial r}{\partial x}$ is the $[n_{meas} \times 2n_L]$ matrix representing the local sensitivity of $\boldsymbol{r}(\boldsymbol{x})$ with respect to $\boldsymbol{x}$. $\phi(\boldsymbol{x}+\Delta\boldsymbol{x})$ is computed by substituting Equation (77) in Equation (76) yielding:

$$\begin{aligned}\phi(\boldsymbol{x}+\Delta\boldsymbol{x}) &= \left(\boldsymbol{r}(\boldsymbol{x}+\Delta\boldsymbol{x})\right)^T\boldsymbol{W^{-1}}\boldsymbol{r}(\boldsymbol{x}+\Delta\boldsymbol{x}) = \left(\boldsymbol{r}(\boldsymbol{x}) + \frac{\partial\boldsymbol{r}}{\partial\boldsymbol{x}} * \Delta\boldsymbol{x}\right)^T\boldsymbol{W^{-1}}\left(\boldsymbol{r}(\boldsymbol{x}) + \frac{\partial\boldsymbol{r}}{\partial\boldsymbol{x}} * \Delta\boldsymbol{x}\right) \\ &= \left(\boldsymbol{r}(\boldsymbol{x})\right)^T\boldsymbol{W^{-1}}\boldsymbol{r}(\boldsymbol{x}) + \boldsymbol{2}(\Delta\boldsymbol{x})^{\boldsymbol{T}} * \left(\frac{\partial\boldsymbol{r}}{\partial\boldsymbol{x}}\right)^{\boldsymbol{T}}\boldsymbol{W^{-1}}\boldsymbol{r}(\boldsymbol{x}) + (\Delta\boldsymbol{x})^{\boldsymbol{T}}\left(\frac{\partial\boldsymbol{r}}{\partial\boldsymbol{x}}\right)^{\boldsymbol{T}}\boldsymbol{W^{-1}}\frac{\partial\boldsymbol{r}}{\partial\boldsymbol{x}} * \Delta\boldsymbol{x}\end{aligned} \tag{78}$$

The approximate gradient $\boldsymbol{G}$ and the approximate Hessian $\boldsymbol{H}$ are defined using Equation (79).

$$\begin{aligned}\boldsymbol{G} &= \left(\frac{\partial\boldsymbol{r}}{\partial\boldsymbol{x}}\right)^{\boldsymbol{T}}\boldsymbol{W^{-1}}\boldsymbol{r}(\boldsymbol{x}) \\ \boldsymbol{H} &= \left(\frac{\partial\boldsymbol{r}}{\partial\boldsymbol{x}}\right)^{\boldsymbol{T}}\boldsymbol{W^{-1}}\frac{\partial\boldsymbol{r}}{\partial\boldsymbol{x}}\end{aligned} \tag{79}$$

Upon replacing the relevant terms in Equation (78) using the definitions of the objective function $\boldsymbol{\phi}(\boldsymbol{x})$ from Equation (75) and the approximate Gradient and Hessian from Equation (79), Equation (78) is simplified as

$$\phi(\boldsymbol{x}+\Delta\boldsymbol{x}) = \phi(\boldsymbol{x}) + 2\Delta\boldsymbol{x}^T\boldsymbol{G} + \Delta\boldsymbol{x}^T\boldsymbol{H}\Delta\boldsymbol{x} \tag{80}$$

Equation (80) is the local quadratic approximation [10] of the objective function $\boldsymbol{\phi}(\boldsymbol{x})$. In the above expression, $\boldsymbol{G} = \left(\frac{\partial r}{\partial x}\right)^{\boldsymbol{T}}\boldsymbol{W^{-1}}\boldsymbol{r}(\boldsymbol{x})$ and $\boldsymbol{H} = \left(\frac{\partial r}{\partial x}\right)^{\boldsymbol{T}}\boldsymbol{W^{-1}}\frac{\partial r}{\partial x}$ are the approximate gradient and Hessian, respectively. Upon subtracting Equation (75) from Equation (80) we obtain:

$$\Delta\phi = \phi(\boldsymbol{x}+\Delta\boldsymbol{x}) - \phi(\boldsymbol{x}) = 2\Delta\boldsymbol{x}^T\boldsymbol{G} + \Delta\boldsymbol{x}^T\boldsymbol{H}\Delta\boldsymbol{x} \tag{81}$$

A stationary point (i.e., local minimum) for the (approximated) objective function is reached when $\frac{d(\Delta\phi)}{d(\Delta x)} = 0$, which yields:

$$\Delta\boldsymbol{x} = -\boldsymbol{H}^{-1}\boldsymbol{G} \tag{82}$$

Equation (82) computes the unconstrained search direction at each iteration. Note that $\frac{\partial \boldsymbol{r}}{\partial \boldsymbol{x}}$ is needed in to update $\boldsymbol{x}$. Because the residual vector $\boldsymbol{r}(\boldsymbol{x})$ only contains steady-state fluxes, $\frac{\partial r}{\partial \boldsymbol{x}}$ is assembled using the sensitivity of fluxes to $\boldsymbol{x}$ based on the chain rule:

$$\frac{\partial \boldsymbol{v}_c}{\partial \boldsymbol{x}} = \frac{\partial \boldsymbol{v}_c}{\partial \boldsymbol{k}} \frac{\partial \boldsymbol{k}}{\partial \boldsymbol{x}} \tag{83}$$

$\frac{\partial \boldsymbol{v}}{\partial \boldsymbol{k}}$ is computed by differentiating Equation (30) with respect to $\boldsymbol{k}$ to yield:

$$\frac{\partial v_{pc}}{\partial \boldsymbol{k}} = k_p\left(\left(\prod_{\substack{i\\ S_{ip}\le 0}} s_{i,c}^{-S_{ip}}\right)\frac{\partial}{\partial \boldsymbol{k}}\left(\prod_{\substack{l\\ E_{lp}<0}} e_{l',c}\right) + \left(\prod_{\substack{l\\ E_{lp}<0}} e_{l',c}\right)\frac{\partial}{\partial \boldsymbol{k}}\left(\prod_{\substack{i\\ S_{ip}\le 0}} s_{i,c}^{-S_{ip}}\right)\right) + \left(\prod_{\substack{i\\ S_{ip}\le 0}} s_{i,c}^{-S_{ip}}\right)\left(\prod_{\substack{l\\ E_{lp}<0}} e_{l,c}\right)\frac{\partial k_p}{\partial \boldsymbol{k}} \qquad \begin{matrix}\forall p \in P\\ \forall c \in C\\ \forall i \in I\end{matrix} \tag{84}$$

In Equation (84), both $\frac{\partial \boldsymbol{e}_c}{\partial \boldsymbol{k}}$ and $\frac{\partial \boldsymbol{s}_c}{\partial \boldsymbol{k}}$ are unknown. They can be inferred by solving the system of linear algebraic equations formed by differentiating Equations (33), (54), and (56), respectively, with respect to $\boldsymbol{k}$ as follows:

$$\sum_{l=1}^{n_L}\left(R_{jl}\frac{\partial e_{l,c}}{\partial \boldsymbol{k}}\right) = 0 \qquad \begin{matrix}\forall c \in C\\ \forall j \in J\end{matrix} \tag{85}$$

$$\sum_{p=1}^{P} E_{lp}\left(k_p\left(\left(\prod_{\substack{i\\ S_{ip}\le 0}} s_{i,c}^{-S_{ip}}\right)\frac{\partial}{\partial \boldsymbol{k}}\left(\prod_{\substack{l\\ E_{lp}<0}} e_{l,c}\right) + \left(\prod_{\substack{l\\ E_{lp}<0}} e_{l,c}\right)\frac{\partial}{\partial \boldsymbol{k}}\left(\prod_{\substack{i\\ S_{ip}\le 0}} s_{i,c}^{-S_{ip}}\right)\right) + \left(\prod_{\substack{i\\ S_{ip}\le 0}} s_{i,c}^{-S_{ip}}\right)\left(\prod_{\substack{l\\ E_{lp}<0}} e_{l,c}\right)\frac{\partial k_p}{\partial \boldsymbol{k}}\right) = 0 \qquad \begin{matrix}\forall l \in L\\ \forall c \in C\\ \forall i \in I\end{matrix} \tag{86}$$

$$\sum_{p=1}^{P} S_{ip}\left(k_p\left(\left(\prod_{\substack{i\\ S_{ip}\le 0}} s_{i,c}^{-S_{ip}}\right)\frac{\partial}{\partial \boldsymbol{k}}\left(\prod_{\substack{l\\ E_{lp}<0}} e_{l,c}\right) + \left(\prod_{\substack{l\\ E_{lp}<0}} e_{l,c}\right)\frac{\partial}{\partial \boldsymbol{k}}\left(\prod_{\substack{i\\ S_{ip}\le 0}} s_{i,c}^{-S_{ip}}\right)\right) + \left(\prod_{\substack{i\\ S_{ip}\le 0}} s_{i,c}^{-S_{ip}}\right)\left(\prod_{\substack{l\\ E_{lp}<0}} e_{l,c}\right)\frac{\partial k_p}{\partial \boldsymbol{k}}\right) = 0 \qquad \begin{matrix}\forall l \in L\\ \forall c \in C\\ \forall i \in I\end{matrix} \tag{87}$$

The partial derivatives of the product operators $\frac{\partial}{\partial k}\left(\prod_{E_{lp}<0}^{l} e_{l,c}\right)$ and $\frac{\partial}{\partial k}\left(\prod_{S_{ip}\leq 0}^{i} s_{i,c}^{-S_{ip}}\right)$ can be expressed in summation form as shown in Equations (63) and (64). Equations (85), (86), and (87) therefore form a $[(n_L + n_M) \times (n_L + n_M)]$ system of linear algebraic equations that can be solved to obtain $\frac{\partial \boldsymbol{e_c}}{\partial \boldsymbol{k}}$ and $\frac{\partial \boldsymbol{s_c}}{\partial \boldsymbol{k}}$ when $\boldsymbol{k}$, $\boldsymbol{e}_c$, and $\boldsymbol{s}_c$ are specified. $\frac{\partial \boldsymbol{v_c}}{\partial \boldsymbol{k}}$ is calculated by substituting $\frac{\partial \boldsymbol{e_c}}{\partial \boldsymbol{k}}$ and $\frac{\partial \boldsymbol{s_c}}{\partial \boldsymbol{k}}$ into Equation (84). Because $\boldsymbol{x}$ contains both WT enzyme fractions and elementary fluxes, $\frac{\partial \boldsymbol{k}}{\partial \boldsymbol{x}}$ is calculated by differentiating by parts Equation (50) with respect to $\boldsymbol{x}$ to yield:

$$\begin{aligned} \frac{\partial}{\partial \boldsymbol{x}}\left(v_{r,l} + v_{l,1}^{(net)}\right) &= \frac{\partial k_{(2l-1)}}{\partial \boldsymbol{x}}\left(\prod_{\substack{l \\ E_{lp}<0}} e_{l,1}\right) + k_{(2l-1)}\frac{\partial}{\partial \boldsymbol{x}}\left(\prod_{\substack{l \\ E_{lp}<0}} e_{l,1}^{-E_{lp}}\right) \\ \frac{\partial v_{r,l}}{\partial \boldsymbol{x}} &= \frac{\partial k_{(2l)}}{\partial \boldsymbol{x}}\left(\prod_{\substack{l \\ E_{lp}<0}} e_{l,1}\right) + k_{(2l)}\frac{\partial}{\partial \boldsymbol{x}}\left(\prod_{\substack{l \\ E_{lp}<0}} e_{l,1}^{-E_{lp}}\right) \end{aligned} \qquad \begin{matrix} \forall p \in P \\ \forall l \in L \end{matrix} \qquad (88)$$

Solution to the $[n_P \times n_P]$ square system of linear algebraic equations formed by Equation (88) yields $\frac{\partial \boldsymbol{k}}{\partial \boldsymbol{x}}$. Flux sensitivities can be obtained by substituting $\frac{\partial \boldsymbol{k}}{\partial \boldsymbol{x}}$ in Equation (84). Having computed the sensitivity of elementary fluxes, the sensitivity of all net reaction fluxes is calculated by substituting $\frac{\partial \boldsymbol{v}}{\partial \boldsymbol{x}}$ in Equations (89) and (90), which are obtained by differentiating Equations (31) and (35) with respect to $\boldsymbol{x}$ as shown below.

$$\frac{\partial v_{l,c}^{(net)}}{\partial \boldsymbol{x}} = \frac{\partial v_{(2l-1),c}}{\partial \boldsymbol{x}} - \frac{\partial v_{2l,c}}{\partial \boldsymbol{x}} \qquad \begin{matrix} \forall l \in L \\ \forall c \in C \end{matrix} \qquad (89)$$

$$\frac{\partial V_{j,c}}{\partial \boldsymbol{x}} = \sum_{l=1}^{n_L} (N_{jl}\frac{\partial v_{l,c}^{(net)}}{\partial \boldsymbol{x}}) \qquad \begin{matrix} \forall j \in J \\ \forall c \in C \end{matrix} \qquad (90)$$

The sequence of steps to be followed to compute the approximate gradients of the objective function and update the variables $\boldsymbol{x}$ in every iteration is described by the algorithmic procedure for K-UPDATE.

**Algorithmic procedure K-UPDATE**

**begin**

Specify and fix $\boldsymbol{k}$, $e$, $\boldsymbol{s}$, and $\boldsymbol{V}$ computed by K-SOLVE and SSF-Evaluator

Specify measured fluxes $\boldsymbol{V}^{(meas)}$ and the weighting matrix $\boldsymbol{W}$

Specify list of $mutants$

Compute $\frac{\partial \boldsymbol{k}}{\partial \boldsymbol{x}}$ by solving the $[n_P \times n_P]$ system of linear Equation (88) using $\boldsymbol{e}_1 \coloneqq \boldsymbol{e}_1^{(SS)}$ and $\boldsymbol{k} \coloneqq \boldsymbol{k}$

**for all mutants:**

Calculate sensitivities $\frac{\partial \boldsymbol{s}_c^{(SS)}}{\partial \boldsymbol{k}}$ and $\frac{\partial \boldsymbol{e}_c^{(SS)}}{\partial \boldsymbol{k}}$ by solving the $[(n_L + n_M) \times (n_L + n_M)]$ system of linear Equations (85), (86), and (87) using $\boldsymbol{e}_c \coloneqq \boldsymbol{e}_c^{(SS)}$ and $\boldsymbol{s}_c \coloneqq \boldsymbol{s}_c^{(SS)}$

Calculate $\frac{\partial \boldsymbol{v}_c^{(SS)}}{\partial \boldsymbol{k}}$ by substituting $\frac{\partial \boldsymbol{s}_c^{(SS)}}{\partial \boldsymbol{k}}$ and $\frac{\partial \boldsymbol{e}_c^{(SS)}}{\partial \boldsymbol{k}}$ in Equation (84).

Calculate $\frac{\partial \boldsymbol{v}_c^{(SS)}}{\partial \boldsymbol{x}}$ by substituting $\frac{\partial \boldsymbol{v}_c^{(SS)}}{\partial \boldsymbol{k}}$ and $\frac{\partial \boldsymbol{k}}{\partial \boldsymbol{x}}$ into Equation (83).

Calculate $\frac{\partial \boldsymbol{v}_c^{(net)}}{\partial \boldsymbol{x}}$ by substituting $\frac{\partial \boldsymbol{v}_c^{(SS)}}{\partial \boldsymbol{x}}$ into Equation (89).

Calculate $\frac{\partial \boldsymbol{V}_c}{\partial \boldsymbol{x}}$ by substituting $\frac{\partial \boldsymbol{v}_c^{(net)}}{\partial \boldsymbol{x}}$ into Equation (90).

Assemble the residual vector $\boldsymbol{r}(\boldsymbol{x})$ from $\boldsymbol{V}$ and $\boldsymbol{V}^{(meas)}$

Assemble the sensitivity matrix $\frac{\partial \boldsymbol{r}}{\partial \boldsymbol{x}}$ from $\frac{\partial \boldsymbol{V}}{\partial \boldsymbol{x}}$

Compute the objective function $\phi$ by substituting $\boldsymbol{r}(\boldsymbol{x})$ and $\boldsymbol{W}$ into Equation (75).

Compute the approximate gradient $\boldsymbol{G}$ and the approximate Hessian $H$ by substituting $\boldsymbol{r}(\boldsymbol{x})$, $\frac{\partial \boldsymbol{r}}{\partial \boldsymbol{x}}$, and $\boldsymbol{W}$ into Equation (79)

**return** $\phi$, $\boldsymbol{G}$, and $\boldsymbol{H}$

**end**

### Algorithmic description of K-FIT

Procedures K-SOLVE, SSF-Evaluator, and K-UPDATE are integrated into the algorithm K-FIT as described below. Briefly, WT enzyme fractions $\boldsymbol{e}_1$ and reverse elementary fluxes $\boldsymbol{v}_r$ satisfying (in)equalities in Equations (40), (44), (52), and (53) are randomly initialized. For the sake of brevity, we combine the action of K-SOLVE and SSF-Evaluator into a single procedure FLUX-SOLVE within K-FIT which calculates steady-state fluxes for the mutant networks given $\boldsymbol{e}_1$ and $\boldsymbol{v}_r$. In the first step of FLUX-SOLVE, kinetic parameters $\boldsymbol{k}$ anchored to WT steady-state fluxes $\boldsymbol{V}_1$ are computed from $\boldsymbol{e}_1$ and $\boldsymbol{v}_r$ using procedure K-SOLVE. The kinetic parameters are then used to evaluate the steady-state fluxes in the mutant networks using procedure SSF-Evaluator. Having computed steady-state fluxes in mutants $\boldsymbol{V}$, relative metabolite concentrations $\boldsymbol{s}$, and enzyme fractions $\boldsymbol{e}$, the objective function $\phi$ and its approximate gradient $\boldsymbol{G}$ and Hessian $\boldsymbol{H}$ are computed using procedure K-UPDATE. $\boldsymbol{G}$ and $\boldsymbol{H}$ are used to check for convergence and update $\boldsymbol{e}_1$ and $\boldsymbol{v}_r$ if optimality is not achieved.

The algorithmic description of FLUX-SOLVE is provided below:

**Algorithmic description of FLUX-SOLVE**

**begin**

> Specify and fix WT enzyme fractions $\boldsymbol{e}_1$ and reverse elementary fluxes $\boldsymbol{v}_r$.
>
> Specify and fix the WT steady-state flux distribution $\boldsymbol{V}_1$.
>
> Specify the list of $mutants$
>
> Compute anchored kinetic parameters $\boldsymbol{k}$ using Procedure **K-SOLVE** with the specified $\boldsymbol{e}_1$, $\boldsymbol{v}_r$, and $\boldsymbol{V}_1$.
>
> **for all *mutants***
>
> > Compute steady-state fluxes $\boldsymbol{V}_c^{(SS)}$, relative metabolite concentrations $\boldsymbol{s}_c^{(SS)}$, and enzyme fractions $\boldsymbol{e}_c^{(SS)}$ in $mutant\ c \in C$ using **SSF-Evaluator** with kinetic parameters $\boldsymbol{k}$
> >
> > Set $\boldsymbol{V}_c \coloneqq \boldsymbol{V}_c^{(SS)}$, $\boldsymbol{s}_c \coloneqq \boldsymbol{s}_c^{(SS)}$, and $\boldsymbol{e}_c \coloneqq \boldsymbol{e}_c^{(SS)}$
>
> **return** $\boldsymbol{V}$, $\boldsymbol{s}$, and $\boldsymbol{e}$

**end**

The overall workflow for the K-FIT algorithm combining procedures FLUX-SOLVE and K-UPDATE is described below and is also pictorially shown in Supplementary Figure S1:

**Overall algorithmic procedure K-FIT**

**begin**

> Specify and fix WT flux distribution $\boldsymbol{V}_1$, measured fluxes $\boldsymbol{V}^{(meas)}$, variance $\boldsymbol{W}$, set of mutants, $xtol$ and $gtol$
>
> Randomly initialize $\boldsymbol{x}$ satisfying constraints in Equations (40), (44), (52), and (53).
>
> Set $stol \coloneqq 10^{-6}$
>
> Using FLUXSOLVE and inputs $\boldsymbol{x}$ evaluate initial steady-state fluxes $\boldsymbol{V}$, relative metabolite concentrations $\boldsymbol{s}$, and enzyme fractions $\boldsymbol{e}$.
>
> Evaluate the initial value of the objective function $\phi(\boldsymbol{x})$ and gradients $\boldsymbol{G}$ and $\boldsymbol{H}$ using K-UPDATE.
>
> Set $done \coloneqq false$
>
> Set $\boldsymbol{x}_{\boldsymbol{best}} \coloneqq \boldsymbol{x}$, $\phi_{best} \coloneqq \phi(\boldsymbol{x})$
>
> **while** (not $done$)
>
> > Compute $\Delta\boldsymbol{x}$ using Equation (82)

**if** $(\|\Delta \boldsymbol{x}\|_\infty \leq xtol)$ **or** $(\|\boldsymbol{G}\|_\infty \leq gtol)$

Set $done \coloneqq true$

**else**

Update $\boldsymbol{x} \coloneqq \boldsymbol{x_{best}} + \Delta \boldsymbol{x}$

Using FLUX-SOLVE and inputs $\boldsymbol{x}$ evaluate steady-state fluxes $\boldsymbol{V}$, relative metabolite concentrations $\boldsymbol{s}$, and enzyme fractions $\boldsymbol{e}$.

Evaluate the initial value of the objective function $\phi(\boldsymbol{x})$ and gradients $\boldsymbol{G}$ and $\boldsymbol{H}$ using K-UPDATE.

**if** $\phi(\boldsymbol{x}) < \phi_{best}$

Update $\boldsymbol{x_{best}} \coloneqq \boldsymbol{x}$, $\phi_{best} \coloneqq \phi(\boldsymbol{x})$

**return** $\boldsymbol{x}_{best}, \phi_{best}$

**end**

Supplementary Figure S1: Overview of the K-FIT algorithm showing the flow of information between various components.

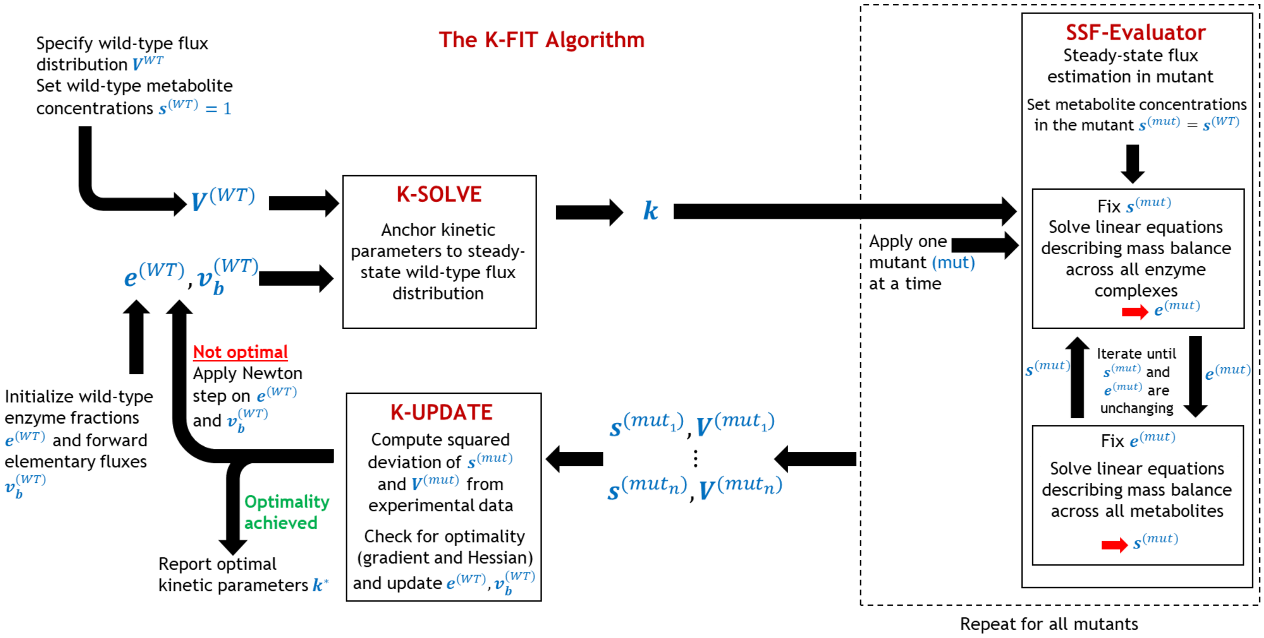
